# Nuclear activation in dual-durotaxing cells on a matrix with cell-scale stiffness-heterogeneity

**DOI:** 10.1101/2021.10.30.466554

**Authors:** Satoru Kidoaki, Hiroyuki Ebata, Kosuke Moriyama, Thasaneeya Kuboki, Yukie Tsuji, Rumi Sawada, Saori Sasaki, Tatsuya Okuda, Kosuke Hamano, Takahito Kawano, Aki Yamamoto, Ken Kono, Kazusa Tanaka

## Abstract

Living organisms are typically composed of various tissues with microscopic cell-scale stiffness-heterogeneity, in which some cells receive dynamically fluctuating mechanical stimuli from the heterogeneous extracellular milieu during long-term movement. Although intracellular stress dynamics (ISD), which are closely related to the regulation of cell functions such as proliferation and differentiation, can be characteristically modulated in cells migrating on a matrix with stiffness-heterogeneity, it has been unclear how the mode of fluctuation of ISD affects cell functions. In the present study, we demonstrate that mesenchymal stem cells (MSCs) dual-durotaxing (i.e., both forward and reverse durotaxis) on microelastically-patterned gels with stiff triangular domains markedly amplify the fluctuation of ISD, nuclear shape, and the spatial distribution of chromatins, which makes the cells remain far from tensional equilibrium. We provide evidence that amplified chromatin fluctuation in the dual-durotaxing MSCs can cause activation of cellular vigor and maintenance of the stemness.

## Main text

In living organisms, cells reside in the extracellular matrix (ECM) of tissues with intrinsic mechanical properties that are often micromechanically heterogeneous across a spatial order of several tens to hundreds of microns^1–3^. Some types of cells migrate on a matrix with cell-scale stiffness-heterogeneity, such as is typically seen in the process of wound healing in disordered tissues^4,5^, biological development in immature growing tissues^6–8^, and long-distance stem cell migration^9–11^. Migrating cells dynamically remodel the adhesion machinery and cytoskeletal architecture through interaction with the mechanical milieu^12,13^, and thus exhibit continuous deformation in cell shape and spatiotemporal fluctuation in intracellular prestress^14^.

The cell shape has been revealed to modulate cell functions such as proliferation and differentiation by altering the intracellular prestress^15,16^ generated by mechanotransductive linkages between the ECM, focal adhesions, cytoskeletons, and the cell nucleus^17^. Such a mechanical network influences chromatin behavior and finally regulates gene expression^18–21^. For example, specific positioning of chromatins and intermingling of different chromatin territories play a crucial role in cell-shape-dependent gene expression^22^. Dynamic cell-shaping during cell migration should induce, in principle, characteristic intracellular stress dynamics (ISD) and the resulting modulation in cell functions.

How does ISD behave on a matrix with stiffness-heterogeneity? We previously clarified that the fluctuation of ISD in a single moving cell is markedly amplified on the matrix over a time scale of hours^23^. In such a model system of a heterogeneous mechanical milieu in vivo, how does the amplified fluctuation of ISD affect cell functions? In this study, to reveal the nature of functional regulation in migrating cells, we demonstrate the essential role played by the amplified fluctuation of ISD and nuclear deformation in the functional activation of migrating mesenchymal stem cells (MSCs).

Our strategy was to make the cells nomadically migrate among different regions of stiffness by alternatively driving both forward and reverse durotaxis over stiff triangular domains^22^ (Fig. 1a). In general, adherent cells durotax and accumulate in the stiff region of a matrix^24,25^. In this sense, durotaxis is a transient phenomenon before migrating cells eventually settle into tensional equilibrium. However, if we introduce cues for both forward and reverse durotaxis^26^ to the matrix, this settling can be avoided, i.e., dual-durotaxing cells should exhibit non-biased random crawling even on a heterogeneous field of stiffness and remain far from tensional equilibrium. This kind of combined-taxis-inducing system, which exists as mixotaxis in vivo^27^, can be a valuable model for investigating dynamic cell behaviors in ECM with mechanical heterogeneity. For instance, bone-marrow-derived MSCs have been reported to delocalize in a mechanically heterogeneous matrix^28^ composed of trabecula (osteoid; 25-40 kPa^29^, precalcified bone; ∼100 kPa^30^), sinusoid (vessel; 2-10 kPa^31,32^), and cellular interstitium (∼0.3 kPa^33^).

**Figure 1.**
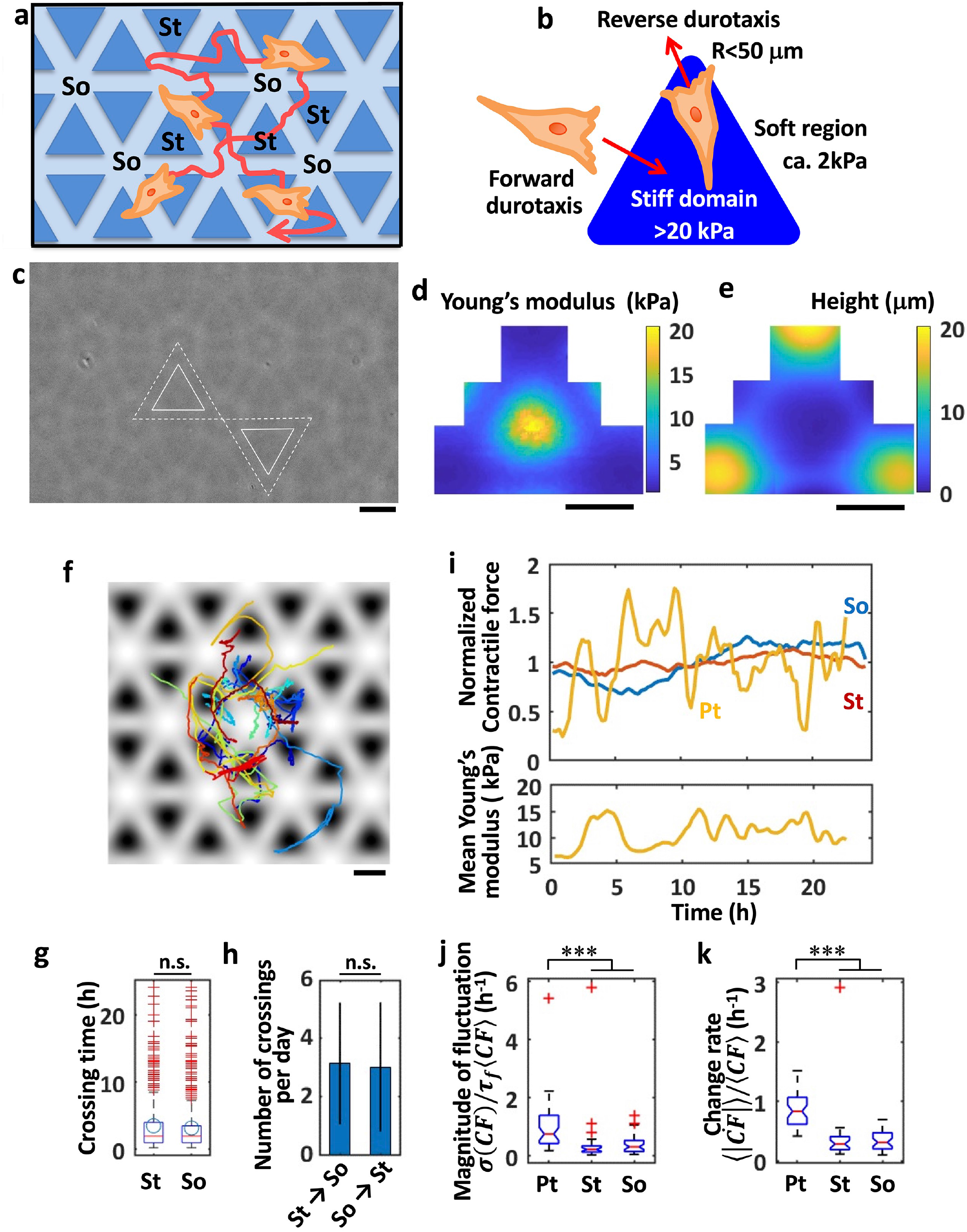
Nomadic cell migration on stiff-triangular patterned gels amplifies ISD fluctuation. **(a)** Schematic representation of nomadic cell migration on regions of different stiffness. **(b)** A stiff triangular pattern to induce both forward and reverse durotaxis. **(c)** Phase-contrast microscopic images of patterned (Pt) gels with stiff triangular domains. Side length: 150 μm, inner vertex distance: 120 μm. White solid and dashed triangles exemplify images for the edge of a domain and a unit pattern. **(d, e)** Distribution of Young’s modulus (d) and height (e) around the unit domain. **(f)** Trajectories of the centroid of the MSCs migrating on Pt gels. Background gray-scaled color: elasticity distribution estimated from (d) (white: soft, black: stiff). n = 20. **(g, h)** Crossing time (g) and the number of crossings per day (h) for MSCs to move across the elasticity boundary between stiff and soft regions. Bars denote mean and standard deviation for the measured cells. n = 223. **(i)** Representative dynamics of contractile force in a single cell moving on Pt (Yellow), stiff (St; red), and soft (So; blue) gels. Contractile force *CF* is normalized by the time-averaged value <*CF*>. The detailed analysis is described in Fig. S1. **(j, k)** The magnitude of fluctuation (j; the ratio of the normalized standard deviation of *CF* to correlation time) and the rate of change in the dynamics of normalized contractile forces (k; time derivative of the fluctuation) calculated for Pt, St, and So gels. n = 46 (Pt), 28 (St), 39 (So). ***: P<0.001. Scale bars: 100 μm.

By using a microelastically-patterned matrix that can induce the perfect nomadic migration of MSCs, we provide evidence that the amplified fluctuation of ISD and nuclear deformation in dual-durotaxing MSCs causes the functional activation of various genes related to cellular vigor as well as the maintenance of stemness.

### Dual-durotaxing-induced amplification of fluctuation of ISD

The design principle for microelastically-patterned hydrogels to induce dual-durotaxis is to introduce stiff triangular regions, the size of which is comparable to single cells and the corners of which have a curved elasticity boundary that is convex toward the soft region (Fig. 1b). We previously clarified that reverse durotaxis from the stiff to the soft region could be induced specifically at the convex boundary with a radius of curvature smaller than 50 μm, while usual forward durotaxis occurs at the side of the triangular domain^26^. First, long-term ISD behaviors (∼24hrs) of dual-durotaxing cells were analyzed using this sophisticated platform.

Stiff triangular domains (elastic modulus: 20–40 kPa) were patterned on the soft base matrix (2–4 kPa) by photolithographic patterning using photo-crosslinkable gelatins^23,34–37^ (Fig. 1c and d). The surface topography of the matrix was carefully set to be smooth by intentionally using out-of-focus irradiation, as seen in the fuzzy phase contrast photograph in Fig. 1c. The height difference in the elasticity boundary region was below 20 μm and was prepared so that natural cell movement was not hindered (Fig. 1e). The triangle’s side length (ca. 150 μm) and apex distance (ca. 120 μm) were optimized to balance the residence time between the stiff and soft regions. The observed trajectories of each cell confirmed that the cells migrate in and out of stiff and soft regions (Fig. 1f, SI Movie 1). Individual cells exhibited perfect nomadic migration between stiff and soft regions approximately every 4 hours (Fig. 1g), and crossed between these regions about 5 times every 24 hours (Fig. 1h).

By using finite element method-based traction force microscopy on a matrix with stiffness-heterogeneity^23^, we measured the magnitude of fluctuation of contractile force *CF* as the normalized standard deviation of the long-term contractile force dynamics (∼24 hrs.) (Fig. 1i and SI Movies 2-4). A significant increase in the magnitude of fluctuation and the rate of change of the normalized contractile force dynamics on the patterned (Pt) gels evidenced that the fluctuation of ISD was markedly amplified in dual-durotaxing cells compared with cells randomly crawling on the control plain stiff (St: 20–40 kPa) and soft (So: 2–4 kPa) gels (Fig. 1j, k, and Fig. S1a) as was the fluctuation of cell shape (Fig. S1b and c), suggesting that the dual-durotaxing cells are forced to remain far from tensional equilibrium.

### Amplified fluctuation of nuclear shape and spatial distribution of chromatin in dual-durotaxing MSCs

ISD are mainly derived from the structural dynamics of the cytoskeletal network that is linked to the cell nucleus through the Linker of Nucleoskeleton and Cytoskeleton (LINC) complex^18,38^ on the nuclear membrane. The amplified fluctuation of ISD in dual-durotaxing MSCs should transmit enhanced mechanical stimuli from the cytoskeleton to the nucleus. To confirm this effect, we characterized the fluctuation of the shape of the cell nucleus by measuring the dynamics of the aspect ratio through live-cell imaging over 12 hrs (Fig. 2a, SI Movies 5,6, Fig. S2). The magnitude of fluctuation of the aspect ratio was defined as the standard deviation of the ratio in the time-course data, which was normalized with the means of both the fluctuation and correlation time of the autocorrelation function. On Pt gels, the magnitude of fluctuation and the deformation rate of the nuclear shape were significantly larger than those on the control culture plate (CP) (Fig. 2b and c, respectively; the comparison of St and So is described in Fig. S2). This amplified nuclear fluctuation in dual-durotaxing MSCs on Pt gels was diminished by the tension inhibitors Y27632, blebbistatin, and cytochalasin D, as seen in the reduced deformation rate of the nuclear aspect ratio (Fig. 2d), which constituted evidence that tensional fluctuation in the actomyosin cytoskeletal network plays a critical role in force transmission to the nucleus.

**Figure 2.**
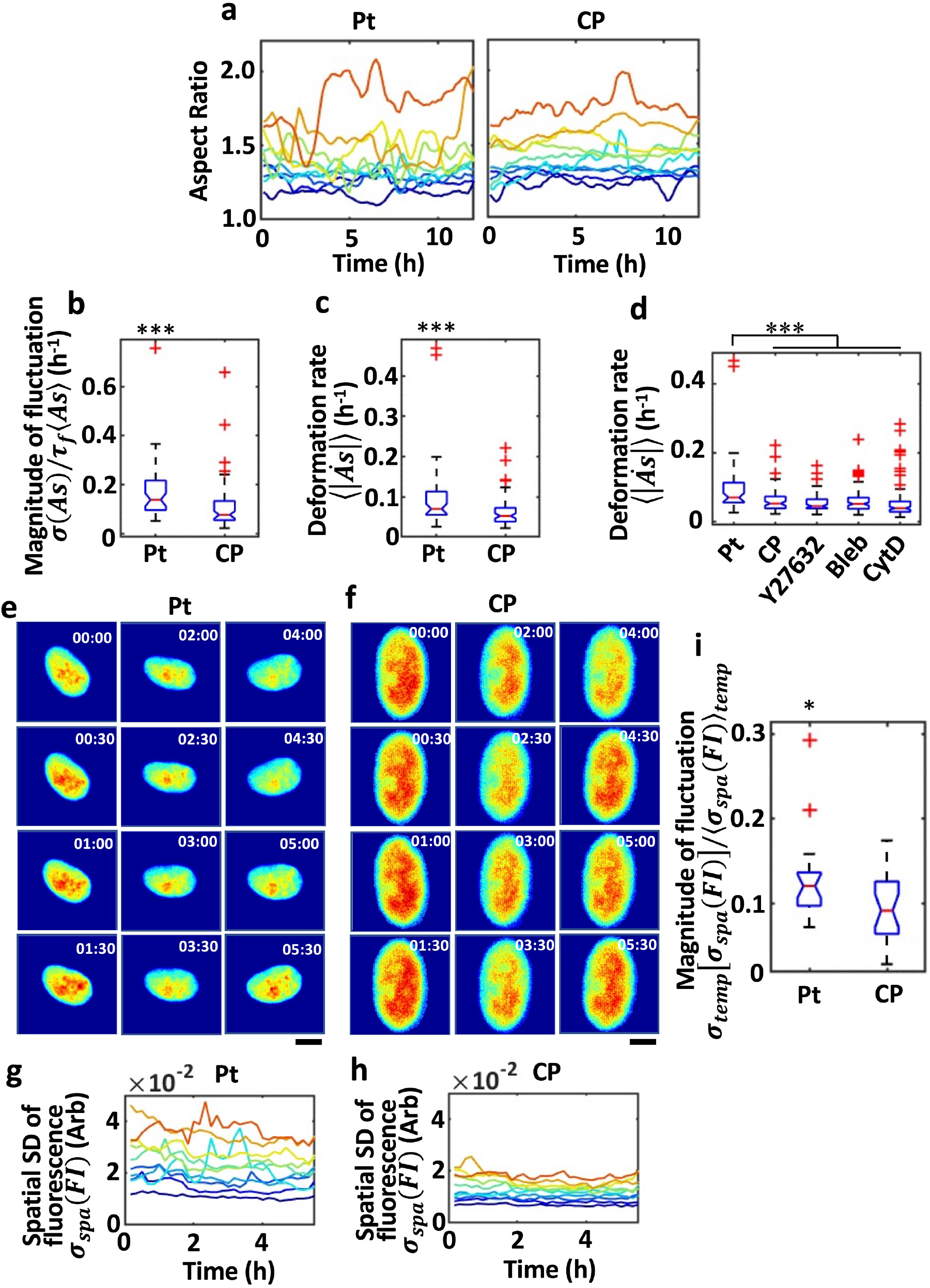
Amplified fluctuation of nucleus shape and spatiotemporal distribution of chromatins in dual-durotaxing MSCs. **(a)** Representative long-term fluctuation of the nuclear shape in cells migrating on Pt gels and CP. **(b)** Comparison of the magnitude of fluctuation of nuclear shape between Pt gels and CP, which was defined as the ratio of the normalized standard deviation to correlation time in the fluctuation of the aspect ratio of the nucleus. n = 51 (Pt), 71 (CP). The detailed analysis is described in Fig. S2. **(c)** The deformation rate of nuclear shape in cells migrating on Pt gels and CP is defined as the time derivative of the dynamics of the nuclear aspect ratio. n = 51 (Pt), 71 (CP). **(d)** Effects of actomyosin inhibitors on the fluctuation of nuclear shape: deformation rate was defined as the time derivative of the dynamics of the nuclear aspect ratio. n = 51 (Pt), 71 (CP), 70 (Y27632), 131 (Blebbistatin), and 124 (Cytochalasin D) **(e, f)** Representative time-lapse images of the chromatin distribution in cells migrating on Pt gels (e) and CP (f). Time interval: 30 min. The color scale shows the fluorescence intensity (*FI*). Scale bars: 10 μm. **(g, h)** Time-course plot of the spatial standard deviations *σ_spa_* (*FI*) for the chromatin distribution in cells migrating on Pt gels (g) and CP (h). **(i)** Comparison of the magnitude of spatiotemporal fluctuation in the chromatin distribution in cells cultured on Pt gels and CP, defined as the temporal standard deviation of *σ_spa_* (*FI*) normalized by the temporal mean of chromatin fluctuation *σ_temp_* [*σ_spa_*(*FI*)] /.〈*σ_spa_*(*FI*)〉 *_temp_*. n= 20 (Pt), 25 (CP). ***: P<0.001. *: P<0.05.

How does the amplified fluctuation in nuclear shaping affect the dynamic behavior of chromatins? Next, we analyzed the spatiotemporal fluctuation of fluorescently labeled chromatins through live imaging over 5.5 hrs for single nuclei in dual-durotactic cells (Fig. 2e and g, SI Movie 7) on Pt gels and usual crawling cells on CP (Fig.2f and h, SI Movie 8). The magnitude of fluctuation of the spatial distribution of chromatins was markedly enhanced on Pt (Fig. 2i). These results are evidence that dual-durotaxing MSCs with amplified fluctuation of ISD have jittered chromatins inside the greatly fluctuating nucleus.

### Stemness of dual-durotaxing MSCs and nucleocytoplasmic shuttling of YAP

How is the stemness affected in MSCs for which ISD and nuclear shaping are amplified during dual-durotactic migration? There was no change in any of the MSC markers (positive markers: CD73, CD105, CD90, and negative markers: CD11b, CD14, CD19, CD34, CD45, HLA-DR) between before and after dual-durotactic culture (Fig. 3a and b). The differentiation potential of MSCs that had been subjected to 1 week of mechanical stimulation from CP, St, So, and Pt gels was characterized directly on each sample (i.e., in situ differentiation induction; see scheme in Fig. 3c-(I)) and on CP after reseeding from the gel samples (i.e., post-Pt standard differentiation induction; see scheme in Fig. 3c-(II)). As for osteogenic in situ induction (Fig. 3d-(I)), dual-durotactic MSCs showed suppressed differentiation (Fig. 3f-(I)) at a level similar to that of cells on So gels, suggesting that the cells either could resist the differentiation bias toward osteogenic lineages or had lost their differentiation potential. To explore these possibilities, standard induction on CP after Pt culture was performed (Fig. 3d-(II)). Osteogenic differentiation was fully recovered on Pt gels compared with on CP (Fig. 3f-(II)), which confirmed that dual-durotaxing MSCs retained their stemness and osteogenic differentiation potential. In addition, there was no accumulation of mechanical dose that would make cells biased for osteogenic lineage and deteriorate their stemness. On the other hand, adipogenic in situ induction for dual-durotaxing MSCs exhibited highly efficient differentiation compared with the control CP (Fig.3e-(I) and g-(I)). After culture on Pt gels, the reseeded cells showed that they had maintained a highly efficient adipogenic differentiation in post-Pt standard induction (Fig.3e-(II) and g-(II)). These results confirmed that dual-durotaxing MSCs maintained their stemness, which significantly suppresses the bias toward an osteogenic lineage.

**Figure 3.**
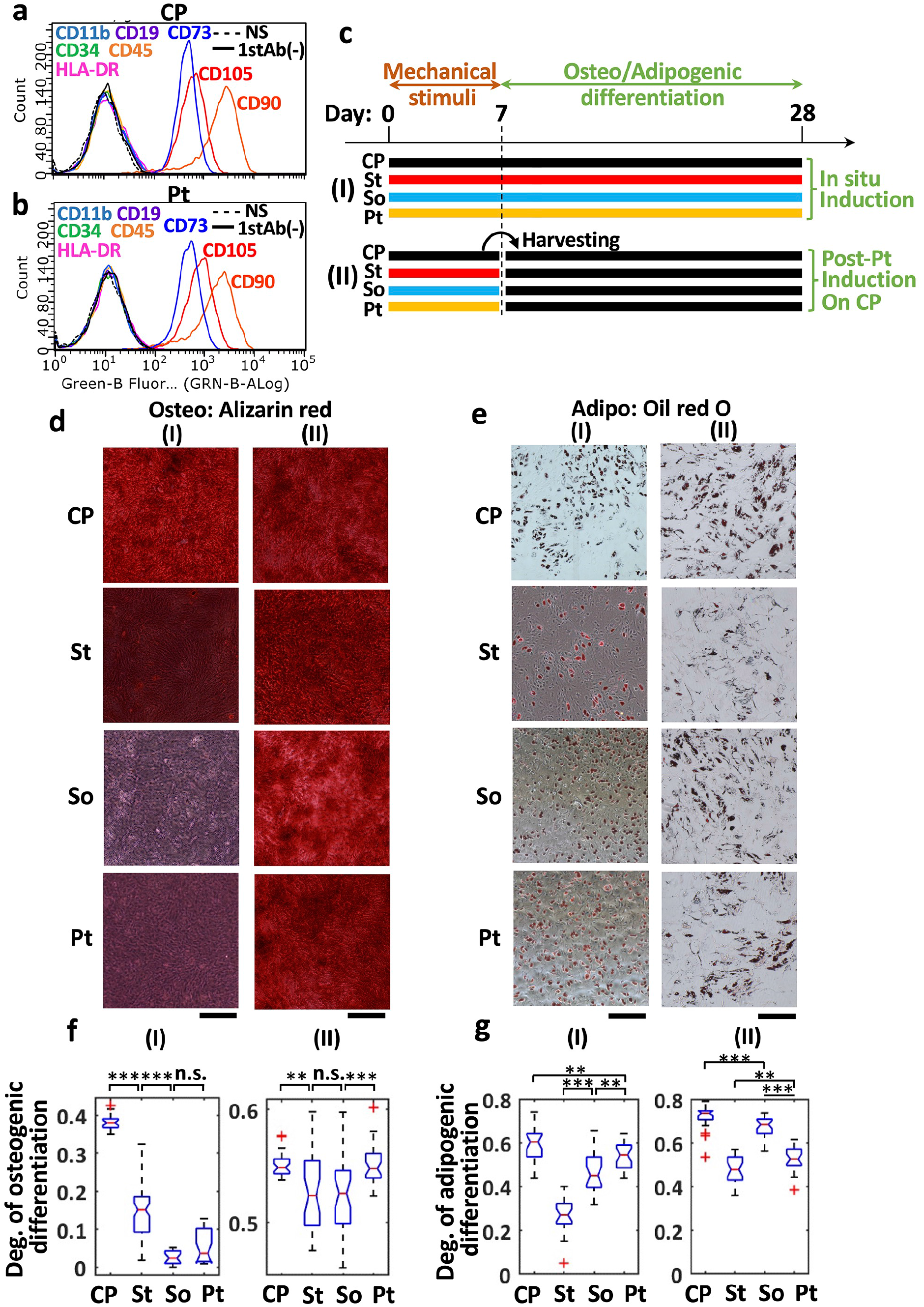
Characterization of stemness of dual-durotaxing MSCs. **(a, b)** Flow cytometric analysis of MSCs cultured for 1week on control CP (a) and Pt gels (b). Positive markers: CD73, CD105, CD90; Negative markers: CD11b, CD14, CD19, CD34, CD45, HLA-DR. N.S: Not stained control, 1st Ab (-): without primary antibody. **(c)** Scheme of the differentiation-induction experiment. Cells were subjected to mechanical stimuli from the culture substrates of CP, St, So, and Pt for one week. Next, osteogenic or adipogenic differentiation was induced for 3 weeks with the cells in situ on each substrate (I) or on the CP after harvesting and reseeding from the substrates (II). **(d, e)** Alizarin red staining and Oil red O staining of MSCs on CP, St, So, and Pt substrates after the experiment c-(I) and c-(II). **(f, g)** Quantification of the degree of osteogenic (f) or adipogenic (g) differentiation: In situ induction (I) and post-gel induction (II) after cells were cultured on CP, St, So, and Pt substrates. N (number of samples); n (number of frames analyzed), and A (frame area analyzed): (f)-(I) CP: 3; 27, St: 8; 72, So: 2; 18, Pt: 3; 27, and 0.95 ± 0.26 mm^2^. (f)-(II) CP: 4; 36, St: 4; 36, So: 4; 36, Pt: 4; 36, and 4.9 ± 1.4 mm^2^. (g)-(I) CP: 3; 27, St: 3; 26, So: 3; 27, Pt: 3; 27, and 14 ± 6.0 mm^2^. (g)-(II) CP: 3; 27, St: 3; 27, So: 3; 27, Pt: 3; 27, and 6.8 ± 2.0 mm^2^. ***: P<0.001. **: P<0.01. Scale bars: 500 μm.

To address the mechanism of this in situ suppression of an osteogenic bias, we characterized the intracellular localization of YAP and RUNX2 (Fig. S3). These were mainly localized in the cell nucleus and cytoplasm on plain St and So gels, respectively. However, on Pt gels they were delocalized inside and outside the nucleus between stiff and soft regions. The distribution measured from the fluorescence ratio indicated the typical trend for stiffness-dependent localization of YAP and RUNX2 on homogeneous gels and delocalization on Pt gels (Fig.S3). This observation carries a corollary that the accumulation of mechanical dose toward an osteogenic lineage should be inhibited through the nucleocytoplasmic shuttling of mechanotransducer during nomadic migration between regions of different stiffness. Under these conditions, the specific differentiation-related signaling pathway should always be perturbed due to the short-term switching based on different levels of mechanical stimuli; thus, no pathway could lead to a specific lineage. Live-cell imaging showed YAP shuttling, which occurred with a time lag of about 30 min after the cell crossed the elastic boundary (SI Movie 9). Indeed, the nomadic migration of MSCs among regions of different stiffness made stiffness-dependent mechanotransducer move nomadically inside cells and contributed to the failure to accumulate the signals required for lineage specification, which we previously referred to as “frustrated differentiation”^39,40^.

### Enhanced gene expression and cytokine production in dual-durotaxing MSCs with amplified nuclear fluctuation

What impact does the amplified fluctuation of chromatins have on the modulation of gene expression and function in dual-durotaxing MSCs with maintained stemness? After four days of culture, a comprehensive gene expression analysis was performed for MSCs on Pt gels and the control substrates (St, So, CP). The numbers of genes that were up-regulated (>1.5-fold) and down-regulated (<0.67-fold) relative to those in cells on CP are shown in Fig. S4. There were 193 and 262 Pt-specific up-regulated and down-regulated genes, respectively. Pt-, St-, and So-specifically regulated genes are listed in Tables S1-S3. By using Ingenuity® Pathway Analysis (IPA), we found canonical signaling pathways related to the Pt-specific regulated genes, as shown in Fig. S5. While signaling related to IL-8, ILK, FGF, PDGF, HGF, Netrin, Mouse ESC pluripotency, and so on, was shown to have high positive Z-scores above 2.0, only a few pathways showed repressed signaling with high negative Z-scores (Insulin secretion and FGF signaling). Interestingly, a comparison with canonical pathway analysis revealed that only Pt-culture was associated with various signaling activation with statistical cut-off criteria of -Log P > 1.3 and Z-score > 2.0. No such predominant response was observed in St- or So-cultures. The locations of the regulated genes in each pathway are shown in Fig. S6.

Another IPA biofunction analysis, which evaluates the degree of significance in knowledge-database-based prediction for functional activation or inactivation in cell function, also suggested a clear tendency for the activation of various functional categories only for the dual-durotaxing cells (Fig. 4a, Fig. S7). Based on these results, a comparison between Pt-, St-, and So-culture with cut-off criteria of -log P>1.3 and Z score > 2.0 in physiological functional categories showed very clearly that the substantial inactivation of morbidity or mortality, organismal death, apoptosis, and necrosis was predicted only on Pt. In contrast, the significant activation of vigor-related functional categories, including cell survival, cell viability, cell movement, and cell migration, was predicted, along with the activation of other various cellular functions (Fig. 4b). The groups of genes associated with these predictions are summarized in Fig. S8.

**Figure 4.**
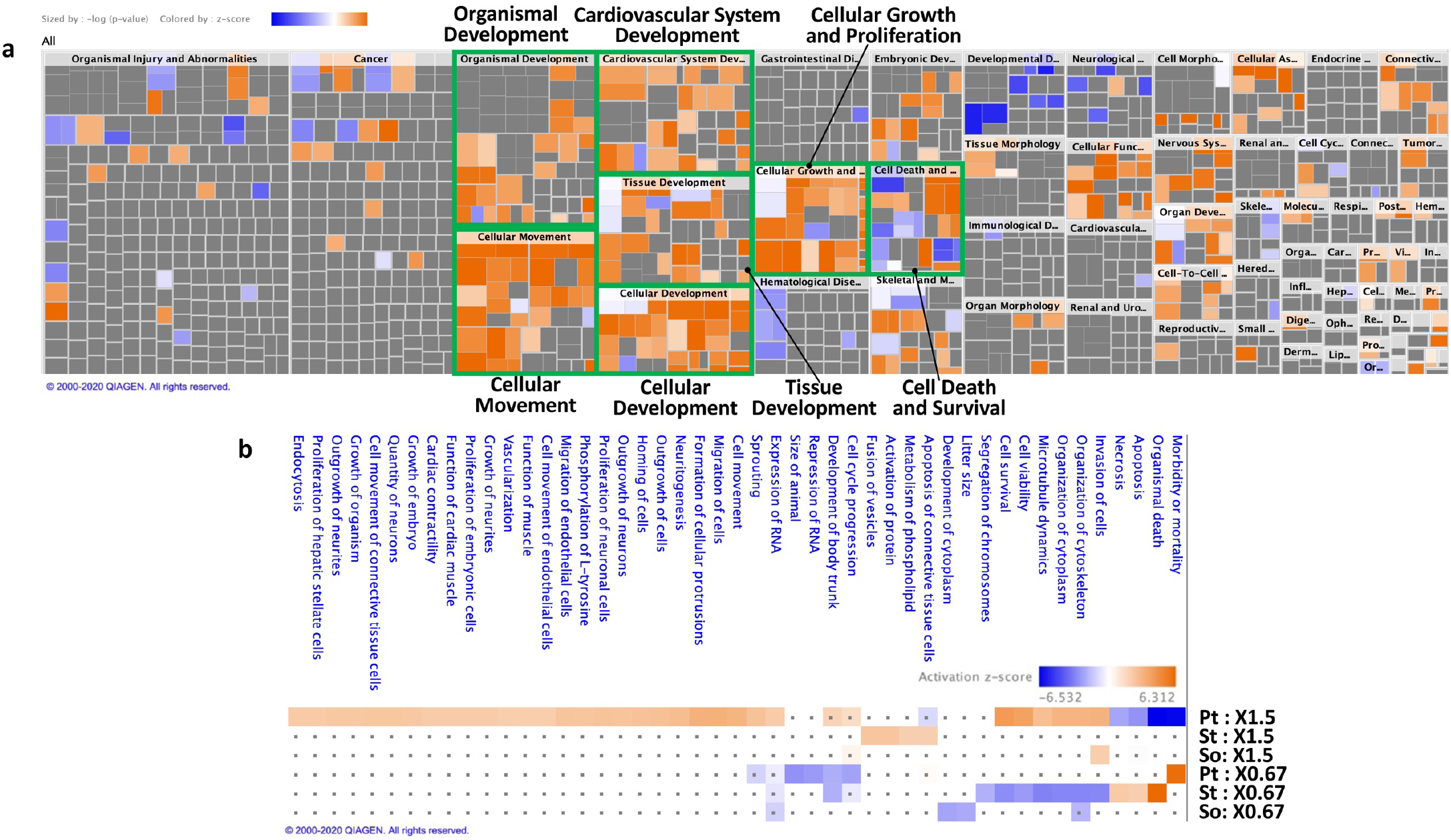
Analysis of the IPA downstream effects in dual-durotaxing MSCs. **(a)** Activation and inactivation of biological functions predicted for dual-durotaxing cells on Pt gels. Orange and blue squares predict an increase and decrease, respectively, in the related functions, colored according to the activation Z-score indicated above the map. All analyses were performed on genes that were up-regulation more than 1.5-fold and down-regulated to below 0.67-fold in the microarray experiment. The categories of functions with marked responses are highlighted with a green box. A detailed comparison of Pt, St, and So is summarized in Fig. S7. **(b)** List of top enriched biofunctions in the comparison of Pt, St, and So. Biofunctions with an activation Z-score plus-minus over 2.0 are indicated and labeled in orange and blue, respectively. Gray dots show *p*-value > 0.05, with no statistical significance. The genes included in each functional category are listed in Fig. S8.

Is the Pt-induced regulation of gene expression homogenous in the entire cell population, or does it result from a specific cell population? Since cultured MSCs are typically heterogeneous ensembles, the differential gene expressions observed above cannot necessarily discriminate whether such regulation is derived from the majority of the cell population or from a limited subset of cells. Therefore, we next performed single-cell quantitative PCR to characterize the properties of MSCs that experienced Pt-culture (46 cells) in comparison to those in the usual CP-culture (46 cells) (Fig. 5a). Forty-eight assays were performed, including for a set of target genes found in the microarray data (*APC, DUSP6, ADGRG1*, etc.), genes involved in the stemness of MSCs (*OCT4, MEFLIN*, etc.), and genes of molecules involved in the therapeutic efficacy of MSCs (*CXCR4, IDO1, TSG6, IL6, BDNF*, etc.) (Table S4). The expression heat map showed two major hierarchical clusters: the Pt-cultured cell cluster showed a higher expression of *INTEGRIN β1, sTNFR1, APC, MFN2, NGF, GDNF, NEUROGRIN, SCF, COL1, MMP2, MEFLIN*, and *Ki67* than the CP-cultured cell cluster. These clusters were divided in the hierarchical analysis (Fig. 5a), tSNE plot, and violin plot (Fig. S9), indicating that the Pt-culture cells were clearly in different populations than those in the CP culture. *bFGF, NGF*, and *BDNF* were primarily up-regulated by qPCR even with an RNA sample of cell ensembles (Fig. S10a). Gene expression profiles were almost homogeneous in the Pt-cultured cell clusters; thus, the Pt-induced activation of cell functions applied to the whole cell population.

**Figure 5.**
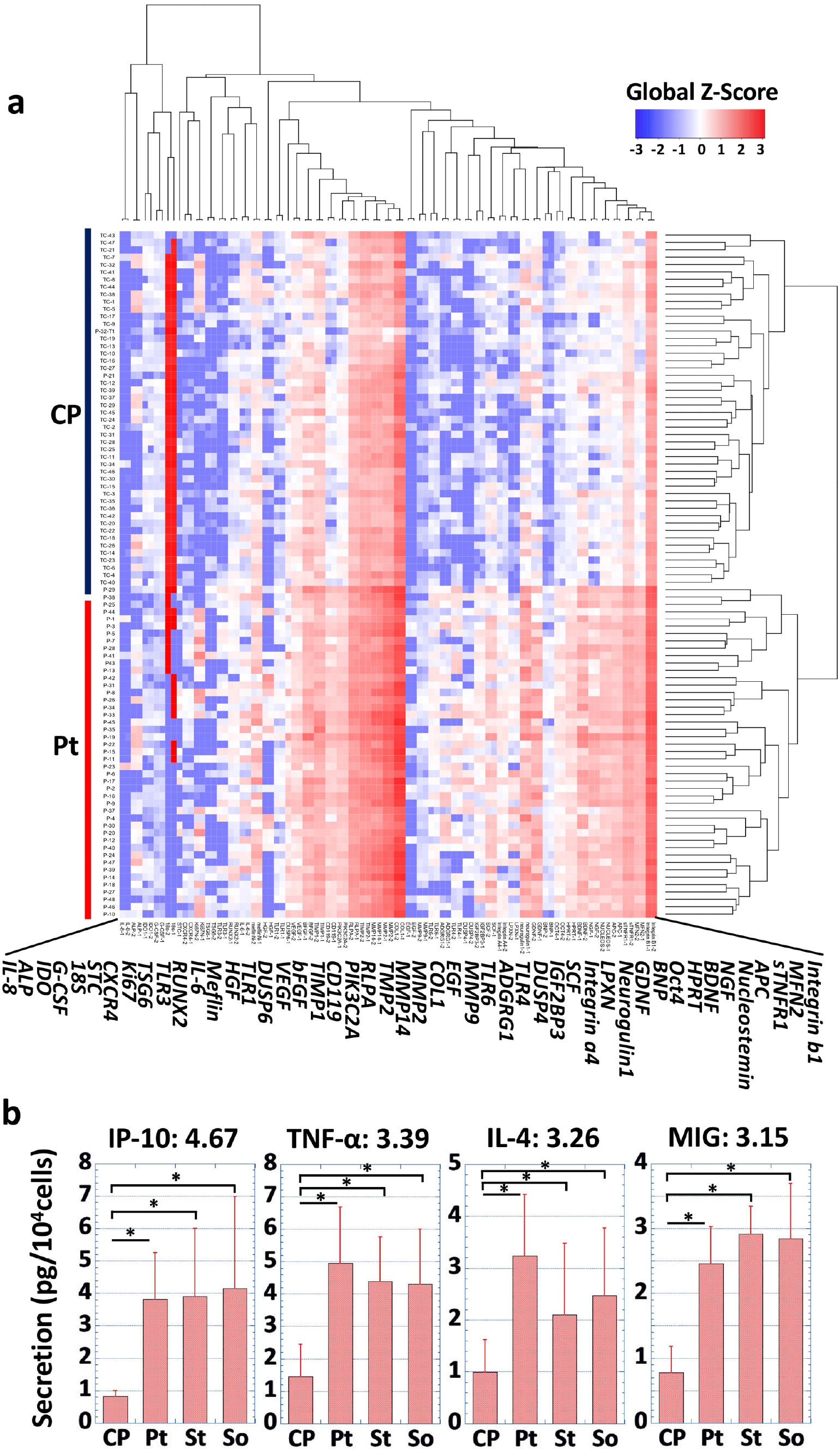
Function analysis of dual-durotaxing MSCs. **(a)** A single-cell analysis of gene expression related to a representative function of MSCs, including the selected top-increased genes, stemness-related genes, and therapeutic-property-supporting genes. Heat-map of expression with the Global Z-score. **(b)** Cytokine production by MSCs cultured on CP, Pt, St, and So. Means and SD for IP-10, TNF-a, IL-4, and MIG are shown. N=5. *: P<0.05. The values in the upper part of each graph show the mean fold-change of the secretion on Pt compared with CP. Results for other cytokines are summarized in Fig. S10.

To evaluate the impacts on therapeutic function, we characterized the secretion of cytokines from dual-durotaxing MSCs. The production of 11 immunomodulatory cytokines involved in the therapeutic efficacy of MSCs (IL-2, IL-4, IL-6, IL-8, IL-10, TNF, IFN-γ, RANTES, MIG, MCP-1, IP-10) was measured (Fig. 5b, Fig. S10b). Specifically, in Pt-cultured MSCs, IP-10, TNF-*α*, IL-4, and MIG production was significantly increased by 4.67-, 3.39-, 3.26-, and 3.15-fold compared to those in the CP-culture cells, respectively, confirming the possibility of an enhanced therapeutic benefit. The functional activation of dual-durotaxing MSCs was predicted for gene expression and some protein production related to the therapeutic effects.

### Dual-durotaxing-induced nuclear activation is caused by the mechanical linkage between the cytoskeleton and the cell nucleus

What mechanistic factors are involved in the above-characterized nuclear activation in dual-durotaxing cells? *APC* (adenomatous polyposis coli) was used as a marker for dual-durotaxing-induced nuclear activation in the inhibition experiments to assess the degree of such an activation response. It was the most up-regulated gene during Pt-culture found in the microarray measurement. Depletion of actomyosin tension decreased *APC* expression (Fig. S11a), suggesting that cytoskeletal force transmission is critical for nuclear activation. Inhibition of LINC complex was performed for LaminA/C of the inner nuclear membrane cortical structure and SUN1 (Sad1-UNP84) with each siRNA. Knockdown of Lamin A/C negated *APC* up-regulation on Pt gels, suggesting that linkage between the nuclear scaffold and chromatins is essential for transmitting the amplified ISD fluctuation inside the cell nucleus and the resulting nuclear activation (Fig. S11b). Interestingly, knockdown of SUN1, which connects the actin cytoskeleton and nuclear scaffold^41^, did not significantly affect the up-regulation of *APC* on Pt gels but did reduce *APC* expression on CP (Fig. S11b), which suggests that the amplified nuclear deformation may not simply be actomyosin-derived, but rather could result from the integrated dynamic contribution of a cytoskeletal cage fluctuating the nucleus.

How does the amplification of nuclear deformation modulate the expression of many genes involved in the activation of vigorous function of the cells? Chromatins directly bound to Lamin A/C in nuclear lamina that connects to SUN1 in the LINC complex should be subjected to the enhanced fluctuation of nuclear strain and stress, which may cause Pt-induced gene activation. The stress fluctuation in the nuclear membrane caused by the amplified fluctuation of ISD can propagate from the outer region toward the central region of the nucleus through interchromatin mingling, i.e., amplified fluctuation of ISD should enhance intranuclear stress fluctuation depending on a characteristic spatiotemporal distribution of chromatins. It has been revealed that the prestressed state of chromatin can affect cellular functions^42,43^, for example, through the structural modulations in the stress concentration site inside the nucleus, such as the intermingling region of chromatin territory^22^. In our system, cell shape was not confined to a particular micropattern and varied dynamically; therefore, it is natural to speculate that the dynamic cell-shape fluctuations exert extensive structural modulations of chromatins, which may affect the higher-ordered structure of stress concentration sites in various intranuclear hierarchical structures of chromatins such as the intermingling region between different chromosome territories, topologically-associated domains, and chromosome band boundaries. Nomadic cell migration among different regions of stiffness may interfere with the heterogeneous intranuclear mechanical structures and affect the regulation of gene expression at the structural boundaries with stress concentrations.

In summary, dual-durotaxing nomadic MSCs on a matrix with cell-scale stiffness-heterogeneity were found to have amplified long-term fluctuations in ISD, nuclear shaping, and spatiotemporal chromatin dynamics (schematically summarized in Fig. 6), which can lead to both the suppression of lineage bias to maintain stemness and the activation of vigorous cell functions. This cellular response was confirmed based on force transmission from the stiffness-heterogeneous matrix to the chromatin through the mechanical linkage between CSK and the nucleus. Significantly, therapeutic trophic effects that MSCs have as one of the most clinically useful stem cells^44^, such as anti-inflammatory, immunomodulatory, and tissue-protection effects, are typically attenuated by passage history involving lineage bias^45^. In this sense, matrix-mechanics-dependent lineage bias of MSCs is undesirable for maintaining its therapeutic properties due to the memory of mechanical stimulation from the culture environment^46^, the accumulation of associated mRNAs^47^, and the epigenetic history^48^. Since lineage specification results from lineage-specific tensional equilibrium as a local stable state as depicted in Waddington’s landscape^49^, avoiding tensional equilibrium with the amplified fluctuation of ISD should inhibit the lineage bias and assure the therapeutic quality of MSCs. A cell manipulation strategy for avoiding long-term tensional equilibrium in self-migrating stem cells was revealed to contribute to the refinement of stemness and functional activation.

**Figure 6.**
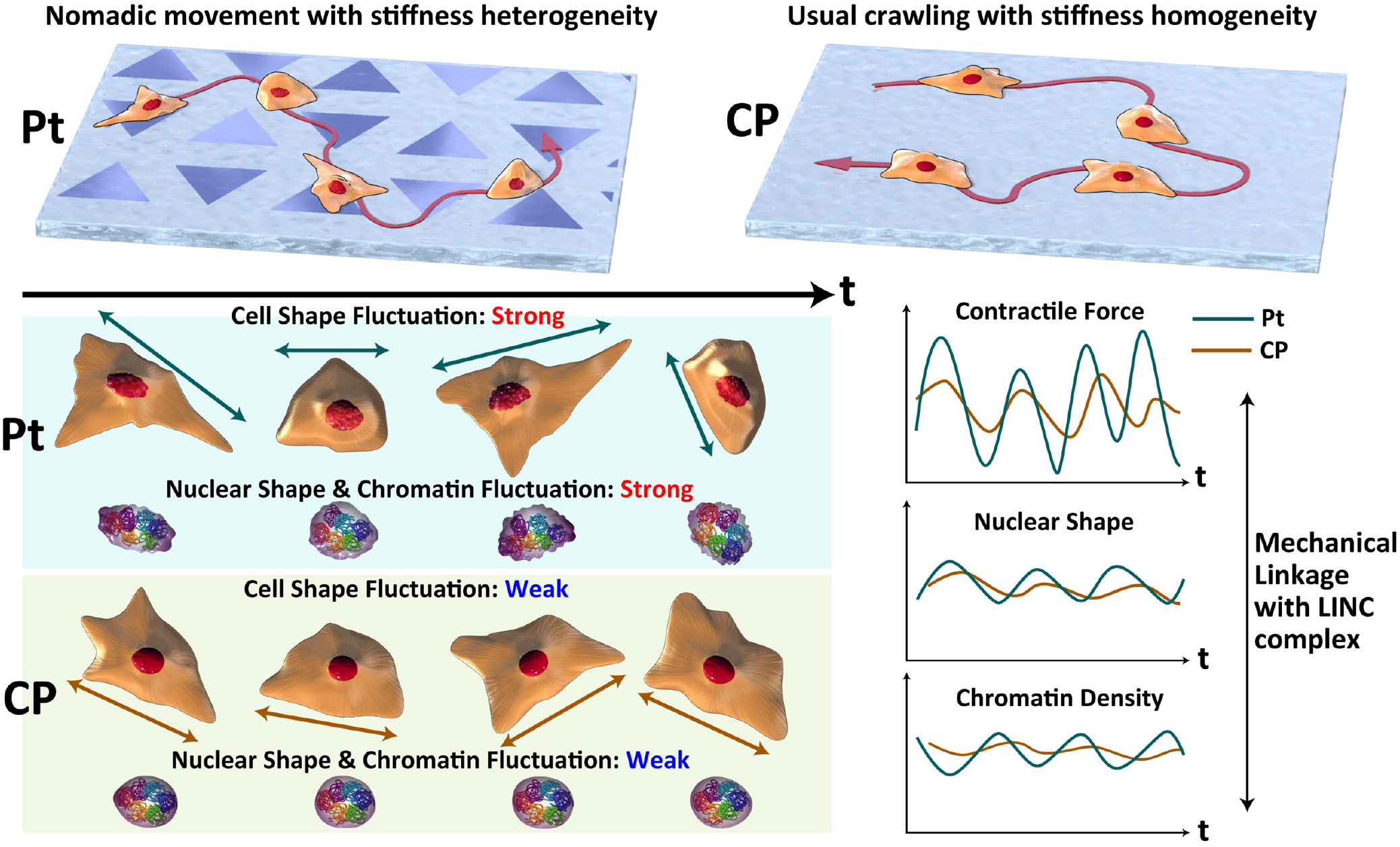
Schematic summary of the nuclear activation of MSCs through dual-durotaxing on a matrix with cell-scale stiffness-heterogeneity. When cells migrate on a matrix with cell-scale stiffness-heterogeneity, this should cause competing mixotactic cues on the matrix, e.g., dual-durotaxis of forward and reverse migration in this study, amplified ISD fluctuation is transferred to the cell nucleus and induces fast deformation of the nucleus and amplified mechanical fluctuation of chromatins. The avoided tensional equilibrium and the amplified chromatin fluctuation contribute to functional activation of the cells, as demonstrated in the present study.

## Methods

A complete description of the methods is attached.

## Acknowledgment

This research was supported by AMED-CREST under Grant Number JP20gm0810002.

## Author contributions

S. K. proposed the project, directed the research, discussed the findings with all members, and wrote the paper. H. E., K. M., T. O., and T. Kuboki prepared microelastically-patterned gels, measured the elasticity distribution by AFM, cultured MSCs on the gels, and performed time-lapse observation of cell migration. H. E. and K. H. conducted traction force microscopy and analyzed the data. T. Kuboki performed q-PCR analysis, constructed YAP-Venus, and conducted live-cell imaging for YAP-Venus and the cell nucleus. Y. T. cultured hMSCs, characterized stemness by flow cytometry and a differentiation experiment, and measured cytokine production. R. S. analyzed microarray data. S. S. supported AFM operations and analysis to measure the elasticity of gel samples. T. O. optimized and established the protocol to prepare the microelastically-patterned gels. T. Kawano first prepared microelastically-patterned gels and established the system for the time-lapse observation of cells on them. A. K. supported the preparation of plain gels with different stiffness values. R. S., K. K., and K. T. characterized the stemness of many lots of hMSCs. Especially, H. E. prepared and characterized about a thousand pieces of microelastically-patterned gels that were needed in this project.

## Methods

### hMSCs culture

Human bone marrow-derived MSCs (PT-2501, Lonza, Basel, Switzerland) were seeded at a density of 5.0 × 10^3^ cells/cm^2^ and expanded in mesenchymal stem cell growth medium (MSCGM^™^ BulletKit^™^, Lonza) on a tissue culture polystyrene dish (CP) (TPP Techno Plastic Products AG, Trasadingen, Switzerland) at 37℃ in a humidified atmosphere with 5% CO_2_. Culture medium was replaced every 3-4 days. Dulbecco’s Modified Eagle Medium (DMEM, low glucose, FUJIFILM Wako Pure Chemical Corp., Osaka, Japan) was basically used in all the experiments for the observation of migration and functional characterizations. MSCs at passages 4-6 were used for all experiments.

### Preparation of microelastically-patterned gelatinous gels

The microelastically-patterned gels were photolithographically prepared with photocurable styrenated gelatin (StG)^1,2^. Briefly, StG (30 wt.%) and sulfonyl camphorquinone (2.0 – 3.0 wt.% of gelatin; Toronto Research Chemicals, ON, Canada) were dissolved in phosphate-buffered saline (PBS), conditioned using an AR-100 deforming agitator (THINKY Corp., Tokyo, Japan). The StG sol solution was sandwiched and spread between vinyl-silanized glass substrates (vinyl-glass) and a normal glass substrate coated with poly(N-isopropylacrylamide) (PNIPAAm, Sigma Aldrich, St. Louis, MO, USA). Volumes of the spread sol solution were 30 and 90 μl for the vinyl-glass with diameters of 18 mm and 32 mm, respectively. As the first step, a soft base gel was prepared by irradiation of the entire sandwiched sol solution with visible light for 120s – 210s (45 – 50 mW / cm^2^ at 488 nm; light source: MME-250; Moritex Saitama, Japan). Next, stiff regions were prepared by irradiation of the base gel with equilateral-triangular irradiation patterns (side lengths: 150 μm, intercorner distances: 120 μm) by a custom-built, mask-free, reduction-projection-type photolithographic system using an EB-1770W liquid crystal display projector (Seiko Epson Corp., Nagano, Japan). To make the triangular irradiation patterns on large gels (32 mm diameter), we also used a photomask on an LED panel light (5.2 mW / cm^2^ at 488 nm; light source: LMG150×180NW; Aitec System Co., Ltd., Yokohama, Japan). Throughout these processes, the temperature of the StG sol and base gel was carefully kept above the LCST of PNIPAAm using a hot plate (Kitazato Corp., Fuji, Japan) so that the PNIPAAm polymers did not dissolve and intermingle with the top surface of patterned hydrogels. In addition, all these processes were performed in N_2_ chambers. Finally, the gels were detached from the sacrificial PNIPAAm-coated normal glass under room temperature and washed thoroughly with PBS at 28 °C over 12 hrs to completely remove adsorbed PNIPAAm. As we reported previously, the surface biochemical conditions were ensured to be the same for all the gel samples with different elasticities^2^. Therefore, the present experiments enabled us to investigate the pure biomechanical aspects of cell migration.

### Measurement of elasticity and topography of the gels

The surface elasticity of the StG gel was determined by nano-indentation analysis using an atomic force microscope (JPK NanoWizard 4, JPK Instruments, Bruker Nano GmbH, Germany). A commercial silicone-nitride cantilever with a nominal spring constant of 0.03 - 0.09 N/m was used (qp-BioAC-CI CB3, Nanosensors™, Neuchatel, Switzerland). Young’s moduli of the surface were evaluated from force-indentation curves by nonlinear least-squares fitting to the Hertz model in the case of a parabolic indenter (tip radius of curvature 30 nm; Poisson ratio: 0.45). To create a spatial distribution of elasticity of the triangular pattern gel, we measured tens of tiles with a 50 x 50 μm elasticity map and a distance of 2.5 μm between measured points. Next, we reconstructed the spatial distribution of elasticity from the tiles.

The topographic condition of the sample surface was also measured by using the atomic force microscope. Although the soft regions were higher than the stiff regions, the connection between the two regions was smooth enough to not disturb natural durotaxis. Previously, we reported that a height difference of less than several tens of microns did not affect the ability to induce durotaxis as long as the topographic connection in the elasticity boundary was appropriately smooth^2,3^.

### Pre-conditioning of StG gels for cell culture

The prepared StG gels were sterilized and equilibrated with a culture medium prior to all the experiments as follows. (1) The untreated StG gels were soaked in 70% aqueous ethanol solution and washed 100 times with paddling. (2) The gels were transferred to 0.2 µm filter-sterilized PBS and washed 100 times with paddling to remove ethanol. (3) The gels were transferred to new sterile PBS and stored in a 37°C incubator so as to be fully swollen until just before use. (4) The PBS was replaced with DMEM with 10% FBS, and the gels were fully equilibrated for 3hrs in DMEM at 37℃ under a 5% CO_2_ atmosphere in an incubator, and were finally ready for use.

### Time-lapse observation of cell migration and analysis of cell trajectories

The migration of cells on the patterned gel was observed using an automated all-in-one microscope with a temperature- and humidity-controlled cell chamber (BZ-X700; Keyence Corp., Osaka, Japan). Prior to the time-lapse observations, cells were seeded onto the gel at a density of 1 × 10^3^ cells / cm^2^ and cultured in DMEM containing 10% FBS for 18 – 22 hours under 5 % CO_2_. Phase-contrast images of cells were captured every 15 min for 24 h. For each experiment, we collected phase-contrast images from 11 positions of 2 gel samples. To eliminate the effect of cell-cell interaction and changes in chemical features of the gel surface due to cell product, we measured single-cell dynamics for one day after seeding at a low density.

Movement trajectories of the cells were analyzed using MATLAB software (MathWorks, MA, USA). Based on the edge-detection of a cell, we extracted the shape of the cell from phase-contrast images. The details of image processing have been explained previously^4^. We traced each cell and measured the time evolution of its trajectory. If the cells collided with each other or if a cell divided, we stopped the trace. When the cells separated again, we renumbered the cells and restarted the trace.

### Traction force microscopy

Traction force microscopy on a field of heterogeneous stiffness was performed according to the method that we previously reported^3^. Briefly, we first made gels with embedded beads. To embed the fluorescent beads near the patterned surface of the gels, a sacrificial glass was coated with the following ordered layers by using a spin coater: PNIPAAm, StG sol, and StG sol with fluorescent beads (Fluorospheres Carboxylate-Modified Microspheres, 0.2 μm, red fluorescent (580/605), Invitrogen, MA, USA). We added 0.1 wt.% Tween 20 to coat the StG sol solution to improve wettability. Next, we spread 25 μL of the StG sol solution between the vinyl-glass and the sacrificial PNIPAAm-coated glass. The procedure for photoirradiation was identical to that for gels without beads.

The images of the cells and fluorescent beads were monitored using an automated all-in-one microscope with a temperature- and humidity-controlled cell chamber (BZ-X700; Keyence). Phase-contrast images of the cells and fluorescent images of the beads were captured every 15 min for 24 h. After the time-lapse, we detached the cells by adding DMEM containing 0.3% Tween 20. We then measured the reference image of the fluorescent beads.

To calculate the traction force of the cells, we used finite-element-method (FEM)-based traction force microscopy. By comparing images with and without cells, the displacement of the beads was calculated using commercial PIV software (Flownizer 2D; Detect Corp., Tokyo, Japan). In the calculation of FEM, a hexahedron was used for discretization (the size of the unit is about 8 x 8 x 6 μm). The nodes of a hexahedron at the substrate surface were chosen to be identical to the nodes of PIV. We assumed that the patterned gel is a linear elastic body with spatial modulation according to Young’s modulus *E*(*x*):

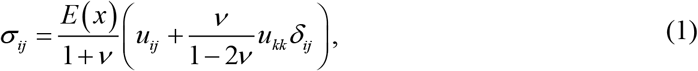

where *i*, *j*, and *k* are *x*, *y*, and *z*, respectively. *σ_ij_* and *u_ij_* are the stress tensor and strain tensor, respectively. *δ_ij_* is the Kronecker delta. Poisson ratio *ν* was set to be 0.45. For homogeneous gels, *E*(*x*) is constant. For patterned gels, we assumed

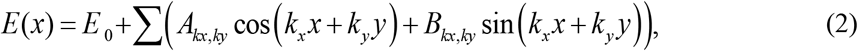

where *E_0_*, *A_kx,ky_*, and *B_kx,ky_* were fitted by the data from the AFM measurement, and *k_x_* and *k_y_* are the *x* and *y* components of the wave vector, respectively. *E_0_*, *A_kx,ky_*, and *B_kx,ky_* are mean elasticity and Fourier coefficients, respectively. With FEM, the equilibrium equation for a linear elastic body Eq. (1) was discretized. Next, we numerically calculated displacement *M_ijkl_* due to a unit force, where *M_ijkl_* is the *k* component of the displacement at node *i* when a unit force is applied in the *l* direction at node *j*. The displacement vector of the gel *U* induced by the traction force vector *F* is described as

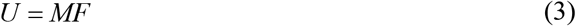

where matrix *M* is composed of *M_ijkl_*. *F* is reconstructed by minimizing a target function *S*, which is a combination of the least-square displacement error and a weighted norm of the forces:

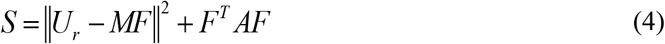

where *U_r_* and *A* are displacements of the substrate measured experimentally and the local penalty matrix, respectively. The minimum of *S*, *dS*/*dF* = 0, gives

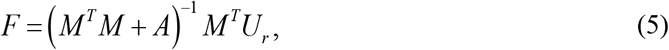

The local penalty matrix *A* is defined as follows^5^.

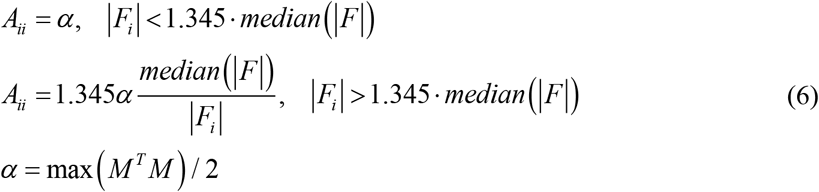

We repeatedly calculated Eqs. (5) and (6) until convergence was reached.

We calculated contractile force *CF* as the integral of the component of traction force *F* toward the centroid of the cell.

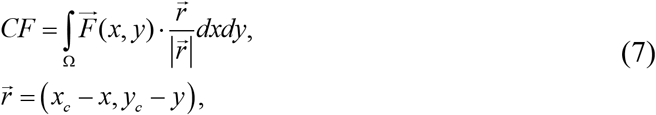

Where 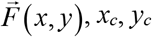 are the traction force at point (*x*, *y*) and the *x* and *y* coordinates of the cell centroid, respectively. The integral was calculated over the cell area *Ω*.

The magnitude of fluctuation of contractile force is defined as the normalized standard deviation of *CF* divided by the correlation time *τ_f_* of *CF*, σ (*CF*) */ τ_f_*〈CF〉, where ((*CF*))and 〈CF〉 are the standard deviation and time-average of *CF*. The correlation time *τ_f_* = ln 2 / *κ* of *CF* was calculated by fitting the autocorrelation function of *CF* by an exponential exp(-*κ*Δ*t*). To measure the strength of fluctuation, we also calculated the average magnitude of the time-derivative of normalized *CF*, 〈-|*ĊF*|〉/〈*CF*〉, which represents the rate of change of contractile force per unit time.

### Time-lapse observation of nuclear shaping dynamics and chromatin fluctuation

The MSCs were seeded onto the homogenous stiff (St: 20–40 kPa), soft (So: 2–4 kPa), patterned (Pt: stiff region: 20–40 kPa, soft region: 2–4 kPa) gels and CP at a density of 1,000 cells per cm^2^. After overnight cultivation, the cells were stained with NucBlue*™* Live ReadyProbe*™* (Molecular Probes, Eugene, OR, USA) according to the manufacturer’s instructions. After staining, the cells were washed twice with phenol-red-free CO_2_ independent Leibovitz’s medium (L-15, Gibco*™*, MA, USA). At least 3 hr before time-lapse observation, the cells were incubated in L-15 supplemented with 10 % FBS, penicillin/streptomycin, and 1 mM VectaCell Trolox*™* (Vector Laboratories, CA, USA). Fluorescent time-lapse observation of the nuclear dynamics was carried out on a BZ-X700 microscope using a 20X objective lens (4 stacking images, 10 μm layer). Movies were taken for 24 hr at 10 min intervals. Inhibitor treatment was performed by first culturing the cells on the substrates for 24 hr. The culture media was then replaced with the media containing 25 μM blebbistatin (FUJIFILM Wako Pure Chemical Corp., Osaka, Japan), 50 μM Y27632 (R&D Systems, MN, USA), 25 μM ML7 (Sigma Aldrich, MO, USA), or 0.4 μM cytochalasin D (Sigma Aldrich, MO, USA). After overnight incubation, the cells were stained with NucBlue*™*, and time-lapse observation was performed in the presence of the inhibitors and Trolox. For the observation of chromatin fluctuation, time-lapse analysis was performed with similar procedures as nuclear shaping dynamics except that the fluorescent intensity of the nuclei was observed using a 20X objective lens with 2X optical zoom for 12 hr at 10-min intervals.

### Analysis of nuclear shaping dynamics

The shaping dynamics of the cell nuclei were analyzed using MATLAB software. First, we adopted a Gaussian filter with a cut-off frequency of 0.13 μm^-1^ for the fluorescent image of the nuclei. Next, we adopted a top-hat filter to eliminate the background. The filtered images were binarized with the threshold determined based on the maximum intensity of an image. We traced each nucleus and measured the time evolution of the trajectory and shape. If the nuclei collided or if a nucleus replicated, we stopped the trace. When the nuclei separated again, we renumbered the nuclei and restarted the trace. We only analyzed data that accumulated for longer than 12 hr. From binarized images, we calculated the aspect ratio of the nuclei. The binarized nuclei were fitted by the ellipsoid through the function implemented in MATLAB, and the aspect ratio was calculated as the ratio of long and short axes of the fitted ellipsoid. Next, we analyzed the time series of the aspect ratio of the nuclei. By fitting the autocorrelation function of the aspect ratio by an exponential exp(-*κΔt*) the correlation time *τ_f_* = ln2/*κ* was determined. Similar to the analysis of contractile force, the normalized standard deviation divided by correlation time, *σ*(*As*)/*τ_f_*〈*As*〉, was defined as the magnitude of the fluctuation of the nuclear aspect ratio *As*.

### Characterization of fluctuation of chromatin distribution

Fluctuation of the chromatin distribution was calculated by an analysis of the fluorescence intensity of the nuclei. Similar to the analysis of nuclear shaping dynamics, we binarized the fluorescent image to determine the interior of the nucleus. We calculated the standard deviation of the fluorescence intensity *FI*(*x*,*y*) inside a nucleus at time *t*, *σ_spa_*(*FI*(*x*,*y*)), which represented the spatial heterogeneity of the chromatin distribution. Next, we traced the nuclei and measured the time evolution of *σ_spa_* (*FI*). We defined the normalized standard deviation, *σ_temp_*[*σ_spa_*(*FI*)]/〈*σ_spa_* (*FI*)〉*_temp_*, as the magnitude of fluctuation of the spatiotemporal distribution of chromatins.

### Immunofluorescence staining of YAP

hMSCs were seeded onto StG gel at 2500 cells/cm^2^, cultured for 4 days, fixed with 4% paraformaldehyde (FUJIFILM Wako) for 15 min, then permeabilized and blocked with PBS containing 0.5% Triton X-100 (Sigma Aldrich, MO, USA), 10% donkey serum (EMD Millipore Corp., CA, USA), and 1% albumin from bovine serum (FUJIFILM Wako) for 45 min. The cells were incubated with a primary antibody, 10 μg/ml rabbit anti-YAP (GTX129151, Genetex, CA, USA) at 4°C overnight. The samples were washed with PBS-0.05% Tween 3 times for 15 min each. Subsequently, the cells were incubated with a secondary antibody (donkey anti-rabbit conjugated with Alexa Fluor 488 (1:1000; Invitrogen) for 60 min at 37°C. After washing, the fluorescent images were acquired using a BZ-X700 microscope.

### Quantitative image analysis of YAP localization

To quantify the magnitude of YAP localization, we calculated the ratio of fluorescence intensity between the nucleus and cytosol (Nuc/Cyt YAP ratio) from fluorescent images of YAP. The intensity of the nucleus was defined by the mean intensity of the region that satisfied *D* < 12 μm, where *D* represented the distance from the center of the nucleus. As for the intensity of cytosol, we used the mean intensity of the region that satisfied 12 < *D* < 24 μm. We defined nuclear localization of YAP for a Nuc/Cyt YAP ratio > 1.6 and cytosol localization for a Nuc/Cyt YAP ratio < 1.2, respectively.

### Cloning and expression of YAP fusion with Venus

hMSCs were cultured and used for RNA preparation using TRIzol*™* reagent. Reverse transcription was performed using random hexamers with high-fidelity PrimeScript Reverse Transcriptase (Takara Bio Inc., Tokyo, Japan). The cDNA of YAP was obtained by RT-PCR with specific primers flanking the coding regions using PrimeStar^®^ Max DNA Polymerase (Takara Bio). The expected PCR products were digested from the agarose gel and purified with a GENECLEAN^®^ II kit (MP Biomedical, CA, USA). The purified products were phosphorylated by kinasing and blunt-end ligating to the HincII/BAP ends of pUC118 (Takara Bio) using a DNA ligation kit (Mighty Mix, Takara Bio), according to the manufacturer’s instructions. The ligation reaction was used for transformation into *E.coli* DH5 alpha and plated onto LB-ampicillin supplemented with X-gal (FUJIFILM Wako) and IPTG (FUJIFILM Wako). The positive clones were selected for plasmid preparation using a QIAGEN plasmid mini preparation kit (Qiagen, Hilden, Germany) and sequencing.

The pEF1-Venus was prepared as previously described^6^. Five micrograms of the plasmid were digested with *EcoR*V (Roche Diagnostics, Rotkreuz, Switzerland) and purified by agarose gel electrophoresis. The purified product was dephosphorylated with calf intestinal alkaline phosphatase (Roche Diagnostics) to prevent self-ligation.

The pUC118 plasmids harboring YAP were used as the template for PCR amplification. The 7-alanine amino acids were introduced to serve as a linker to facilitate correct folding of the fusion protein. Primers containing *EcoRV* restriction site (underlined) and nucleotides encoding for 7 alanine (italics) were used for PCR amplification (forward 5’ GC G<u>GA TAT C</u>CA *GCT GCT GCT GCA GCT GCT GCA* ACC **ATG** GAT CCC GGG CAG 3’, reverse 5’ G CG<u>G ATA</u> <u>TC</u>T **CTA** TAA CCA TGT AAG 3’) for cloning in-frame with the C-terminus of pEF1-Venus. The start codon (ATG) and stop codon (CTA) are indicated in bold. The PCR products with expected sizes were digested from the gel and purified using a GENECLEAN^®^ II kit.

The purified vector and inserts were ligated at 16°C for 4 hr using a Mighty cloning reagent kit (Takara Bio), and the ligation products were then used for transformation. Positive colonies grown on LB-kanamycin plates were selected for plasmid DNA preparation and sequencing.

### Live-cell imaging of nucleocytoplasmic shuttling of YAP

hMSCs were seeded on Pt gels attached inside a 35mm dish at a density of 2,500 cells/cm^2^ and cultured overnight. At 3-5 hr before transfection of YAP-Venus, the culture medium was replaced with fresh medium supplemented with 30 μg/ml radical-containing nanoparticles (RNPs). Transfection was performed using Lipofectamine^TM^ Stem transfection reagent (Invitrogen). The plasmid DNA 1-2 μg was first mixed with 100 μl of Opti-MEM medium. The Lipofectamine Stem reagent (4-8 μl) was mixed with 100 μl of Opti-MEM in separate tubes. The DNA mixture was then transferred to the Lipofectamine suspension, mixed and incubated for 5 min at room temperature. The DNA-Lipofectamine complex (140 μl) was dropped into the MSCs cultured on the gels and incubated at 37°C overnight. After overnight transfection, the culture medium was replaced with L-15 supplemented with 30 μg/ml RNPs. The time-lapse observation was performed using a 20X objective lens of BZ-X700 with 1.5X optical zoom (4 stacking images, 3 μm layer). Movies were taken for 2 days at 15 min intervals.

### Flow cytometry of CD marker expression

The cells were harvested with Accutase (Innovative Cell Technologies, Inc., CA, USA) from culture on the StG gels, passed through a cell strainer with 70 μm mesh, washed with FACS buffer (0.5% BSA/PBS, 2mM EDTA), and collected by centrifugation at 300×g for 5 min. The collected cells were fluorescently stained with the following mouse anti-human primary antibodies for 30 min on ice: Positive markers; CD73 (Miltenyi Biotec, Germany), CD90 (R&D Systems), and CD105 (Santa Cruz Biotechnology Inc., TX, USA). Negative markers; CD11b (MilliporeSigma, MA, USA), CD19 (R&D Systems), CD34 (Santa Cruz), CD45 (Abcam plc., Cambridge, UK), and HLA-DR (Santa Cruz). After being washed to remove primary antibodies, the cells were reacted with fluorescein isothiocyanate (FlTC)-labeled secondary anti-mouse antibodies for 30 min on ice (Alexa 488-anti mouse IgG, Invitrogen). The samples were measured with a flow cytometer (Guava® easyCyte®) with a termination count of 5000 cells.

### Differentiation induction of hMSCs on the gel samples

Differentiation of hMSCs on the gel samples was assayed as follows: (1) hMSCs were seeded on the gels at a density of 2,000 cells/cm². (2) To make the cells experience the stiffness stimulus from the gels, preculture on the gels was performed for 7 days with DMEM medium changed once during culture. (3) After the preculture, the medium was replaced with an induction medium for osteogenic or adipogenic differentiation. (4) To evaluate the maintenance of differentiation ability after culture on the gels, the cells were detached from the gels with trypsin after 7 days of culture, re-seeded onto CP, and then cultured with induction medium for osteogenic or adipogenic differentiation for 2 or 3 weeks.

Osteogenic induction medium was prepared by adding dexamethasone (final. conc: 100 nM), 2-phospho-L-ascorbic acid trisodium salt (final. conc: 50 μg/ml), *β*-glycerol phosphate disodium salt pentahydrate (final. conc: 10 mM) to *α* MEM basal media (FUJIFILM Wako) including 10% FBS and 2M L-glutamine. Adipogenic induction medium was prepared by adding adipogenic supplement (sc006, R&D Systems) to *α* MEM basal media. The medium was changed to a fresh induction medium every 3-4 days.

### Staining with Alizarin Red S and Oil Red O

The formation of mineralized matrix nodules in osteogenic induction and lipid vacuoles in adipogenic-induced cells was assessed by staining with Alizarin red and Oil Red O, respectively. In brief, the cells after differentiation-induction culture were fixed in 4% paraformaldehyde PBS solution for 15 minutes at room temperature, rinsed three times with distilled water, treated with Alizarin Red S solution (Sigma-Aldrich) for 10 minutes under shaded light, or with Mix Oil Red O saturated solution (FUJIFILM Wako) for 15 minutes at room temperature, and then rinsed three times using distilled water to reduce nonspecific staining. The stained samples were observed in PBS with an all-in-one fluorescence microscope (BZ-X700, Keyence).

### Quantitative image-analysis of osteogenic and adipogenic differentiation

The degree of osteogenic differentiation was quantified by analyzing the RGB color of the bright field image of Alizarin red staining. At the early stage of osteogenic differentiation, the samples were stained red. On the other hand, control gels (cells without osteogenic differentiation medium) were stained to be violet, while the control CP became almost white. Thus, to quantify the early stage of osteogenic differentiation, we used the spatially averaged value of *ΔI* = *I_r_* -*I_b_*, where *I_r_* and *I_b_* were the intensity of the red and blue components of the image, respectively. *ΔI* became almost 0 for control samples, and *ΔI* had a large value for red-stained images. As osteogenic differentiation proceeded, the stained color became darker, which caused a decrease of *ΔI*. Therefore, to quantify the late stage of osteogenic differentiation, we used *ΔI’* = 1- (*I_r_* - *I_b_*) instead. For all the conditions, we divided the samples into 8 ∼ 9 sub-regions and calculated *ΔI* or *ΔI’* for each sub-region. To eliminate the contribution of the background color, we subtracted *ΔI* of the control samples from the data.

To quantify the degree of adipogenic differentiation, we calculated the ratio of the cells stained by oil red O. To determine the stained cells, we classified the stained and non-stained areas from the bright field image of oil red O staining. For the machine learning-based classifier, we used a support vector machine (MATLAB, MathWorks). As training data, we used five typical images of stained areas and the background containing non-stained cells for each sample. The software learned the representative distributions of the intensities in RGB space and calculated the separate plane in RGB space that classifies the two groups^7^. After training, stained and non-stained areas of the image were classified by the software. Next, we considered that the cell was differentiated when the stained area was included in the region *D* < 2*R_n_*, where *D* and *R_n_* represented the distance from the center of the nucleus and the average radius of the nucleus. We then counted the numbers of differentiated cells and calculated the ratio of the numbers of differentiated cells and total cells. For all the conditions, we divided the samples into 8 ∼ 9 sub-regions and calculated the ratio of differentiated cells in the sub-regions.

### DNA microarray and bioinformatic analysis

Whole-genome expression was analyzed after 4 days of culture of hMSCs on three kinds of gel (P, St, and So) or CP. Total RNA (n=3, biological replicates) was isolated from hMSCs cultured on each sample using a miRNeasy Micro Kit (Qiagen). Total RNA quantity was assessed on a NanoDrop 2000 (ThermoFisher Scientific, Waltham, MA USA), and its quality was assessed on an Agilent 2100 Bioanalyzer (Agilent Technologies, CA USA); 100 ng of total RNA was used to generate biotin-modified amplified RNA (aRNA) with the GeneChip 3’IVT PLUS Reagent Kit (ThermoFisher). Reverse transcription of first-strand complementary DNA (cDNA) with a T7 promoter sequence was performed with T7 oligo(dT) primer. Second-strand cDNA synthesis was used to convert the single-stranded cDNA into a double-stranded DNA template. The reaction used DNA polymerase and RNase H to degrade the RNA and synthesize second-strand cDNA simultaneously. In vitro transcription of biotin-modified aRNA with IVT Labeling Master Mix generated multiple copies of biotin-modified aRNA from the double-stranded cDNA templates. The aRNA was purified and quantified; after fragmentation, it was hybridized to the GeneChip Human Genome U133 Plus 2.0 Array (ThermoFisher). The arrays were stained with phycoerythrin and washed at the GeneChip Fluidics Station 450 (Affymetrix, CA USA). The microarrays were scanned, data were extracted using GeneChip scanner 3000 7G (Affymetrix), and images were analyzed using the Affymetrix GeneChip Command Console Software and digitized using Affymetrix Expression Console.

The results from microarray analysis were uploaded into Qiagen Ingenuity® Pathway Analysis (IPA) for core analysis with the IPA knowledge base. IPA was performed to obtain the responses in canonical pathways, the candidates of upstream regulators, and the predicted modulation in biofunctions of the tested cells. A comparative analysis was conducted between the cells cultured on Pt, St, and So gels.

### Single-cell qPCR

The cells cultured on StG gels (Pt, St, and So) and CP at a density of 3,000 cells per cm^2^ for 4 days were trypsinized, counted, and filtered through a 40 μm cell strainer. The cells were resuspended in complete DMEM culture media to make the final concentration approximately 1-2 × 10^5^ cells per ml. Single-cell separation was performed using the C1 system (Fluidigm, CA, USA). The cell suspension was mixed with the cell suspension reagent from the C1 Single-Cell Auto Prep Reagent kit (Fluidigm) at a ratio of 3:2 (cell suspension 6 μl + cell suspension reagent 4 μl). Next, 6 μl of the cell mixture was loaded onto the primed Integrated Fluidic Circuit (IFC, C1 Single-Cell Auto Prep for Preamp 17-25 μm, Fluidigm), and cells were then captured into the IFC according to the manufacturer’s instructions.

The forward and reverse gene-specific primers were pooled to a final concentration of 500 nM for 48 target genes. The Ambion Single Cell-to-CT qPCR kit (Invitrogen), C1 Single-Cell Auto Prep Reagent kit, and the pooled primers were used to prepare Lysis Final Mix, RT Final Mix, and Preamp Final mix reactions. According to the manufacturer’s instructions, cell lysis, reverse transcription, and preamplification were performed in the C1 system. After preamplification was complete, the amplicon was harvested from the inlets and transferred to 25 μl of C1 dilution reagent. The diluted cDNA was then loaded for real-time PCR using Biomark HD (Fluidigm), which finally provided expression arrays of 48 genes on 48 cells. The obtained data were analyzed with the Singular Analysis Toolset (Fluidigm) on R software, to give expression heat maps, hierarchical clustering, and a principal component analysis.

### Cytokine production assay

Cytokine production was characterized using a bead array (Cytometric Beads Array, Becton Dickinson, NJ, USA) according to the manufacturer’s protocol. Briefly, the supernatants of media of MSCs cultured on StG gels and CP were centrifuged at 430×g for 3 min, passed through a 0.2 μm filter, thoroughly mixed with the capture beads (Human Chemokine Kit (IL-8, RANTES, MIG, MCP-1, IP-10); Human Th1/Th2 Cytokine Kit II (IL-2, IL-4, IL-6, IL-10, TNF, IFN-γ)), and then incubated for 3 hours at room temperature, protected from light. Serially diluted standards of capture beads were prepared simultaneously. The sample solutions of reacted beads were centrifuged at 200×g for 5min, washed once with a wash buffer, and resuspended in a wash buffer. Finally, the resuspended bead samples were acquired and analyzed with a flow cytometer (Guava® easyCyte™, Luminex, TX, USA).

### Assessment of nuclear activation under inhibitors and knockdown treatment

hMSCs were seeded at a density of 2,000 cells/cm^2^ on CP and Pt gels in 12-well plates and cultured for 4 days. On day 3, the test groups were treated with 25 μM blebbistatin (FUJIFILM Wako) or 50 μM Y-27632 (R&D Systems) for an additional one day before RNA isolation. LINC complex was disrupted by siRNA treatment of SUN1 (Santa Cruz Biotechnology, TX, USA), LAMIN A/C (Santa Cruz Biotechnology), and control scramble (Santa Cruz Biotechnology). siRNA transfection was performed after 3 days of culture using Lipofectamine RNAiMAX^®^ Reagent (Invitrogen). The siRNA (20 pmol) was mixed with 100 μl Opti-MEM I Reduced Serum Medium (Gibco*™*, MA, USA). In separate tubes, 6 μl of RNAiMAX^®^ Reagent was diluted with 100 μl Opti-MEM medium. The diluted siRNA was mixed with the diluted Lipofectamine RNAiMAX^®^ Reagent at a ratio of 1:1, and 100 μl of the mixture (10 pmol of siRNA per well) was dropped onto the cells after 5 min of incubation at room temperature. After one day of treatment with siRNAs, the samples were collected for RNA isolation using the TRIzol*™* reagent (Invitrogen). RNA preparation was performed using a Direct-zol*™* RNA Microprep kit (Zymo Research, CA, USA). One microgram of total RNA was used for DNase digestion using Turbo*™*DNase (Invitrogen). The DNase-treated RNA was further used with an iScript*™*

Advanced cDNA synthesis kit (BioRad, CA, USA). A quantitative real-time polymerase chain reaction (qPCR) was performed using SsoAdvanced*™* Universal SYBR^®^ Green Supermix (BioRad) in a CFX Connect real-time PCR detection system. The expression of the gene of interest was characterized using the standard curve method. The reference genes Hypoxanthine Phosphoribosyltransferase1 (HPRT1), Ribosomal Protein L13A (RPL13A), and Glyceraldehyde-3-Phosphate Dehydrogenase (GAPDH) were used for the normalization of gene expression.

### Statistical analysis

For all the statistical analyses, Kruskal Wallis test followed by post-hoc test with Mann–Whitney U test was conducted. The number of gel samples and cells tested are described in the legend of each figure. P values were calculated using MATLAB software.

## Supplementary Information

### Supplementary Figures

**Figure S1.**
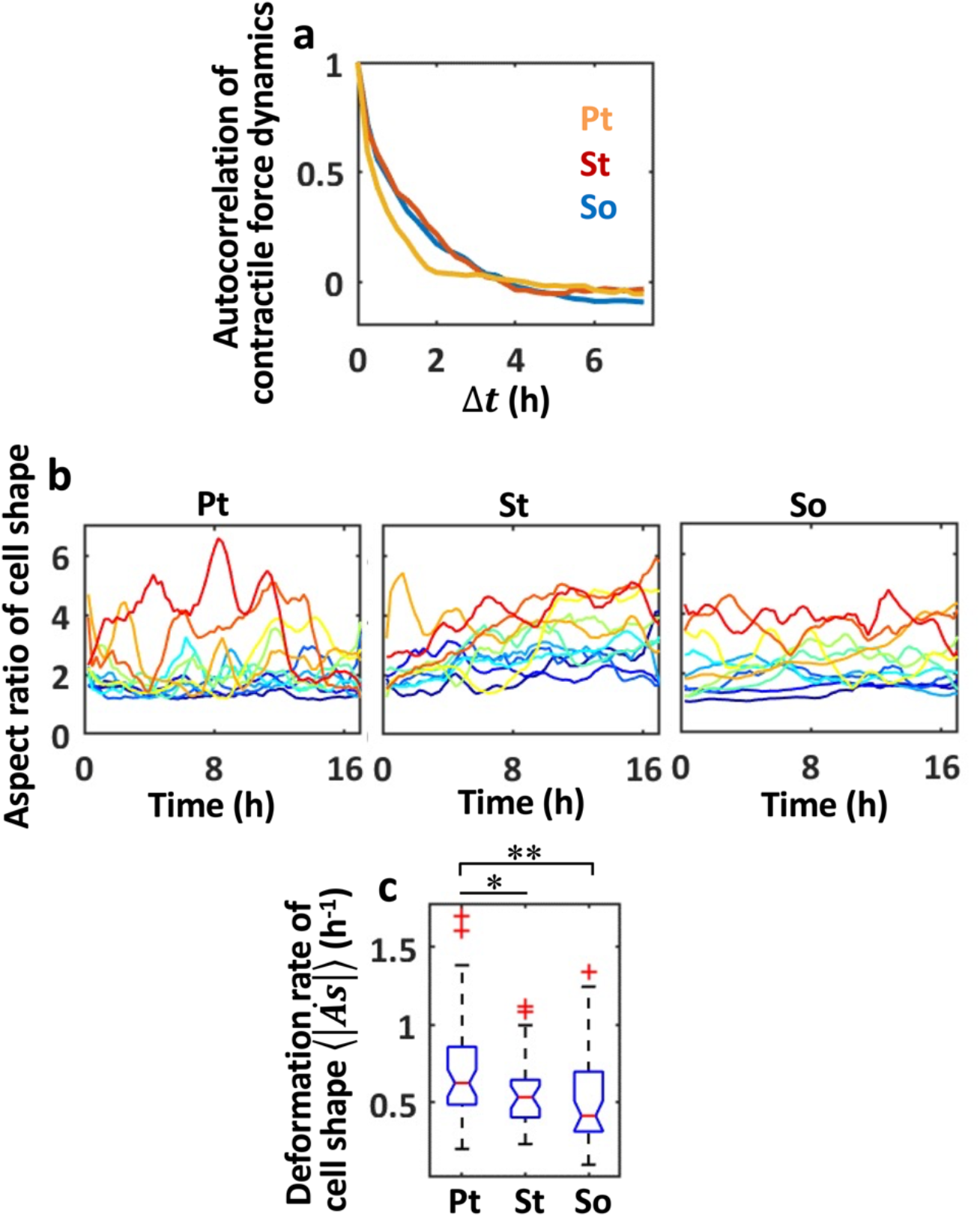
Analysis of dynamics in contractile force and shape of the cells cultured on Pt, St, and So gels. **(a)** Autocorrelation function of time variation of contractile forces described in Fig. 1i. Correlation time *τ_f_* in Fig. 1j was determined from these data. **(b)** Representative time variation of the aspect ratio of cell shape. n=11 for Pt, St, and So. **(c)** Deformation rate of cell shape calculated from (b). The intensity of fluctuation of cell shape was highest in dual-durotaxing cells on the Pt gels. n= 46 (Pt), 28 (St), 39 (So). **: P<0.01. *: P<0.05.

**Figure S2.**
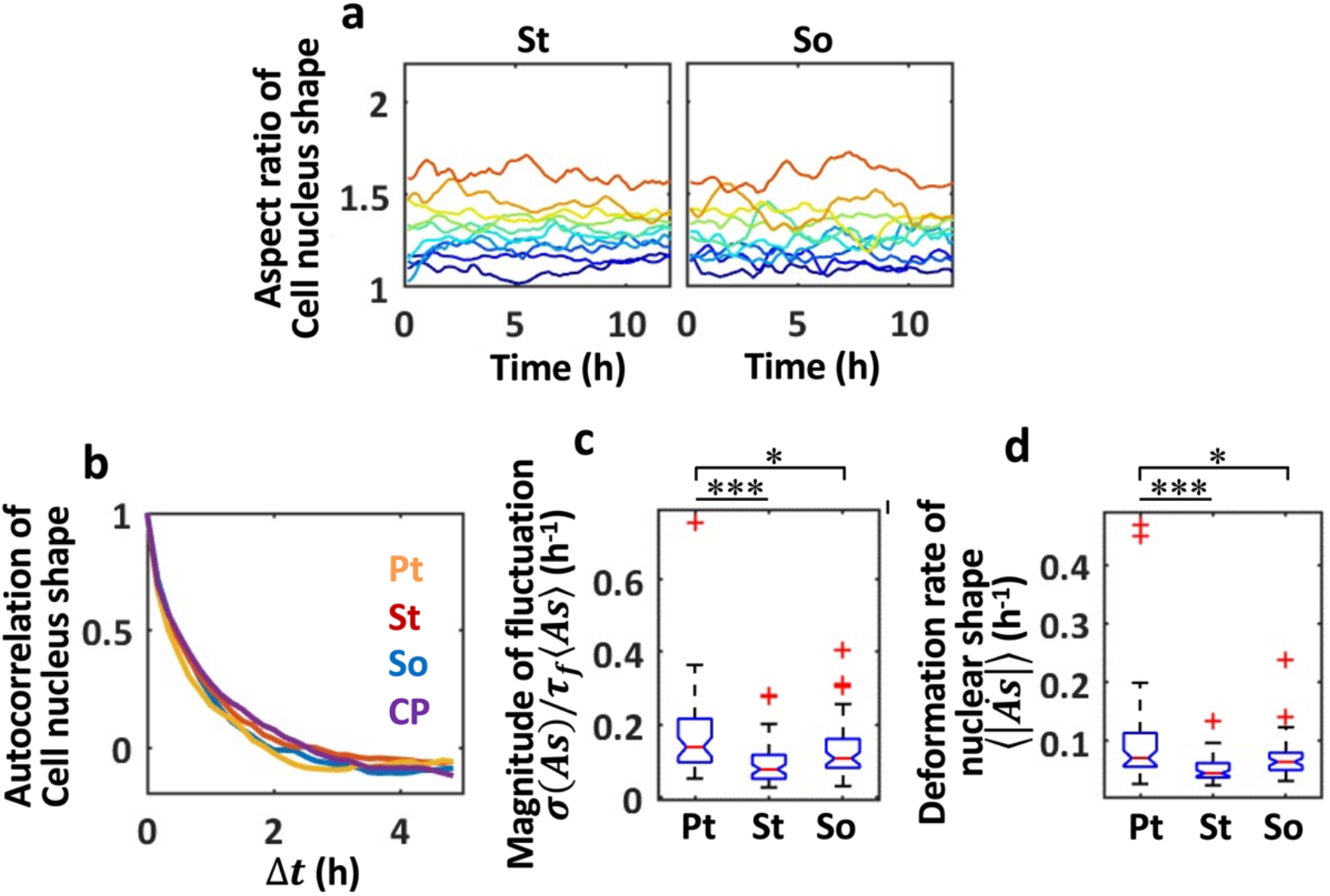
Analysis of dynamics in nuclear shape of cells cultured on Pt, St and So gels, and on CP. **(a)** Representative time variation of the aspect ratio of the cell nucleus on St and So gels. n=10 for St and So. **(b)** Autocorrelation functions of the time variation of nuclear shape described in Fig. 2a and Fig. S2a. **(c, d)** The magnitude of fluctuation (c) and the deformation rate (d) in the nuclear shaping fluctuations. The degree of fluctuation and the deformation rate of nuclear shape were defined as the ratio of normalized standard deviation to correlation time, and the time derivative of fluctuation, respectively. ***: P<0.001. *: P<0.05. n= 51 (Pt), 59 (St), 56 (So).

**Figure S3.**
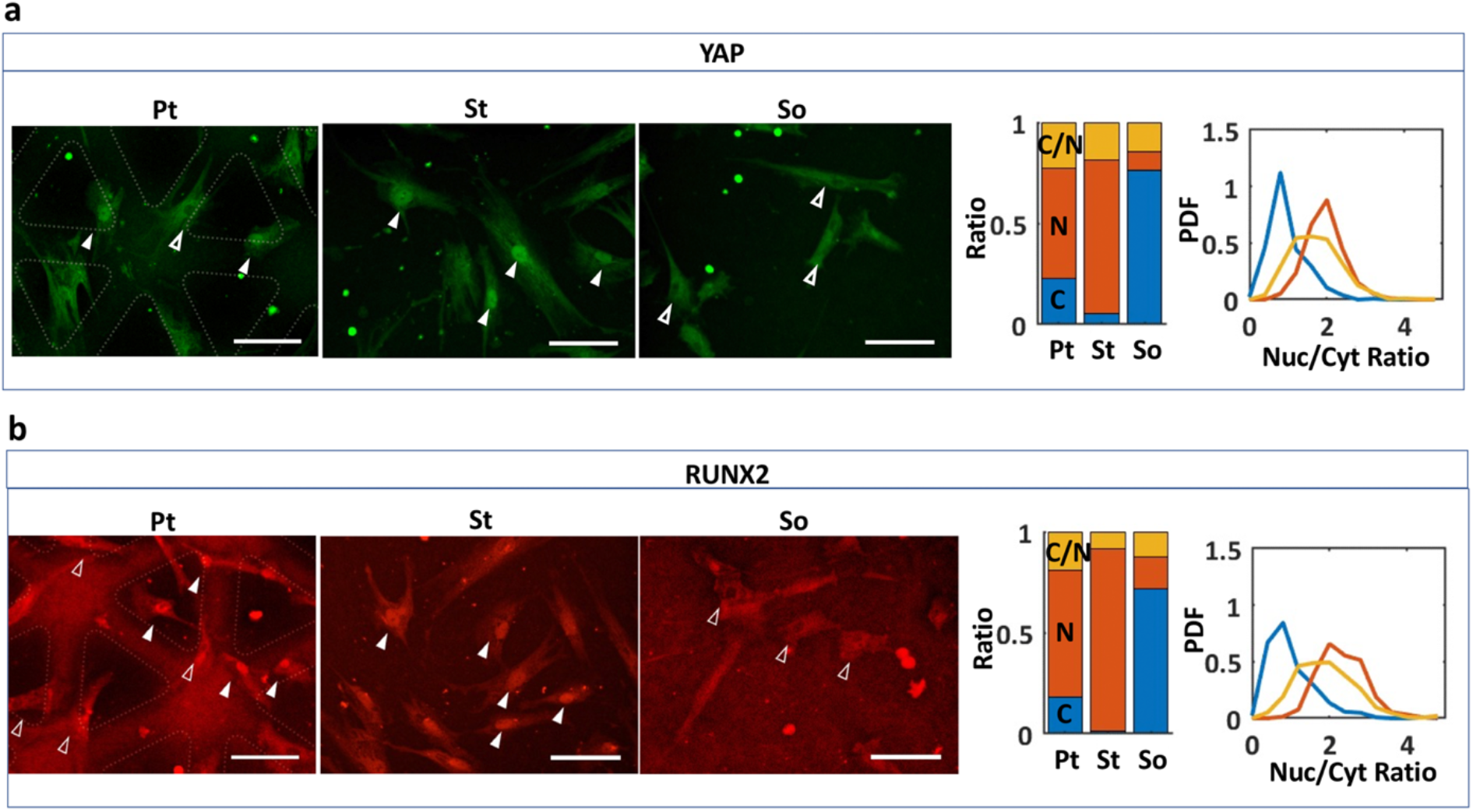
Intracellular localization of YAP (a) and RUNX2 (b) in cells cultured on Pt, St, and So gels. **Left:** immunofluorescence microscopic images. Solid arrows; nuclear localization. Open arrows; cytoplasmic localization. Scale bars: 100 μm. **Middle:** cellular ratio measured for cytoplasmic localization (C), nuclear localization (N), and cytoplasmic/nuclear co-localization (C/N). n= 497 (Pt), 314 (St), 384 (So). **Right:** probability density functions of the ratio of nuclear to cytoplasmic localization.

**Figure S4.**
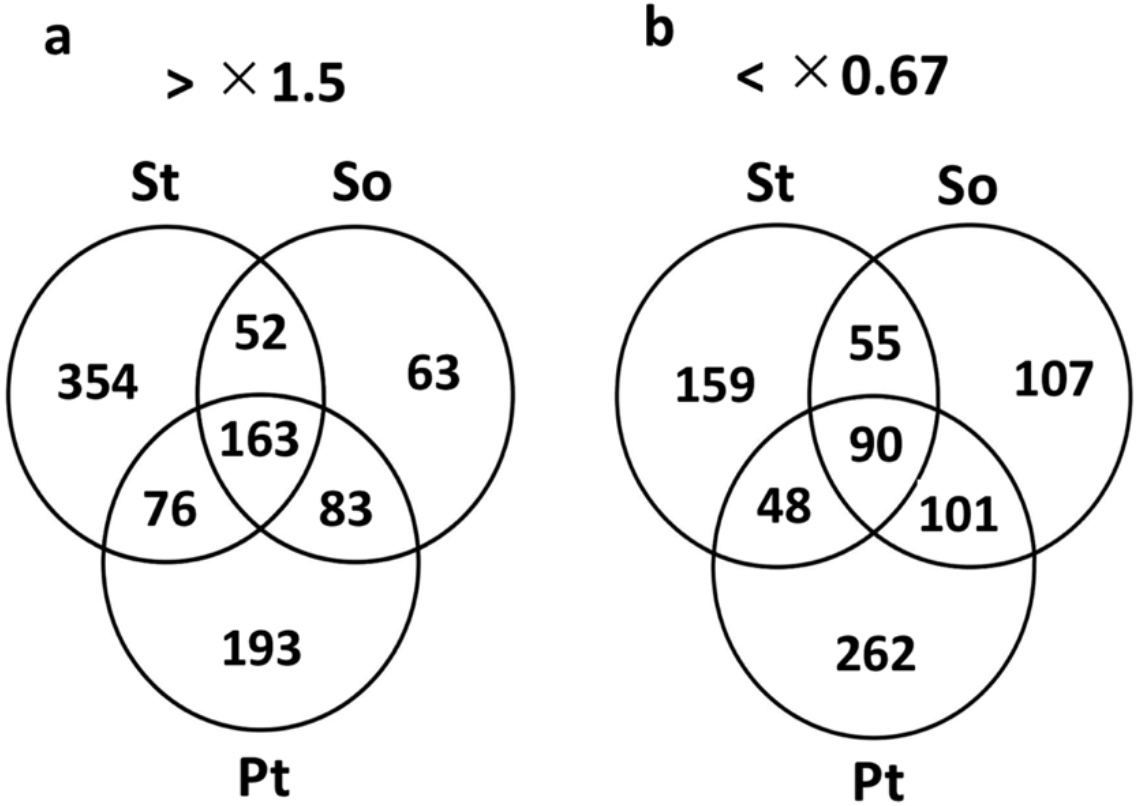
Venn diagrams of up-regulated (a; >1.5-fold relative to CP) and down-regulated (b; <0.67-fold relative to CP) genes analyzed by DNA microarrays with cells cultured on Pt, St, and So gels.

**Figure S5.**
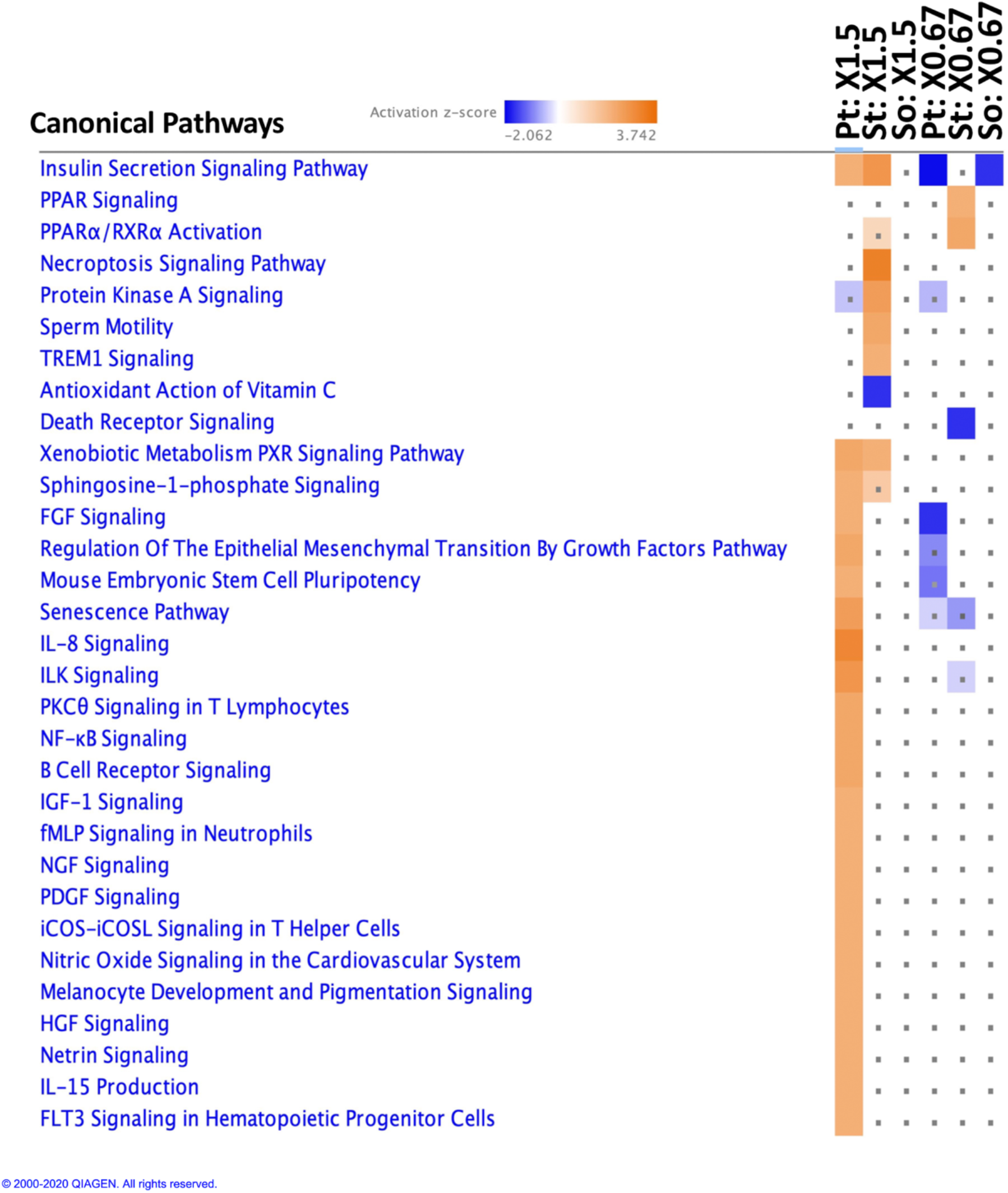
Comparison of the representative canonical pathways predicted to be activated or inactivated based on each expression profile in the DNA microarray data observed in cells cultured on Pt, St, and So gels. Generated by IPA software. Grey dots show insignificant with P > 0.05.

**Figure S6a.**
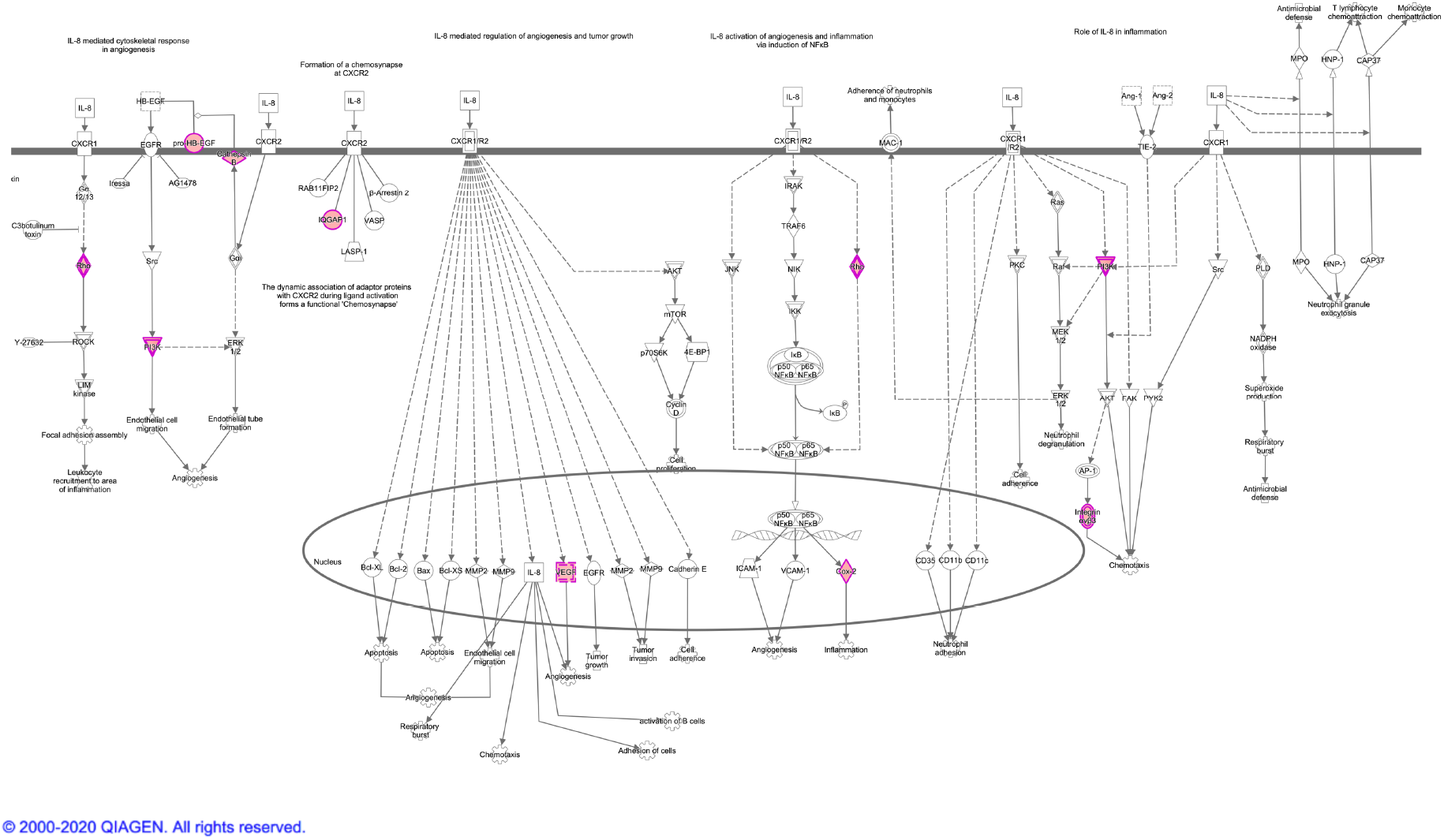
IL-8 signaling pathway map including the observed up-regulated genes (highlighted in pink).

**Figure S6b.**
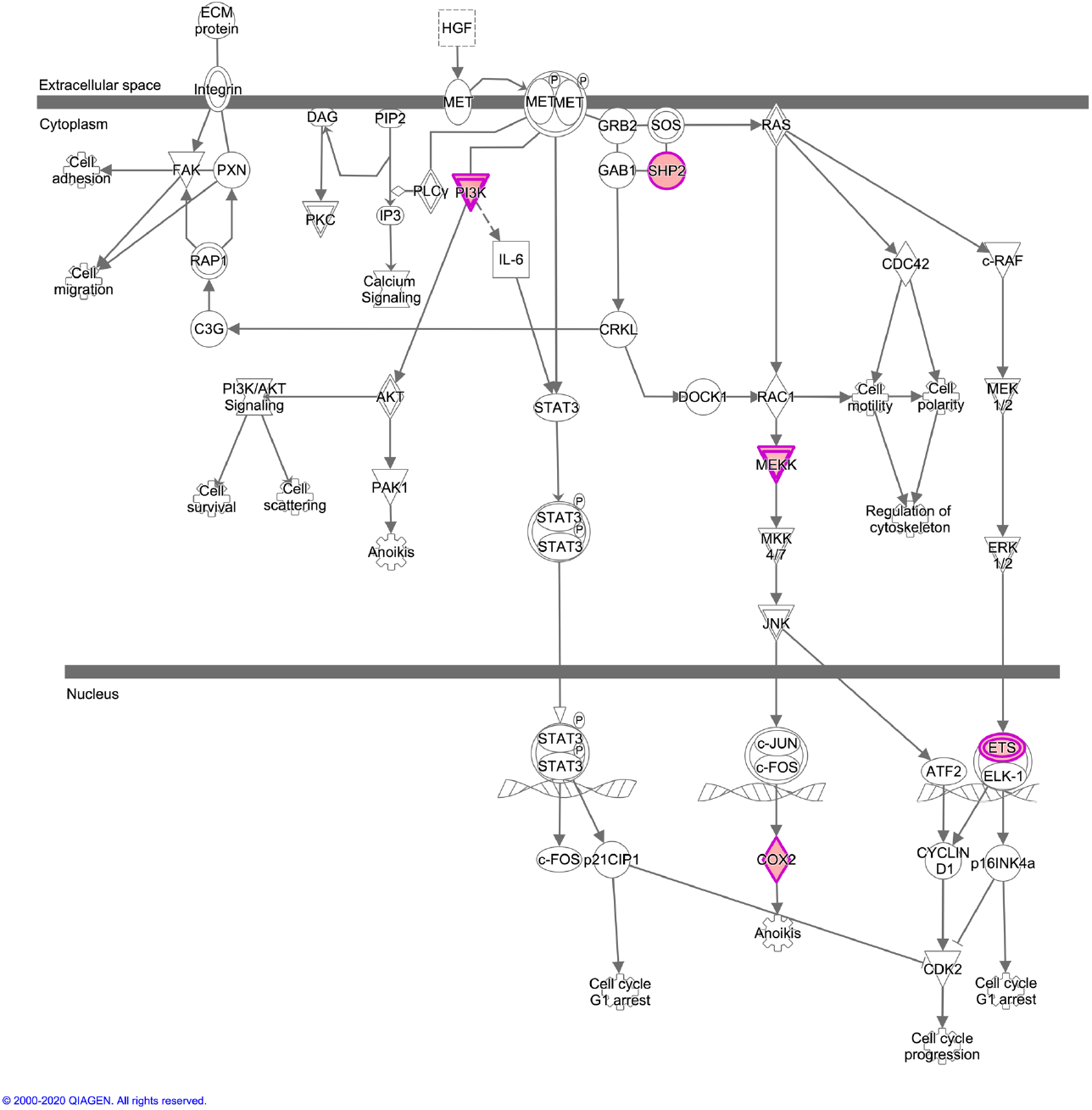
HGF signaling pathway map including the observed up-regulated genes (highlighted in pink).

**Figure S6c.**
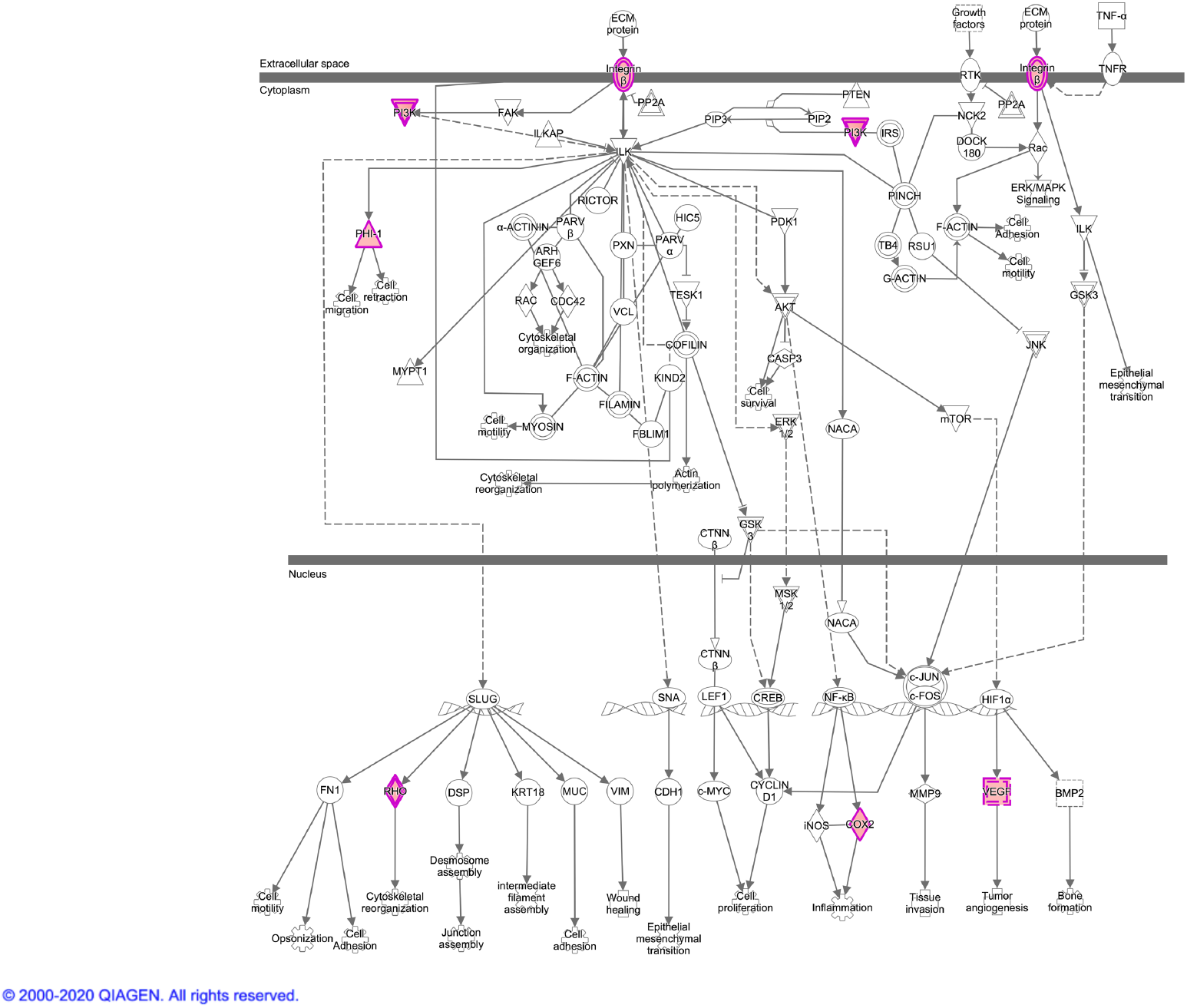
ILK signaling pathway map including the observed up-regulated genes (highlighted in pink).

**Figure S6d.**
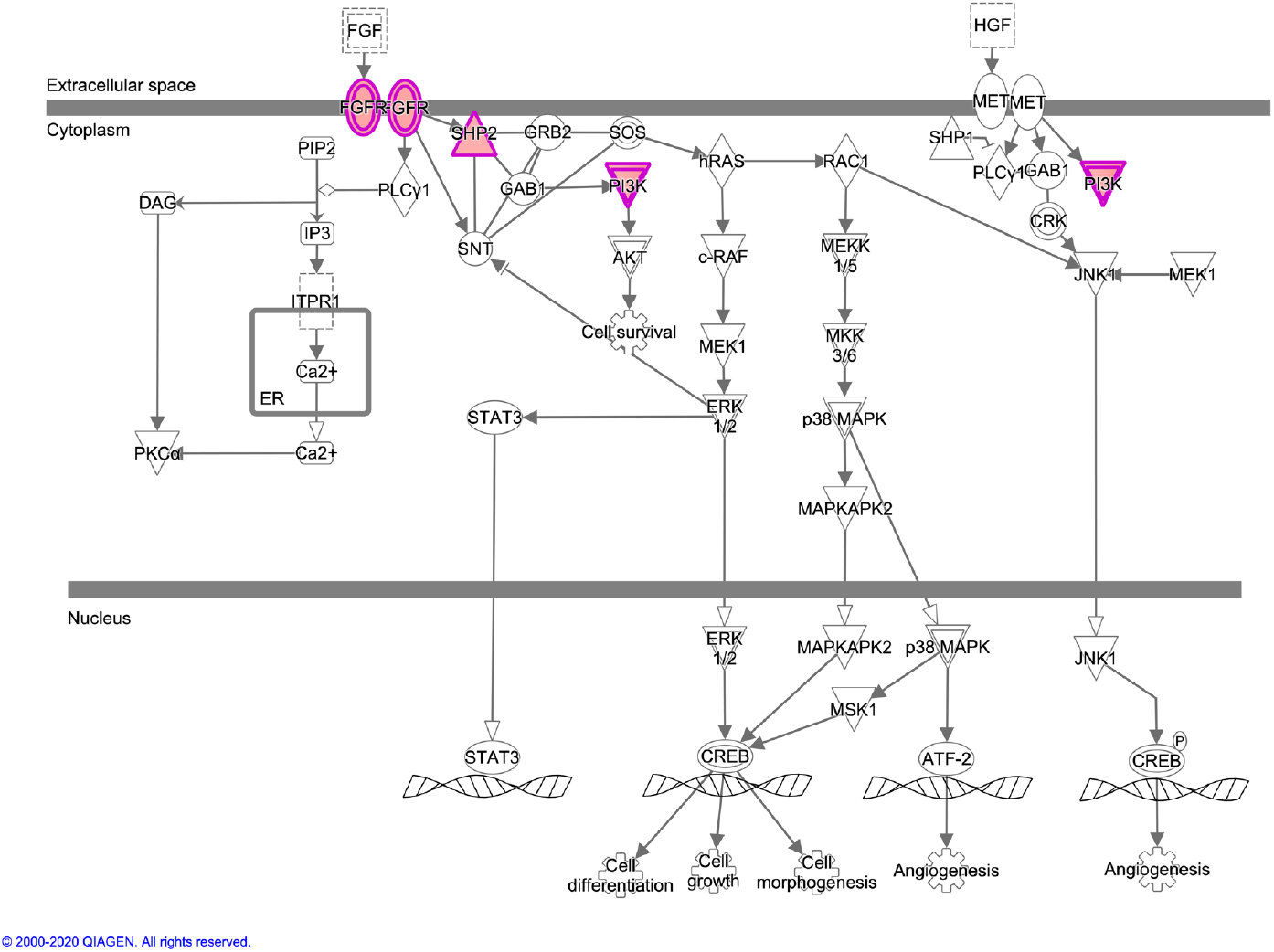
FGF signaling pathway map including the observed up-regulated genes (highlighted in pink).

**Figure S6e.**
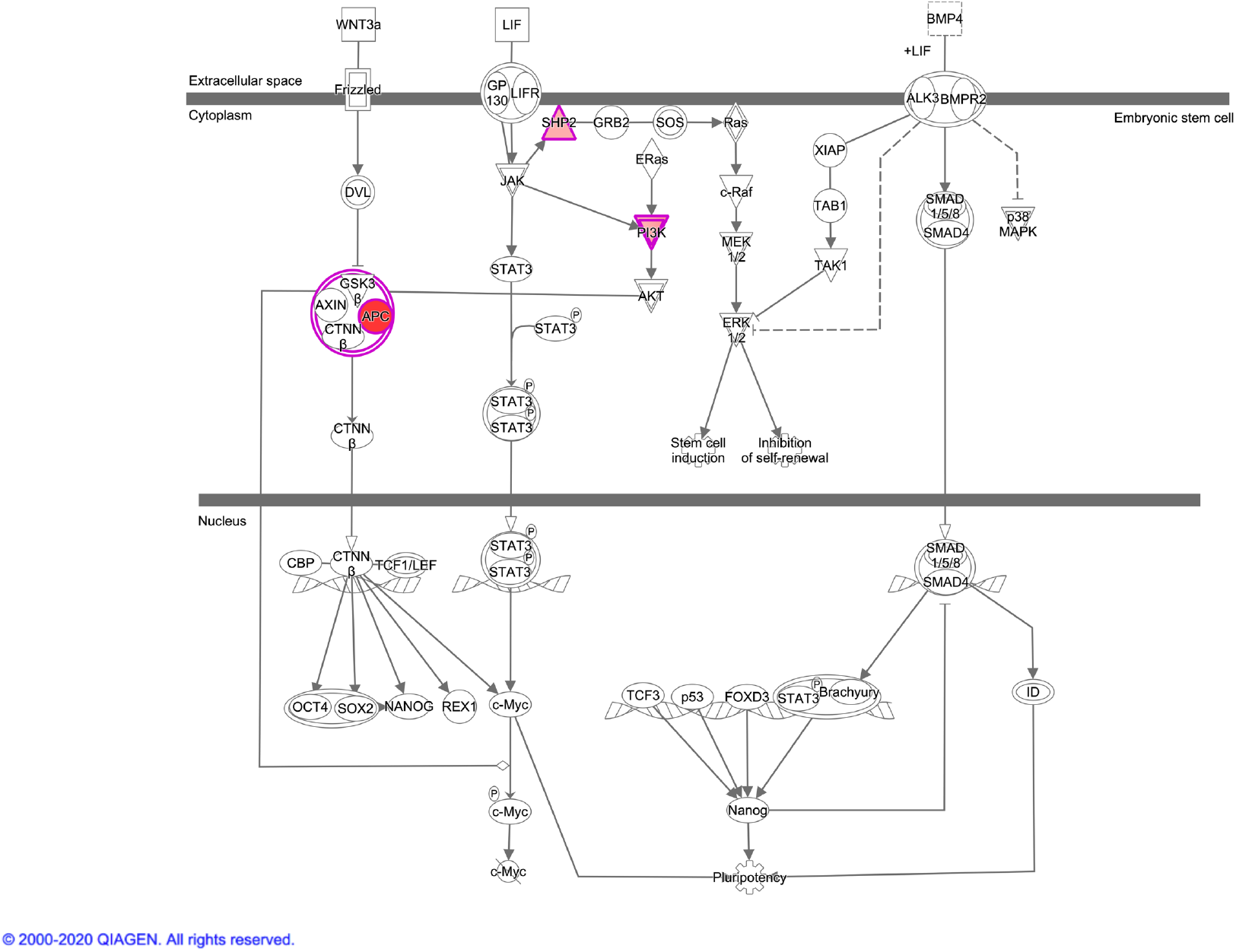
Mouse embryonic stem cell pluripotency signaling pathway map including the observed up-regulated genes (highlighted in pink).

**Figure S7a.**
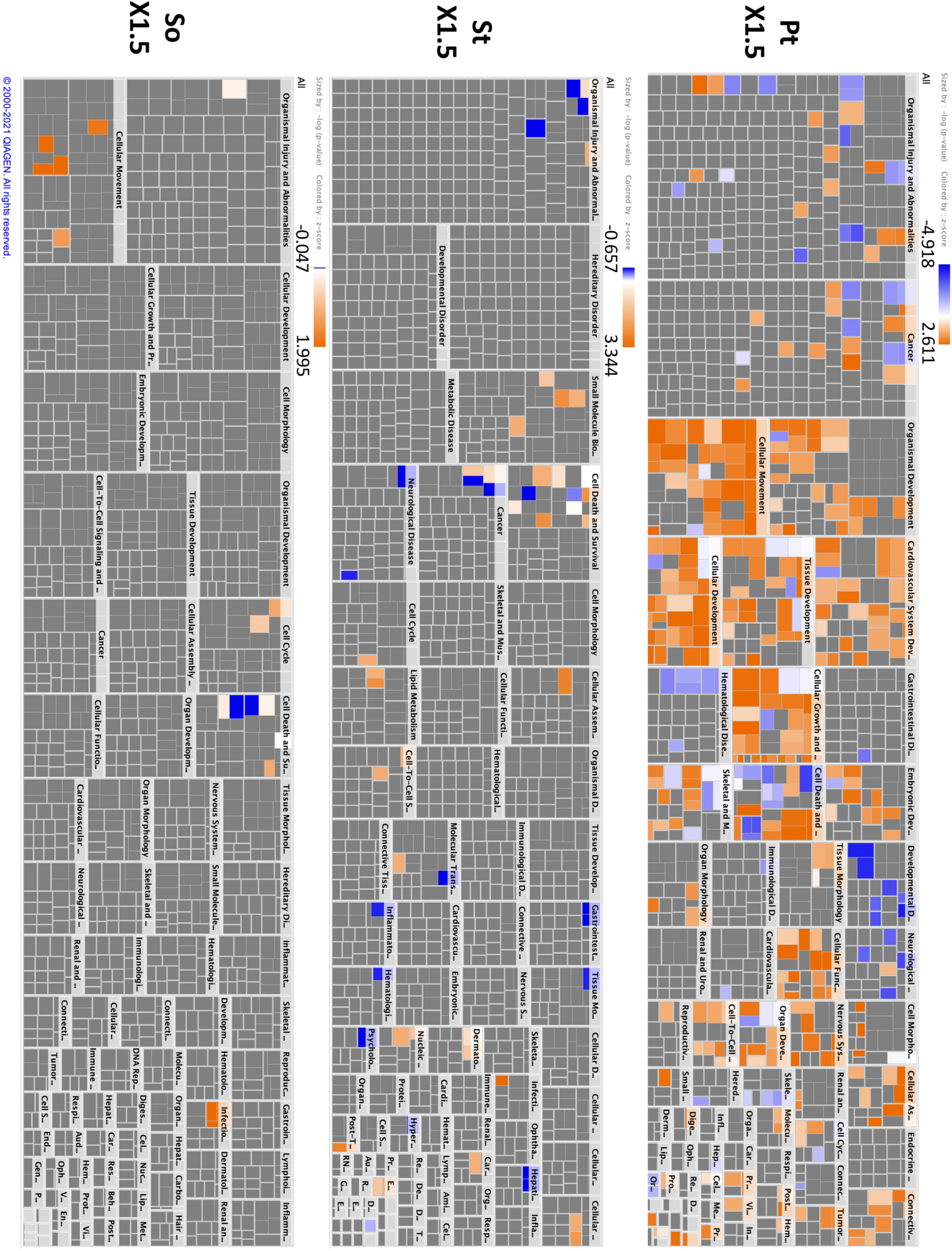
Diagrams of biofunctions predicted to be activated or inactivated based on each up-regulated expression profile (>1.5-fold) in the DNA microarray data observed in cells cultured on Pt, St, and So gels.

**Figure S7b.**
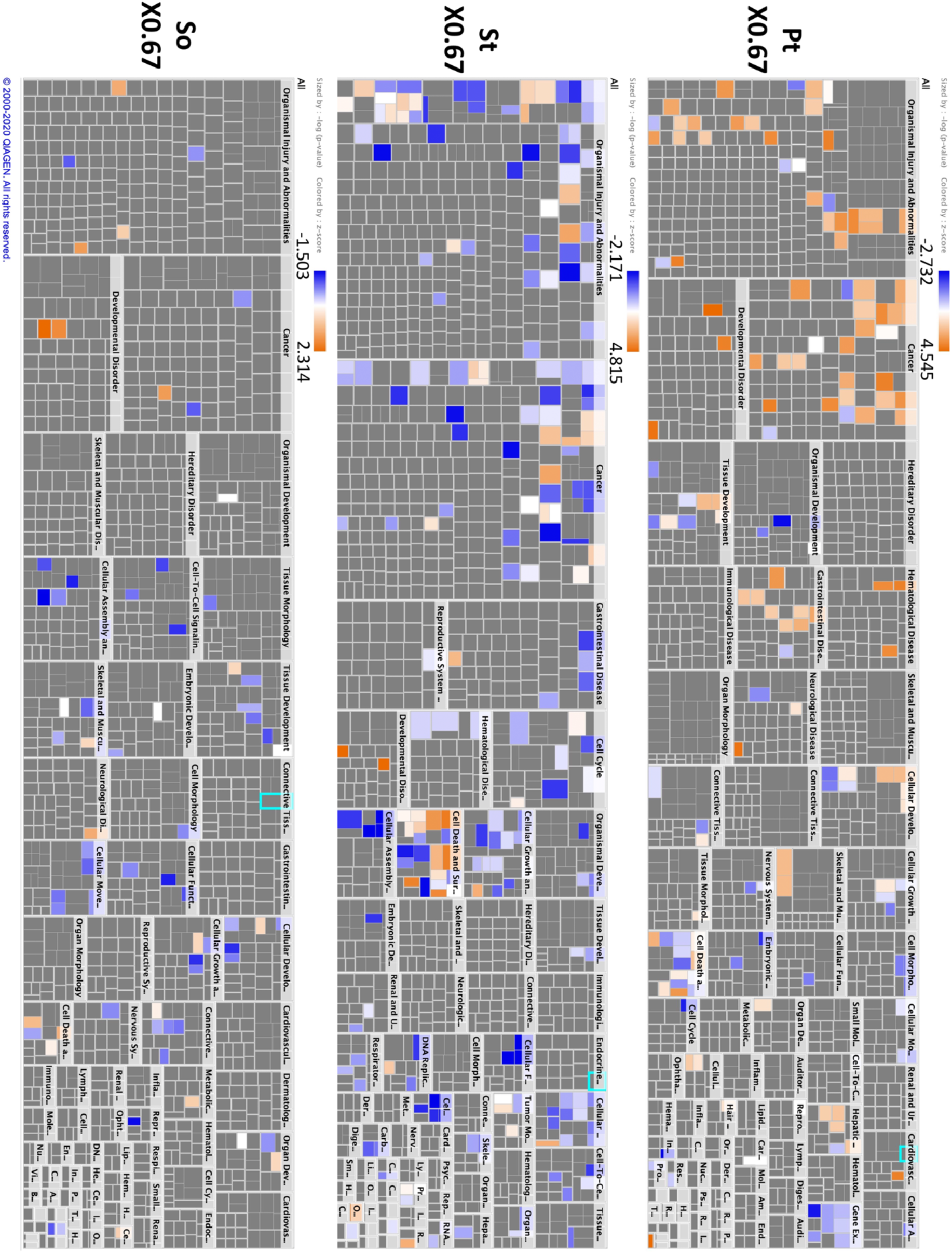
Diagrams of biofunctions predicted to be activated or inactivated based on each down-regulated expression profile (<0.67-fold) in the DNA microarray data observed in cells cultured on Pt, St, and So gels.

**Figure S8a.**
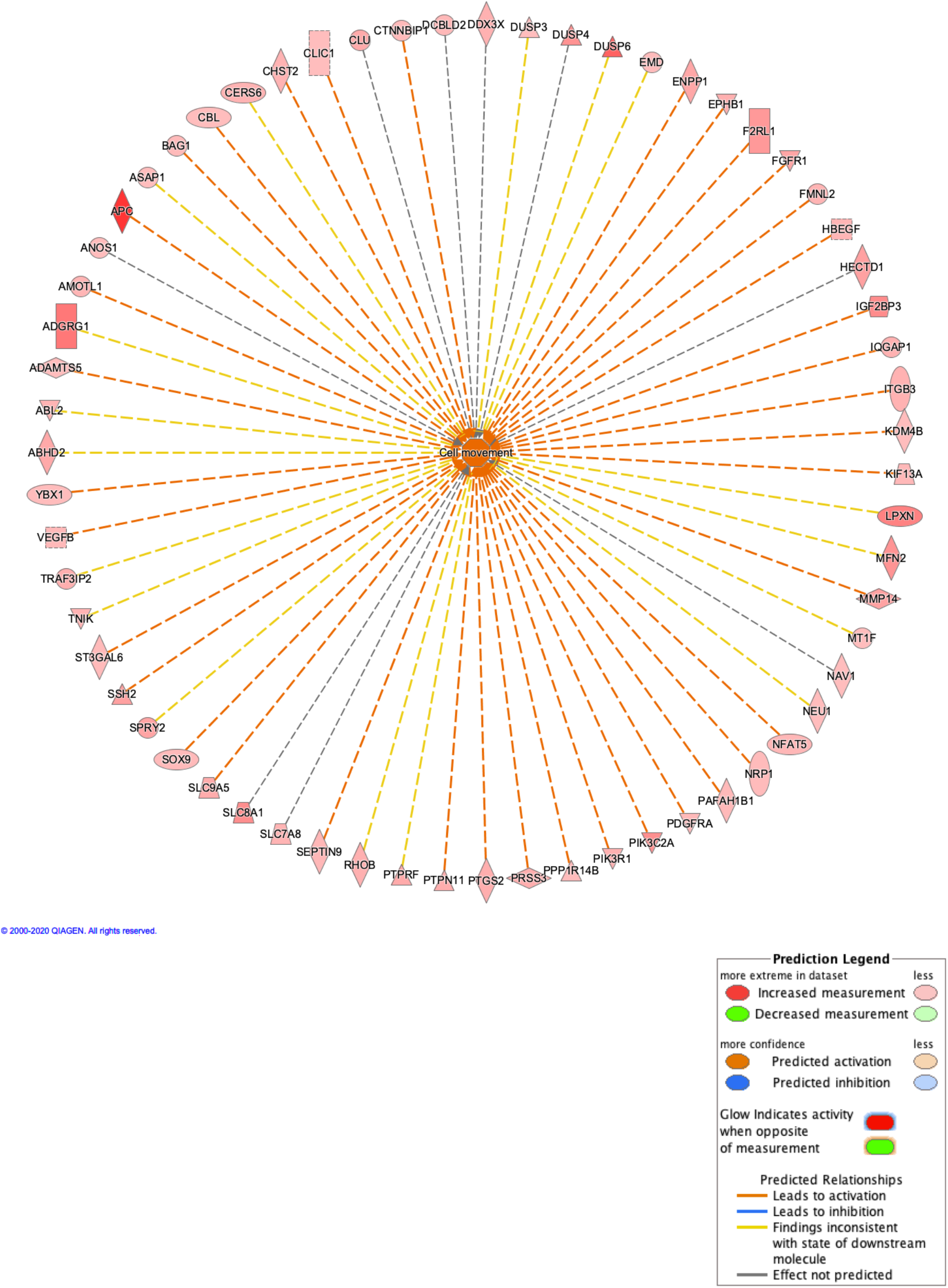
Network map of the Pt-specifically modulated genes related to the functional category of “Cell movement”.

**Figure S8b.**
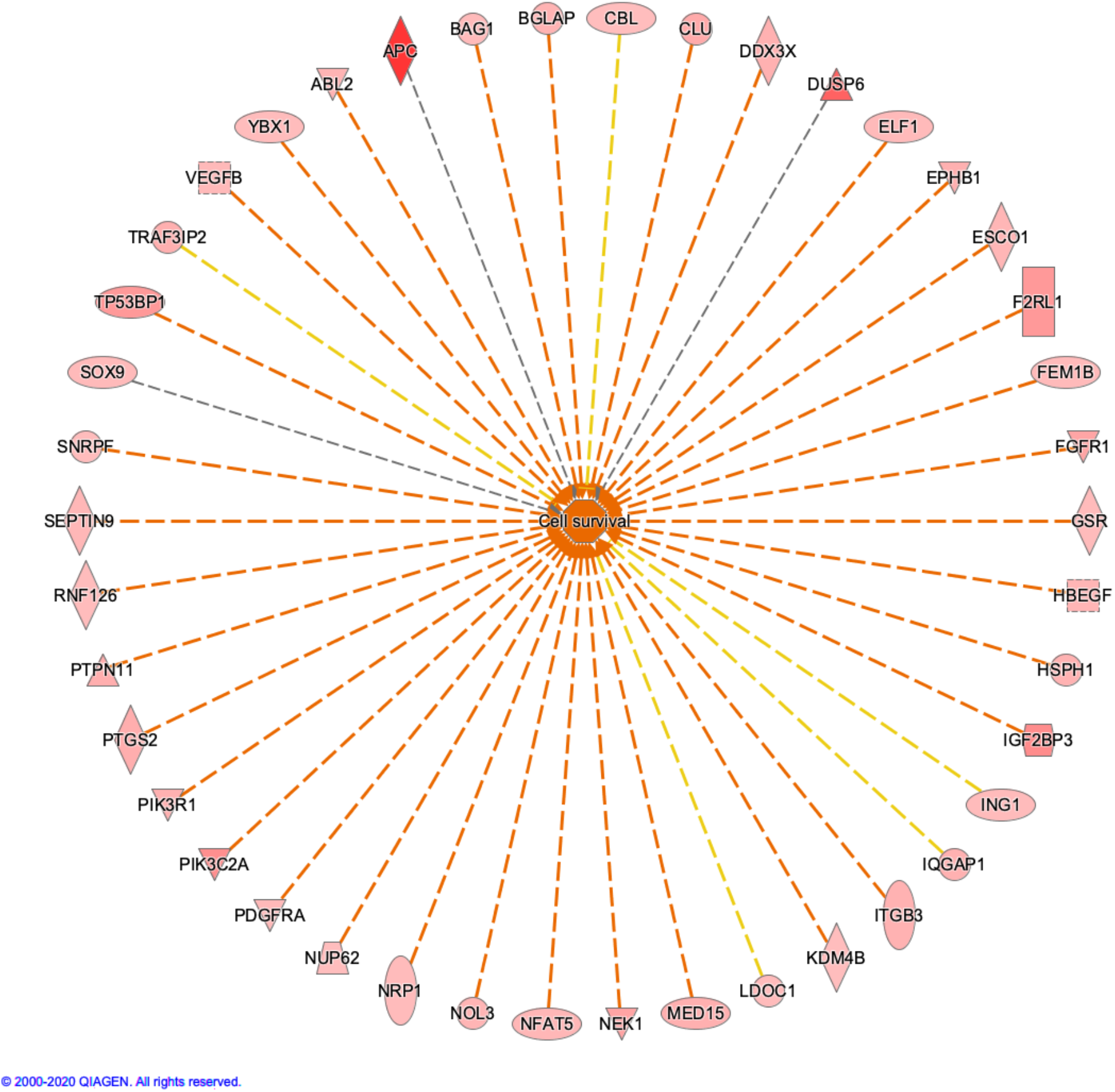
Network map of the Pt-specifically modulated genes related to the functional category of “Cell survival”.

**Figure S8c.**
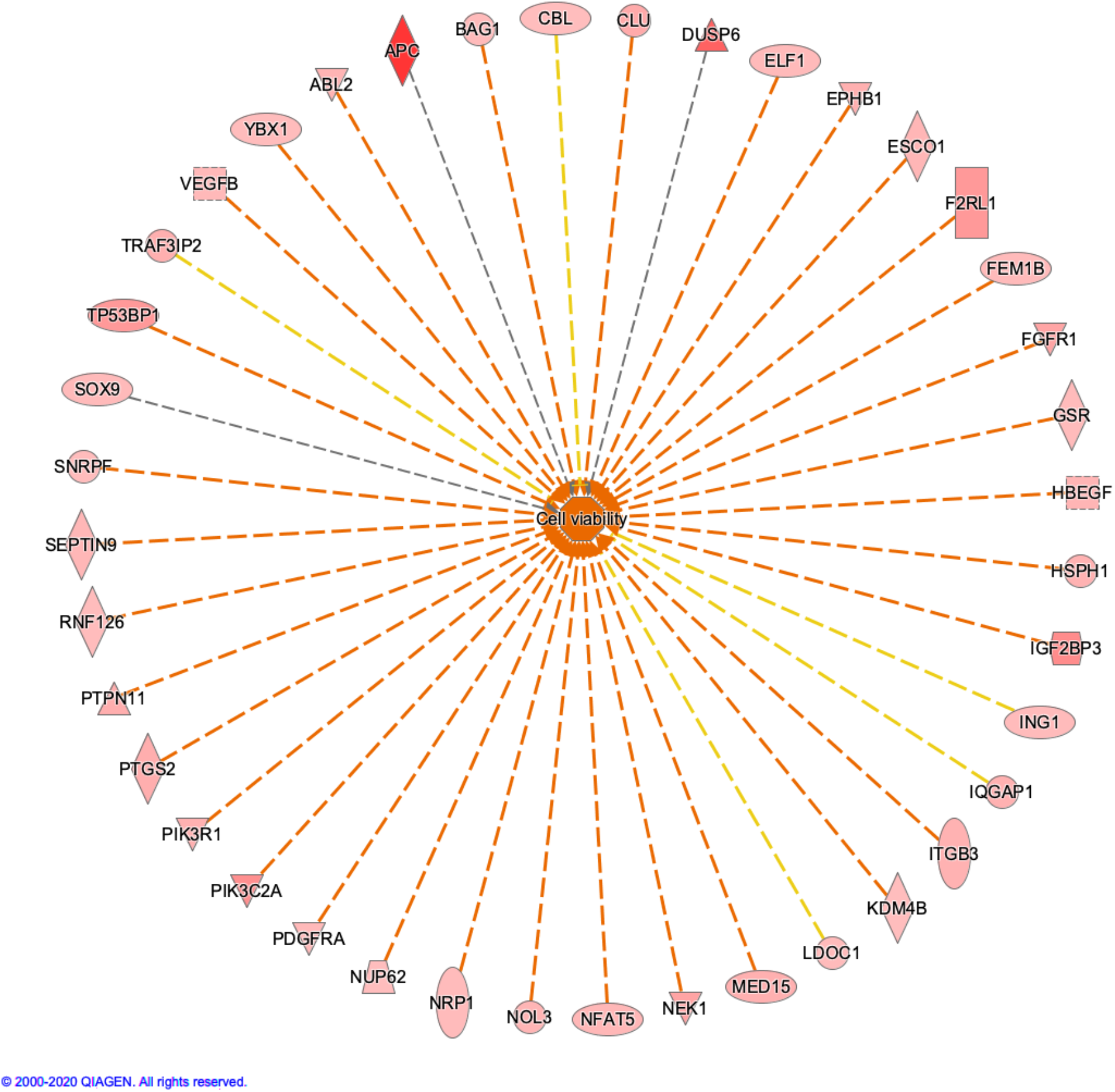
Network map of the Pt-specifically modulated genes related to the functional category of “Cell viability”.

**Figure S8d.**
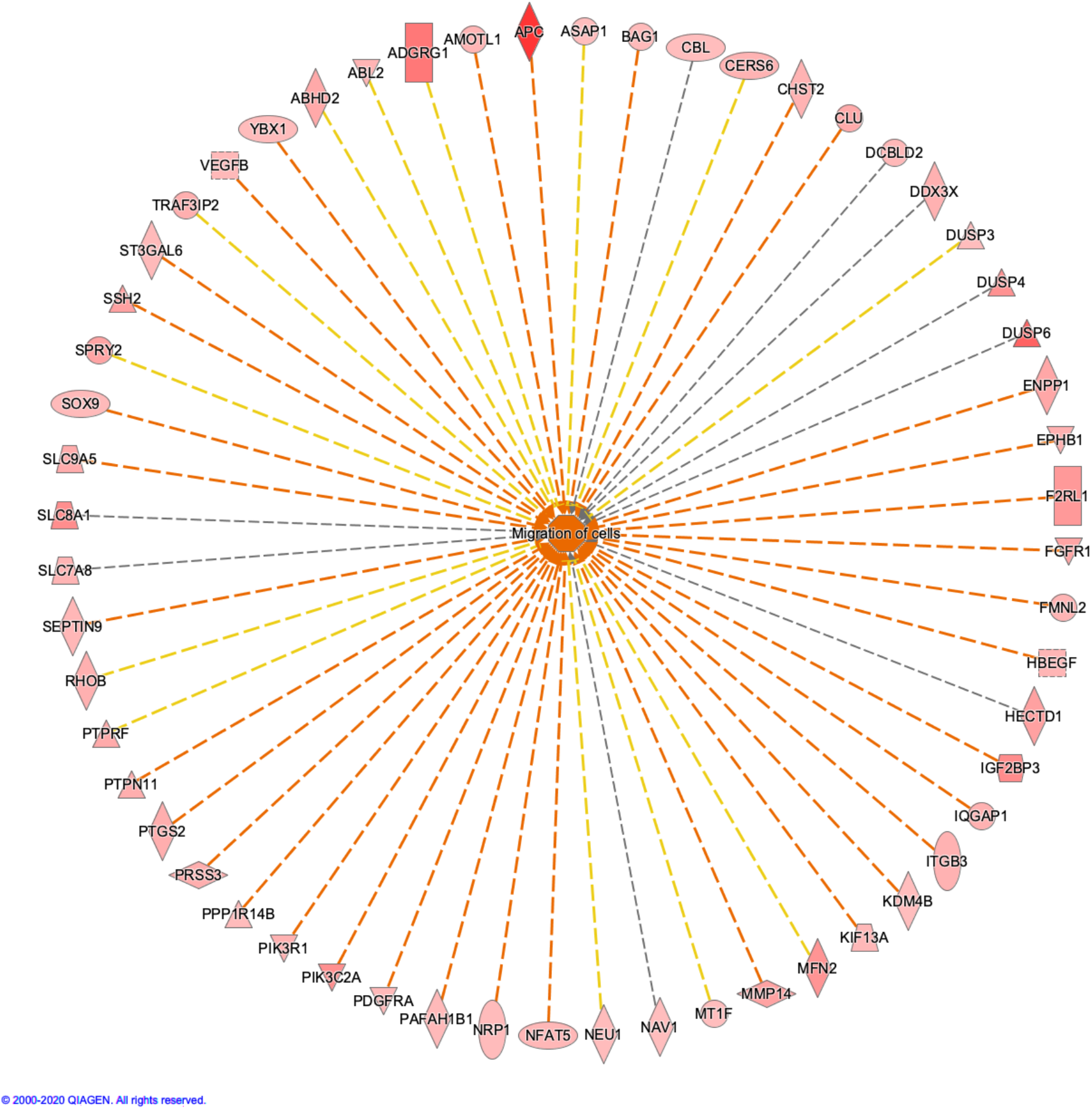
Network map of the Pt-specifically modulated genes related to the functional category of “Migration of cells”.

**Figure S8e.**
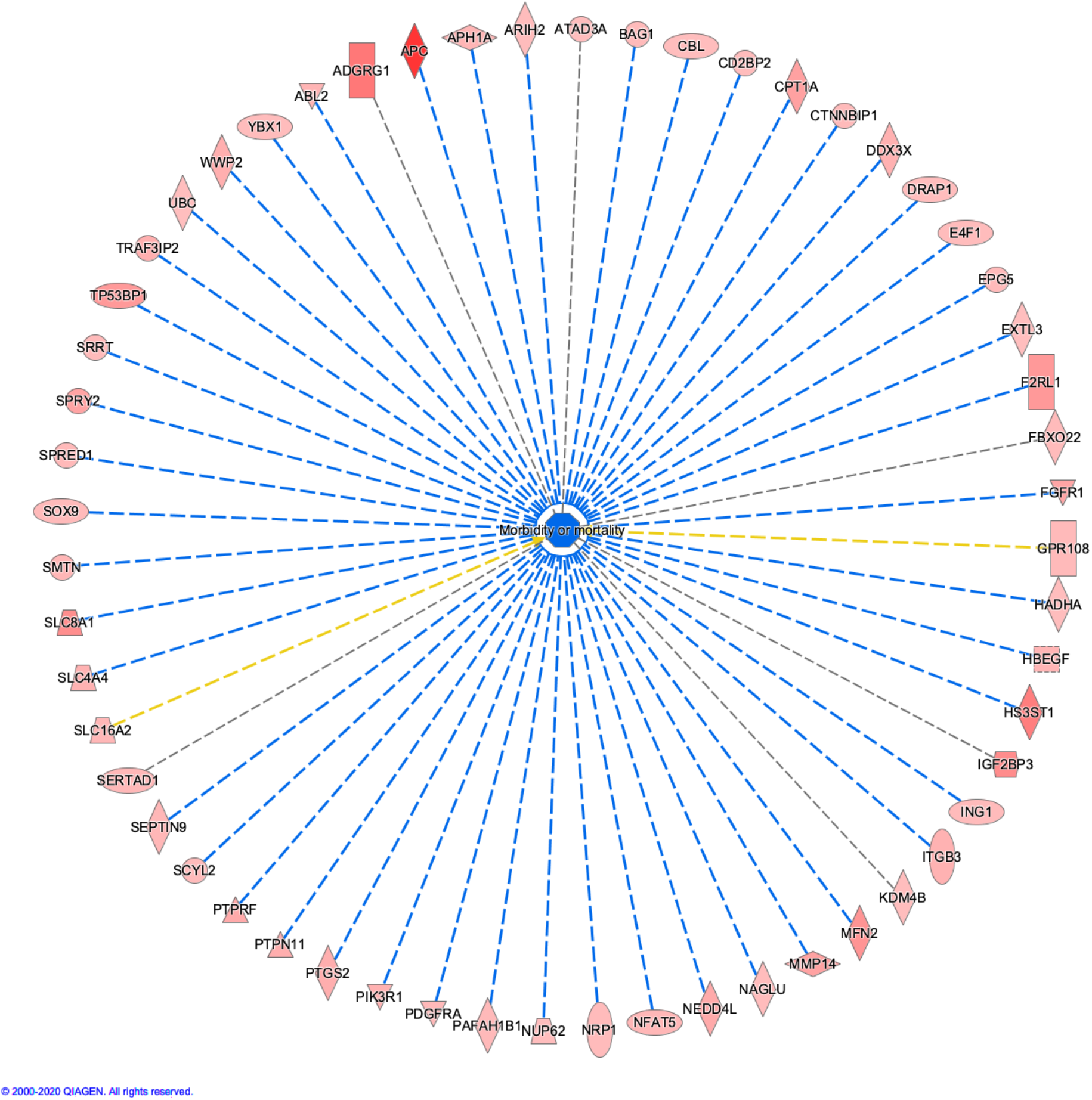
Network map of the Pt-specifically modulated genes related to the functional category of “Morbidity or Mortality”.

**Figure S8f.**
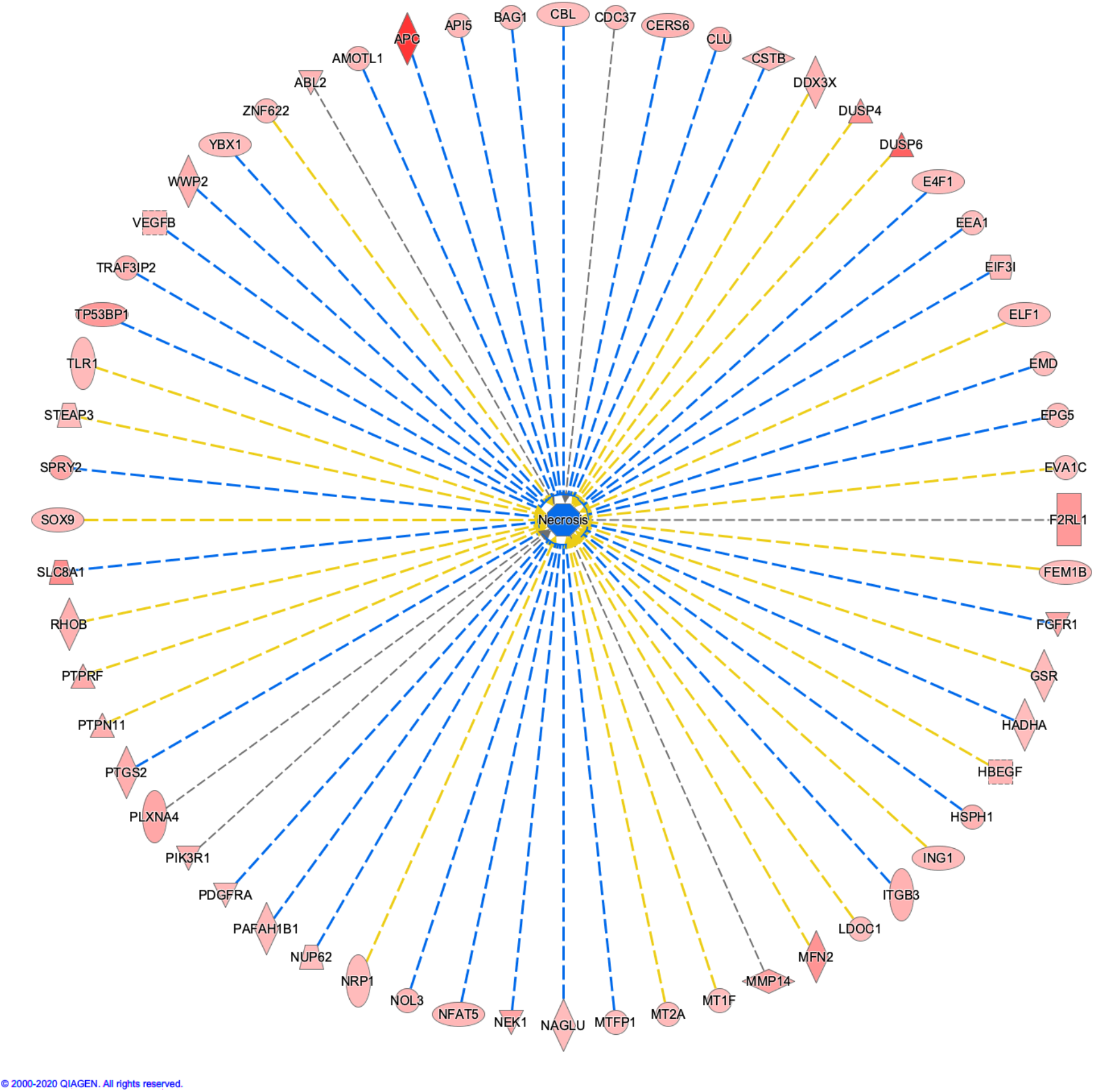
Network map of the Pt-specifically modulated genes related to the functional category of “Necrosis”.

**Figure S8g.**
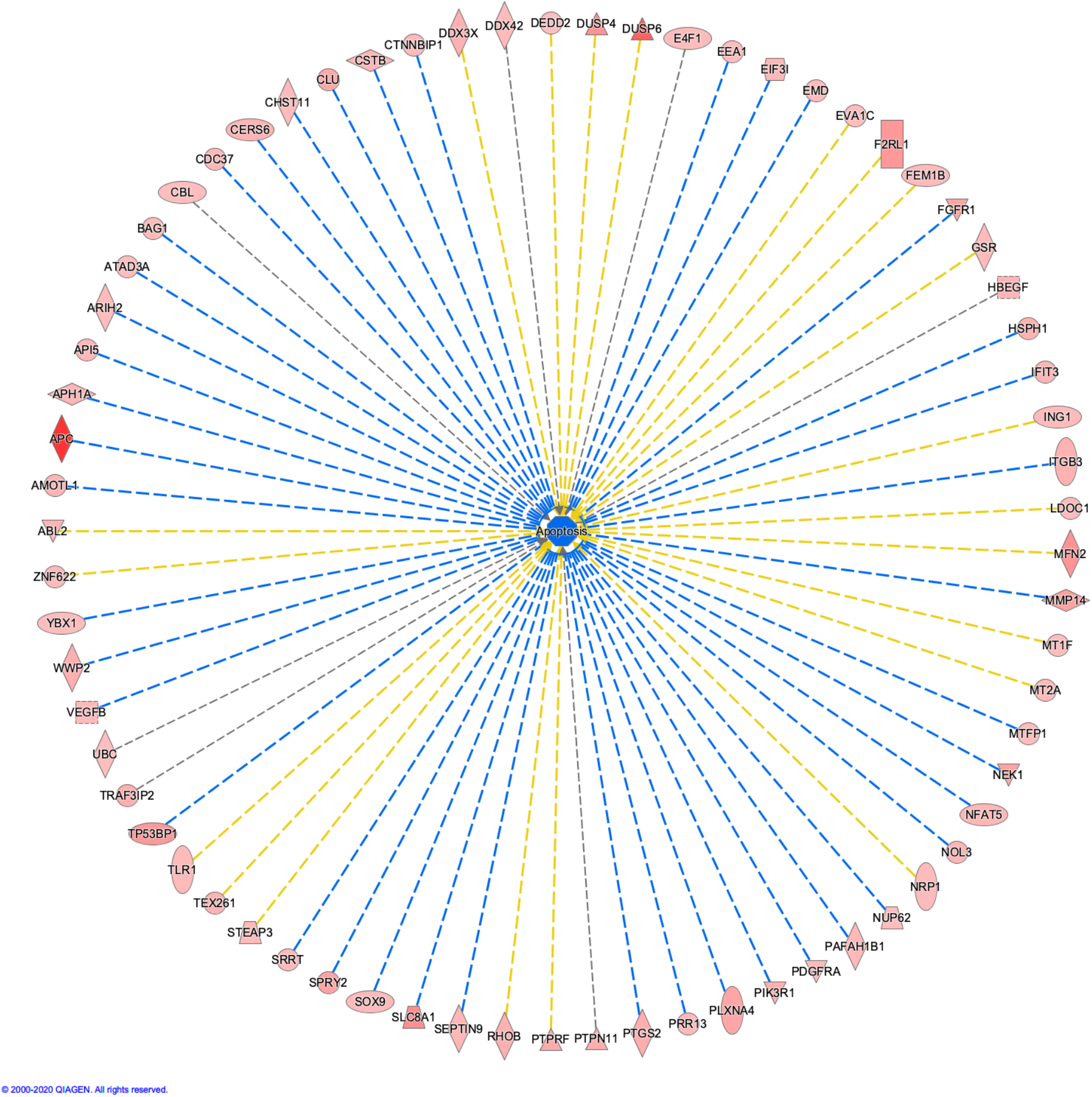
Network map of the Pt-specifically modulated genes related to the functional category of “Apoptosis”.

**Figure S8h.**
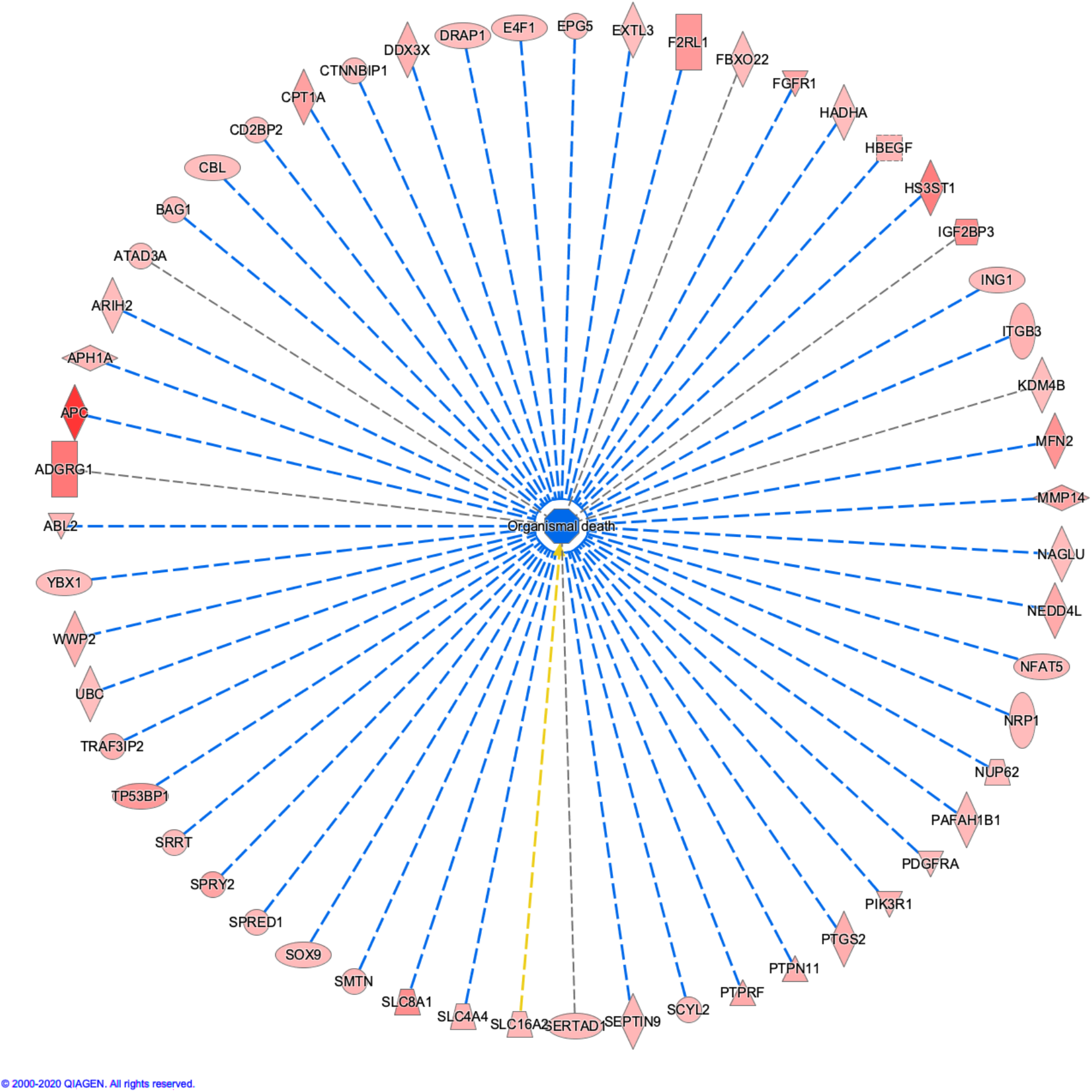
Network map of the Pt-specifically modulated genes related to the functional category of “Organismal death”.

**Figure S8i.**
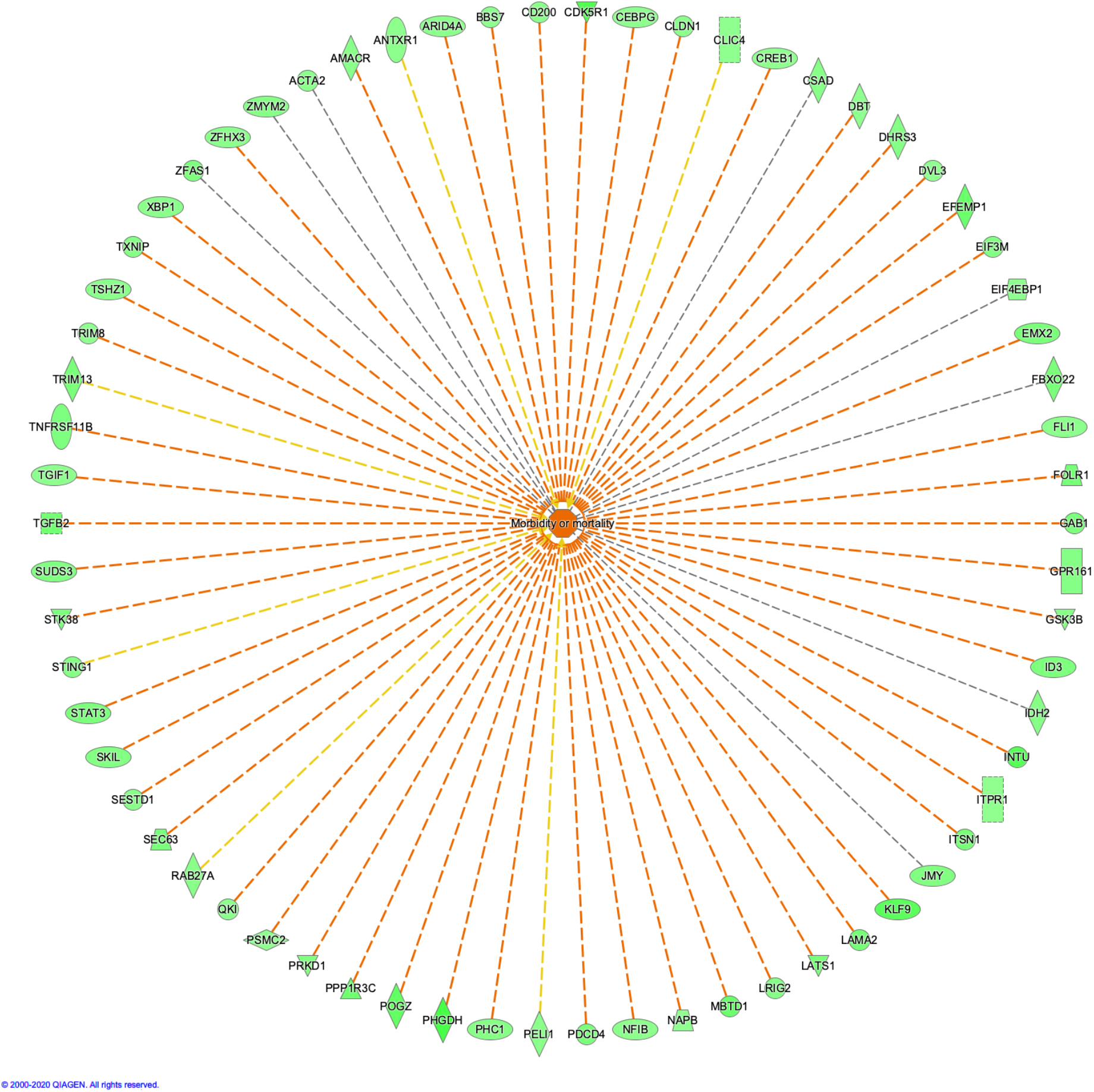
Network map of the Pt-specifically modulated genes related to the functional category of “Morbidity or Mortality”.

**Figure S9.**
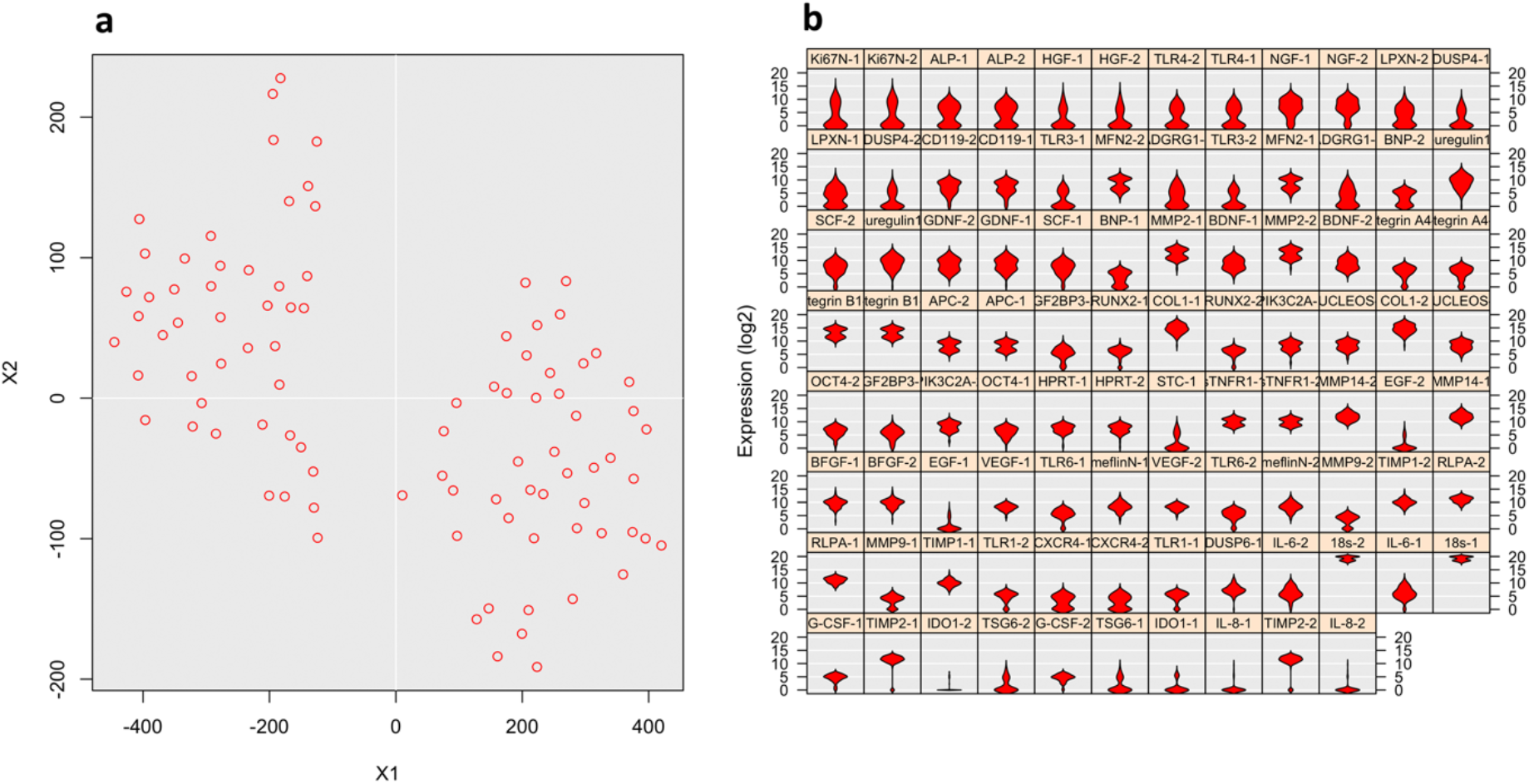
Single-cell population analysis of gene expression in cells cultured on Pt gels. (a) tSNE plot. (b) Violin plot.

**Figure S10.**
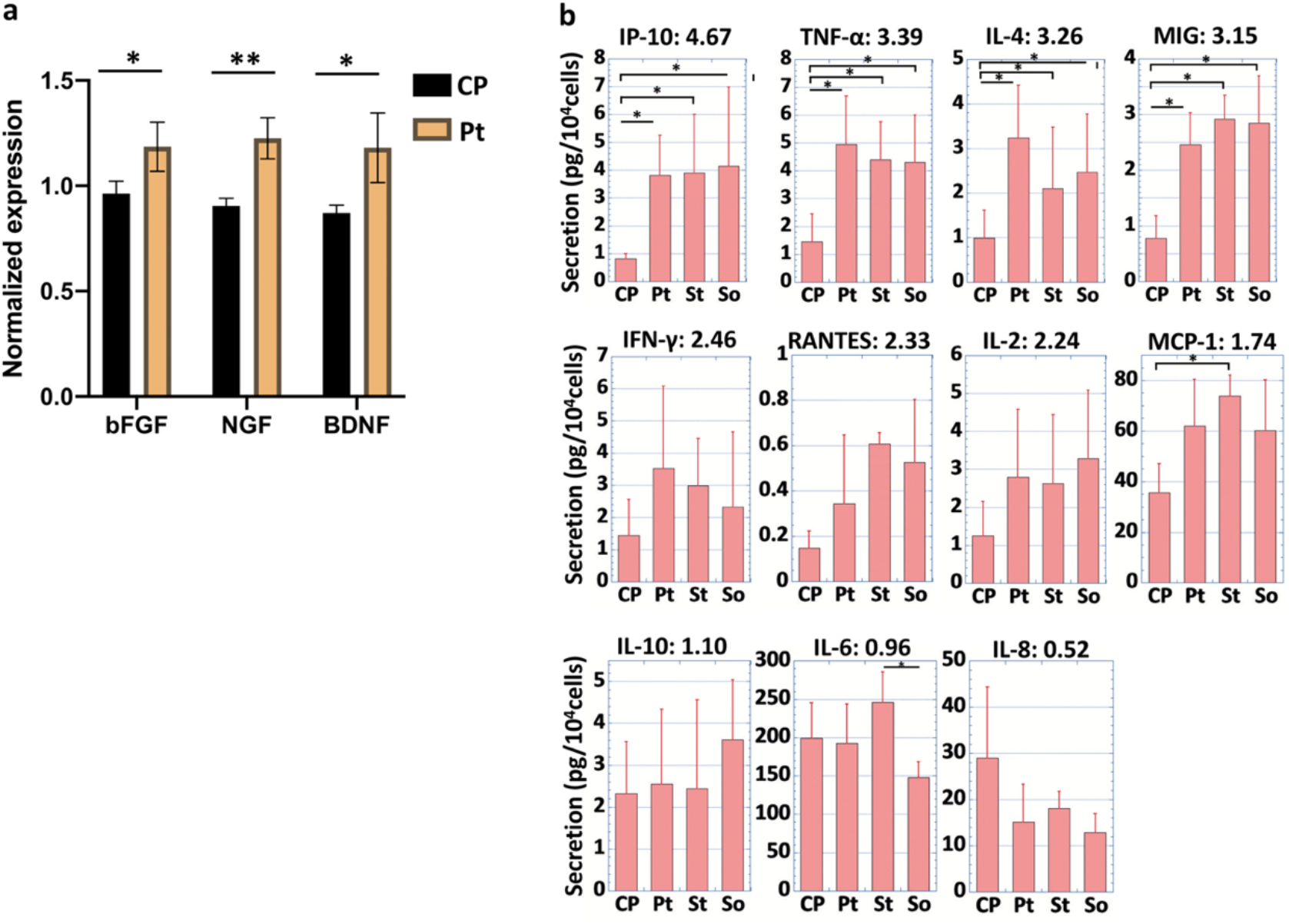
Expression analysis of therapeutic-related molecules in dual-durotactic MSCs cultured on Pt gels. **(a)** Gene expression of bFGF, NGF, and BDNF measured with q-PCR. **(b)** Secretion of cytokines quantified with a microbead array and flow cytometer. The values on each graph show the ratio of the mean secretion level of Pt to that of control CP. ** P<0.01, * P<0.05.

**Figure S11.**
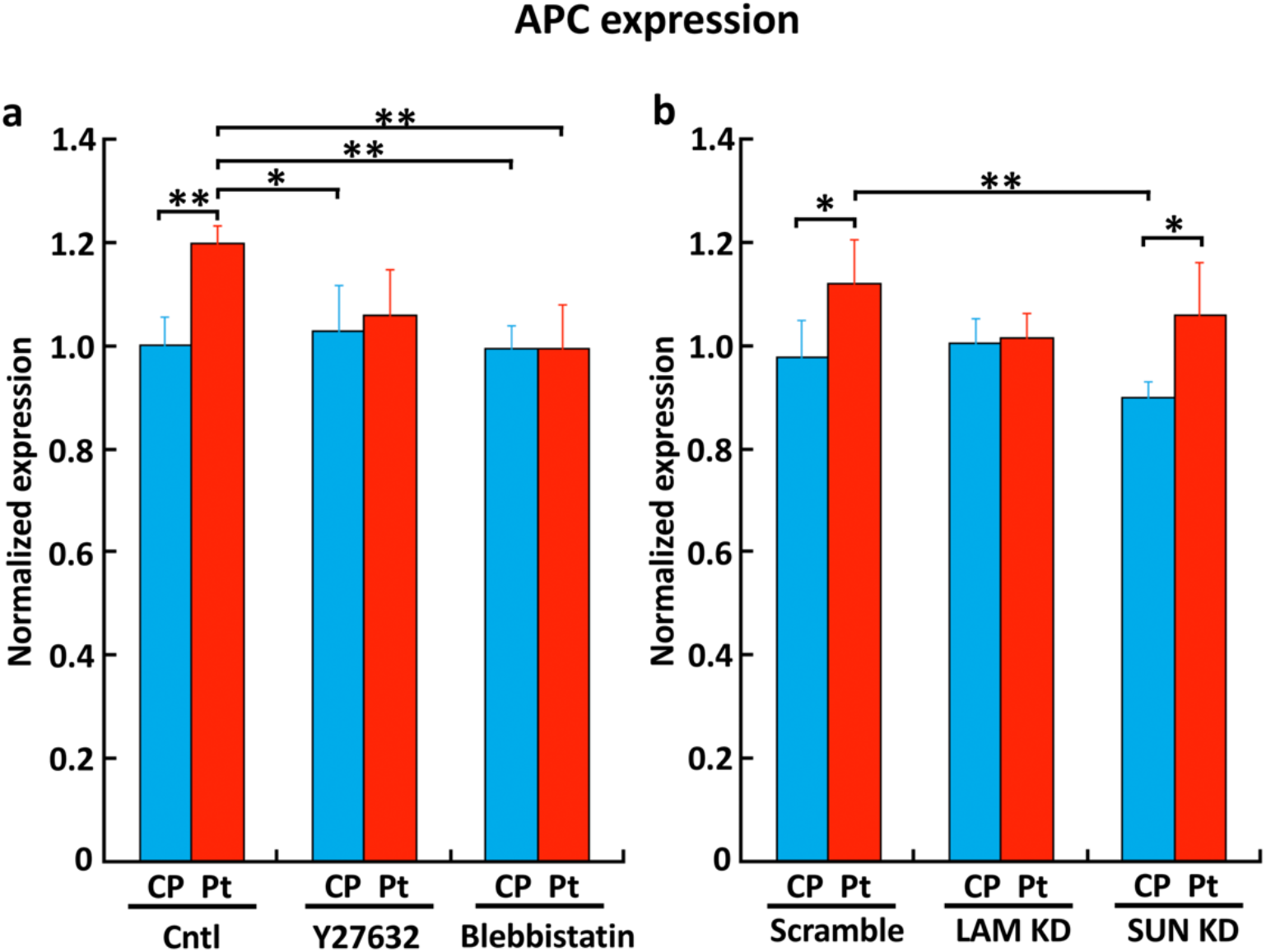
Effects of cytoskeletal tension inhibitors and knockdown of LINC-related molecules on the expression of APC gene. **(a)** APC expression in cells cultured on CP and Pt gels in the presence of Y27632 and blebbistatin measured with q-PCR. **(b)** APC expression in cells cultured on CP and Pt gels under knockdown with siRNA of LaminA/C and SUN1 measured with q-PCR. ** P<0.01, * P<0.05.

### Supplementary Movies

**SI Movie 1. Time-lapse observation of nomadic migration of dual-durotaxing MSCs on a microelastically-patterned gel with stiff triangles**

**SI Movie 2. Representative traction force dynamics in dual-durotaxing MSCs on the Pt gel**

**SI Movie 3. Representative traction force dynamics in MSCs moving on the St gel**

**SI Movie 4. Representative traction force dynamics in MSCs moving on the So gel**

**SI Movie 5. Live imaging of the cell nuclei in dual-durotaxing MSCs on the Pt gel**

**SI Movie 6. Live imaging of the cell nuclei in MSCs moving on CP**

**SI Movie 7. Representative live imaging of chromatin dynamics in a single cell nucleus in MSCs moving on the Pt gel. Scale bar: 10 μm**

**SI Movie 8. Representative live imaging of chromatin dynamics in a single cell nucleus in MSCs moving on CP. Scale bar: 10 μm**

**SI Movie 9. The most successful live-cell imaging of the moment of nucleocytoplasmic shuttling of YAP in a dual-durotaxing MSC that is crossing the elasticity boundary of a stiff triangle on Pt gel. By 6hr, YAP is mainly localized in cytoplasm inside the stiff triangle pattern though it is usually localized in the nucleus on the plain stiff gel, which is a typical response on Pt gel due to the time-lag effect of YAP translocation when the cell detects the stiff region. YAP translocation into the nucleus requires several hours after the cell enters the stiff region, while translocation into the cytoplasm also needs several hours after the cell feels the soft region. The dual-durotaxing cell moves over the cell-size-scale stiff region about 4hr in the present patterning, therefore the time-lag of YAP translocation emerges. The cytoplasmic localization of YAP in the stiff triangle region is the delayed response of translocation when the cell felt the soft surroundings. Actually, after 6hr, when the cell crosses the elasticity boundary around the corner of the stiff triangle from the stiff to soft region, YAP starts to translocate from the cytoplasm to the nucleus even though the cell starts to feel the soft region. This is because the cell sensed the stiff region.**

### Supplementary Tables

**Table S1.**
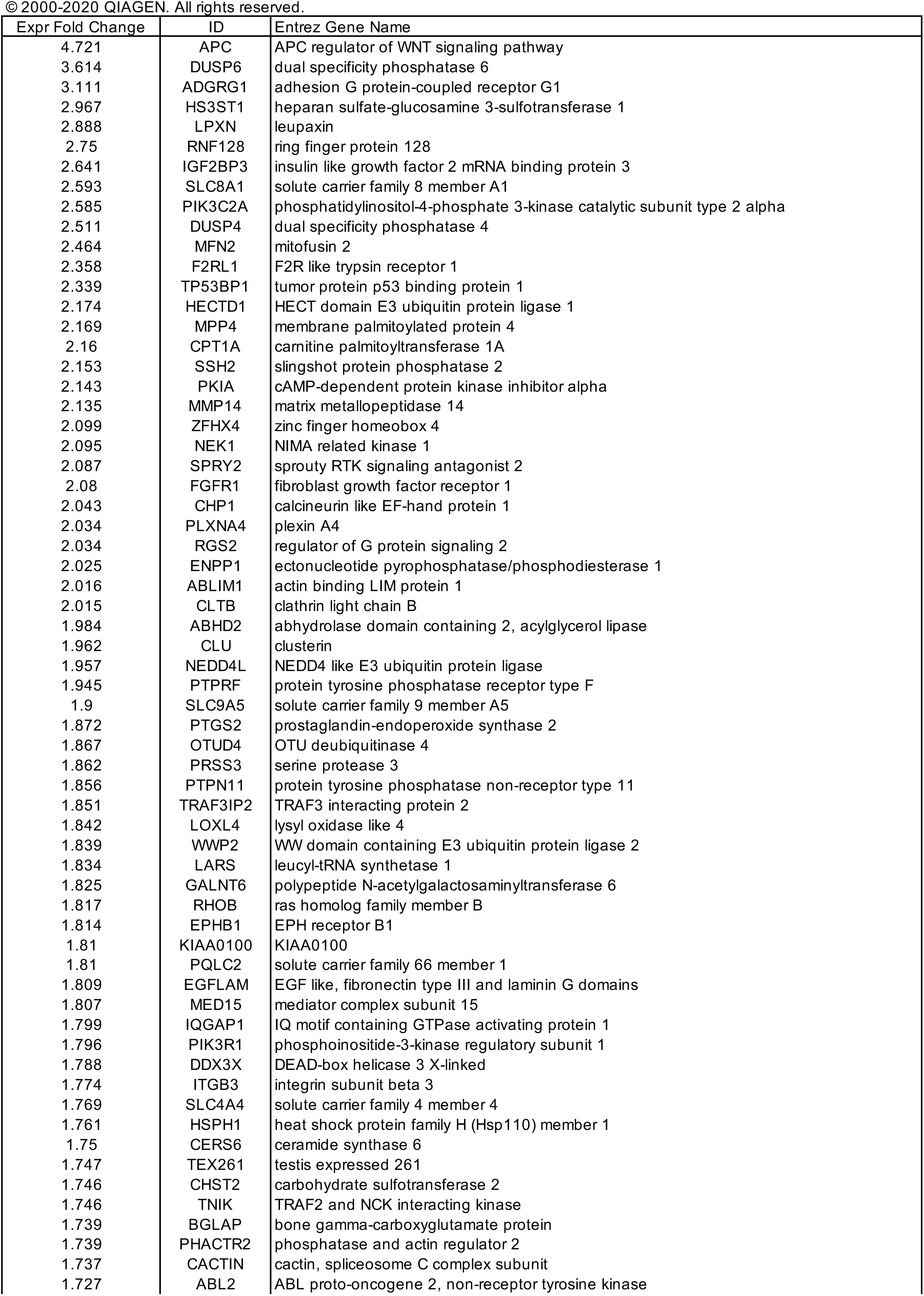
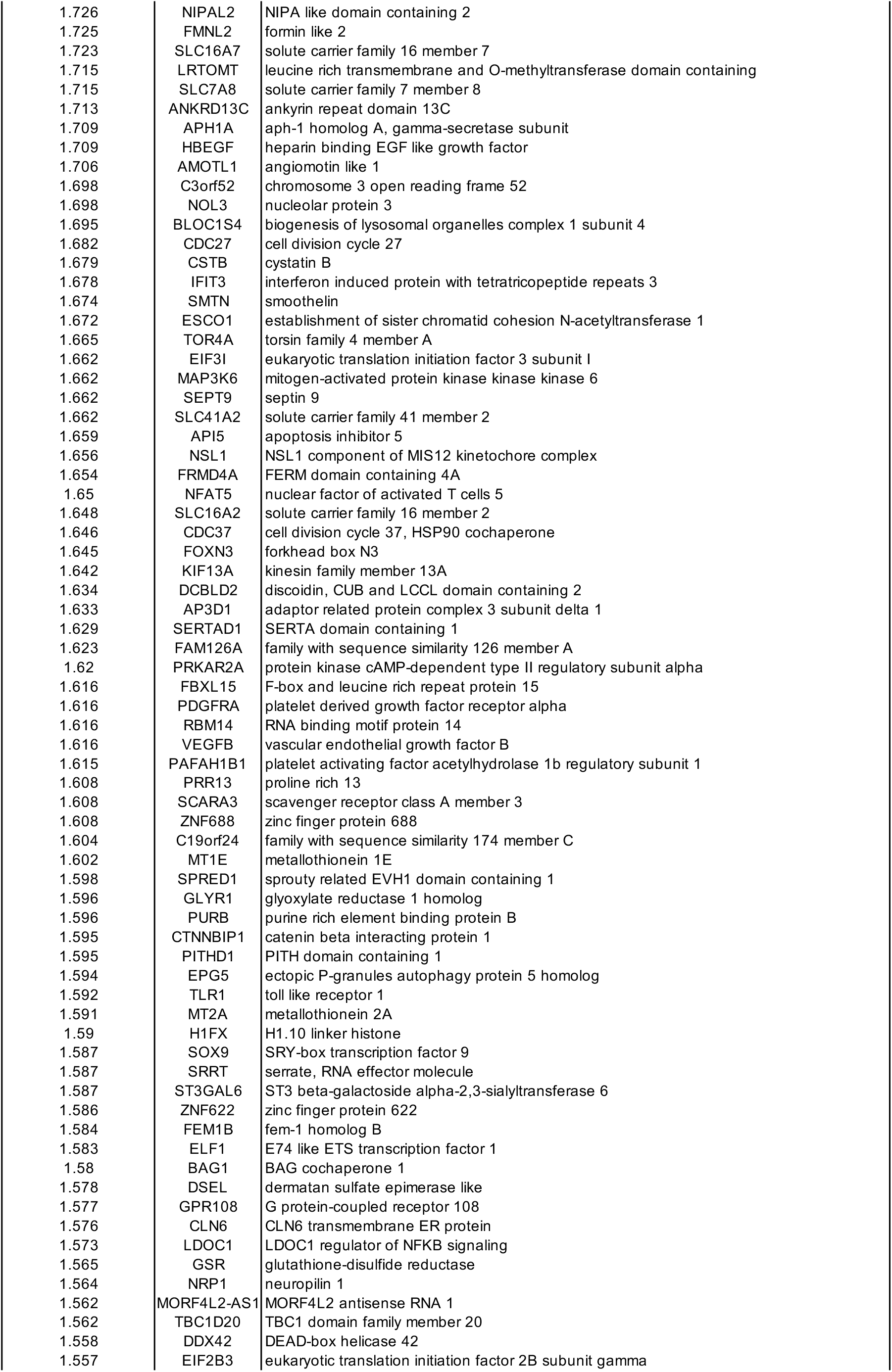
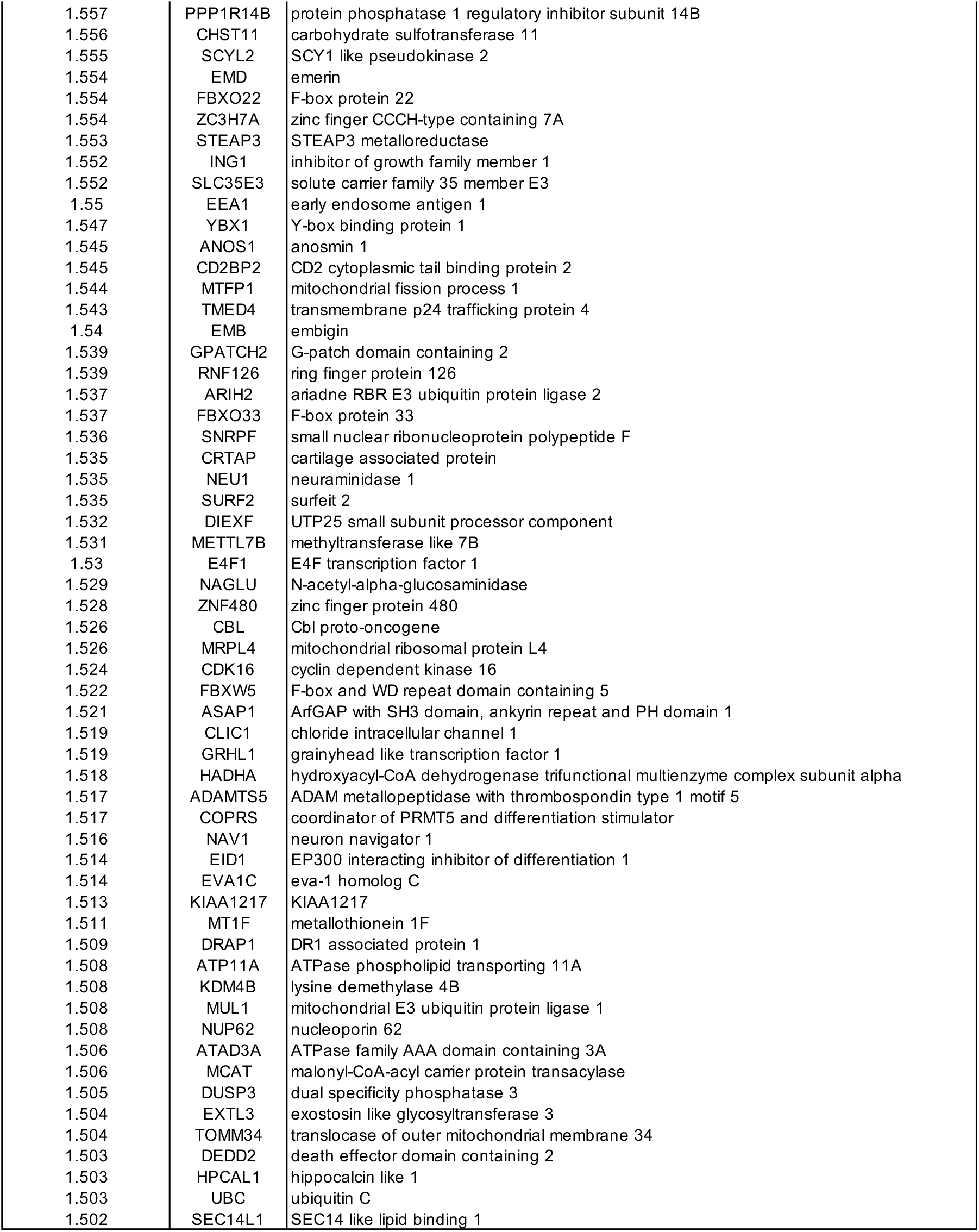
Pt-specifc up-regulated genes (>1.5-fold)

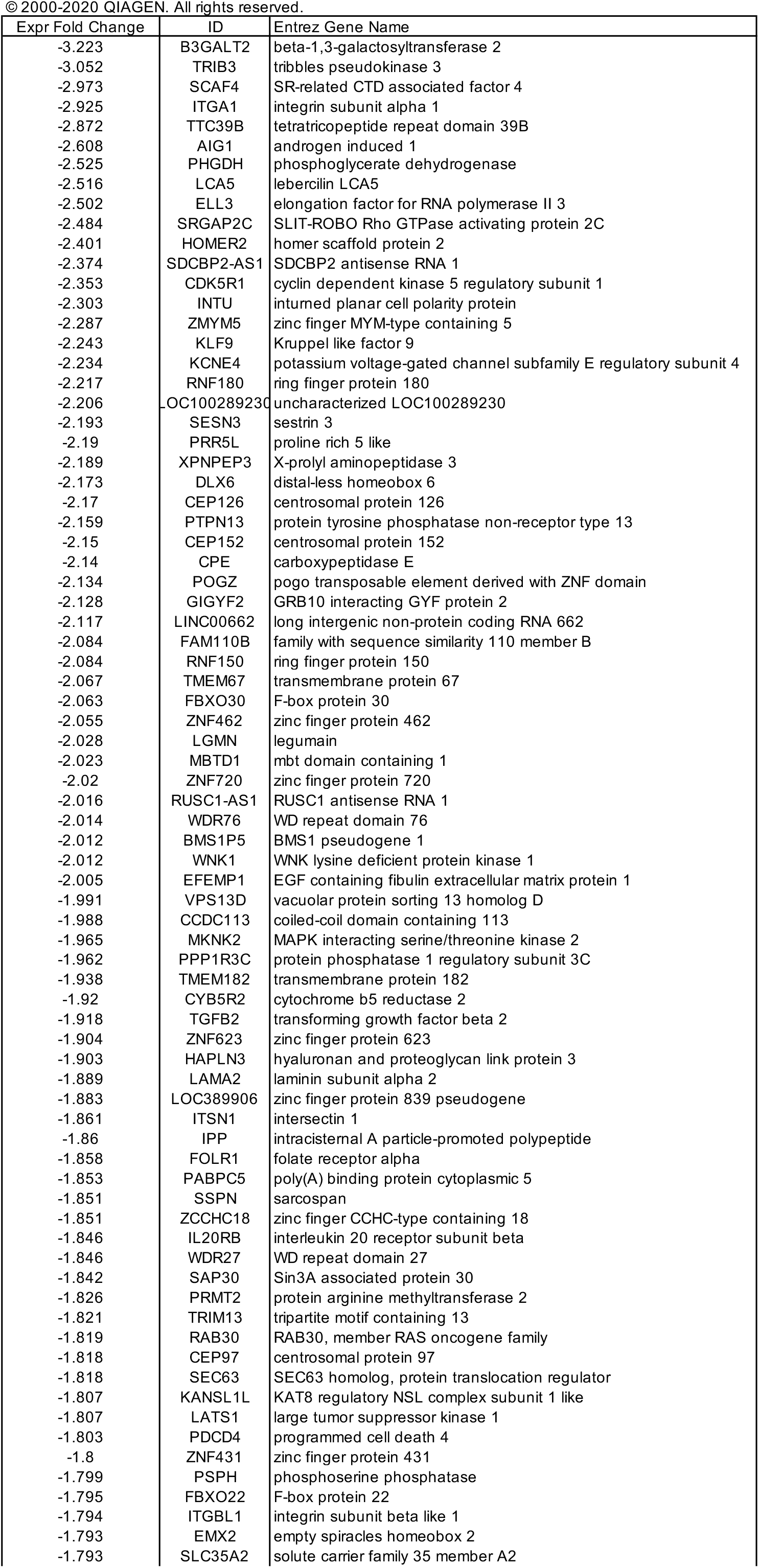
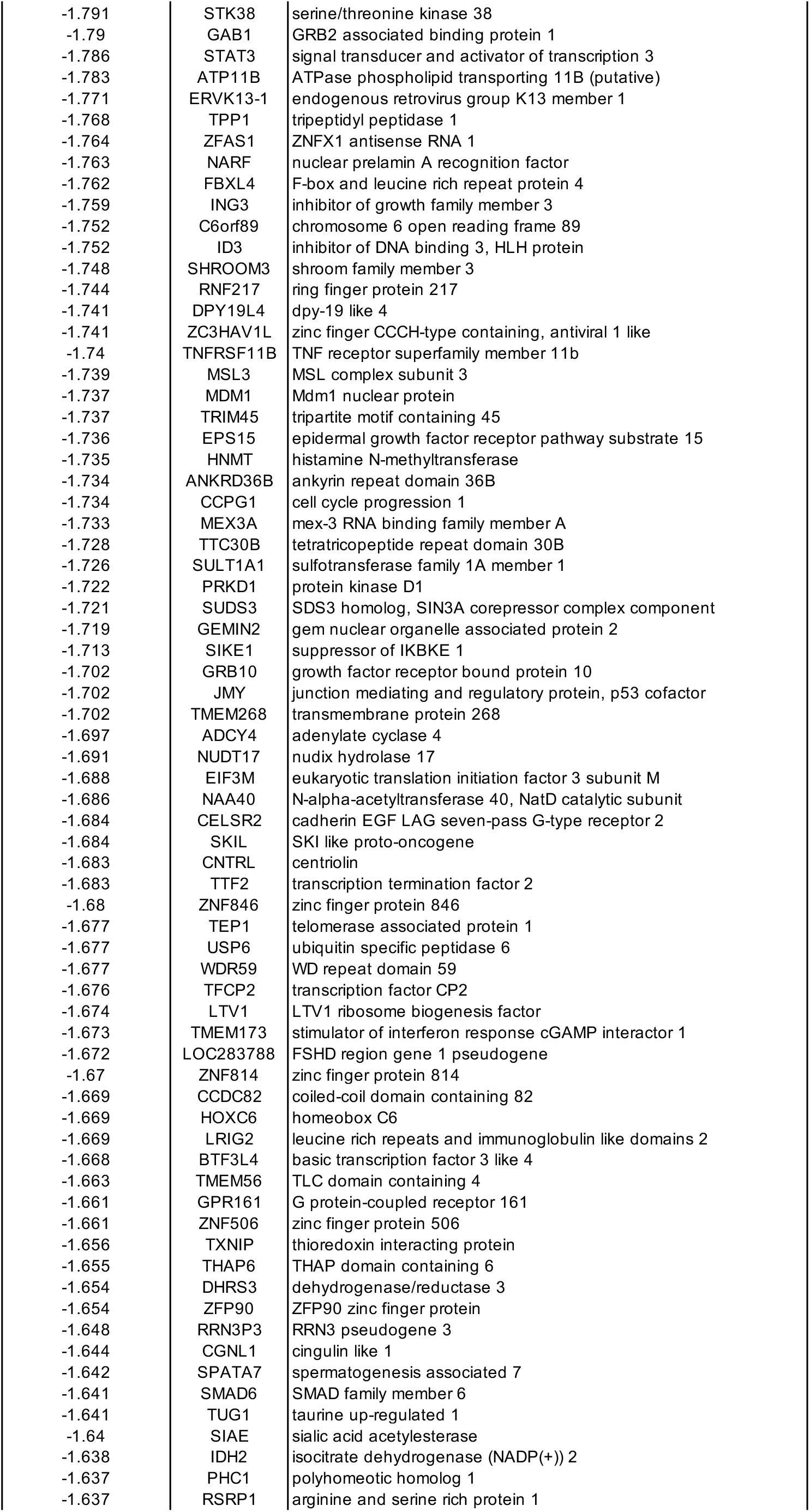
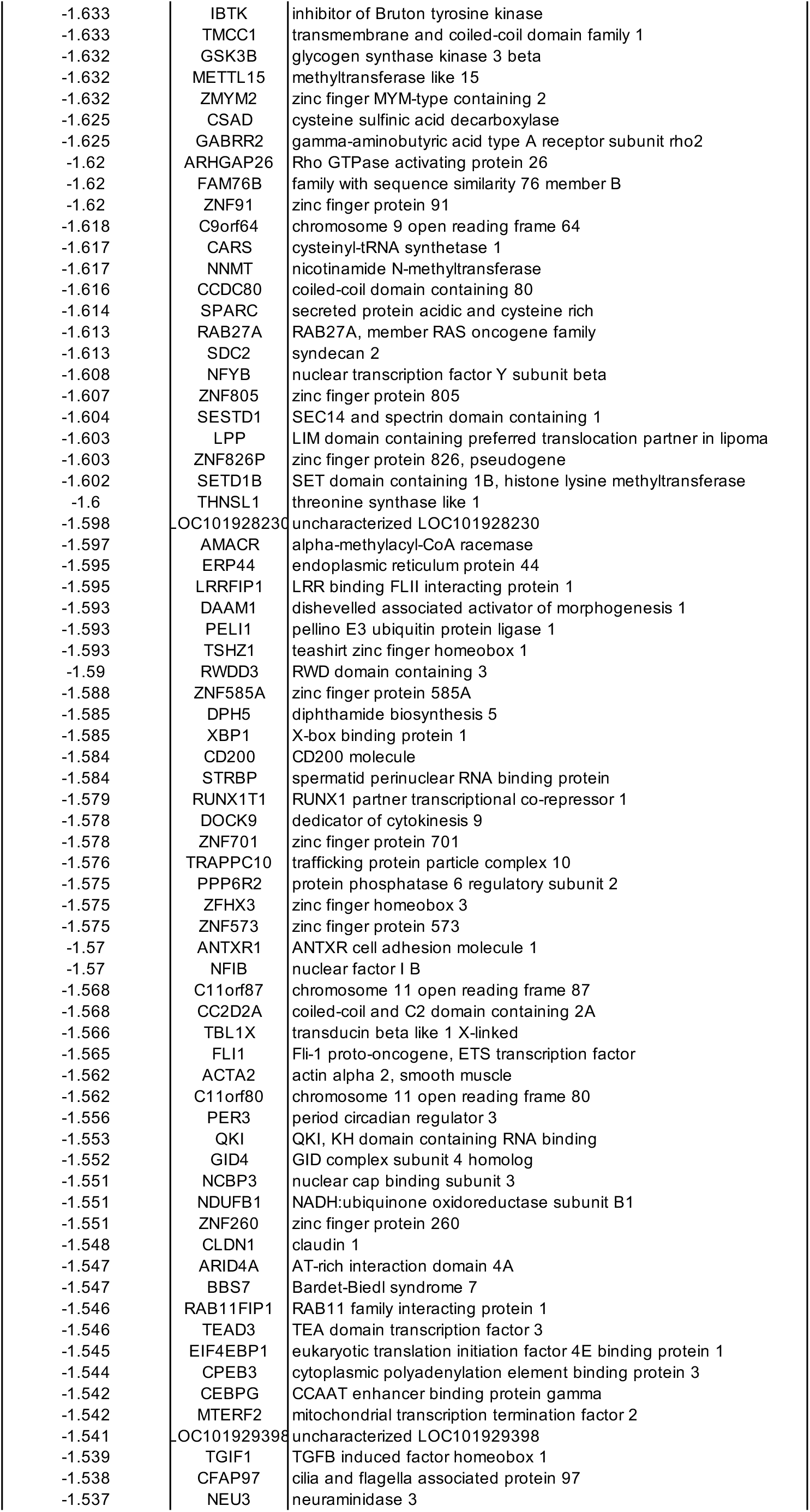
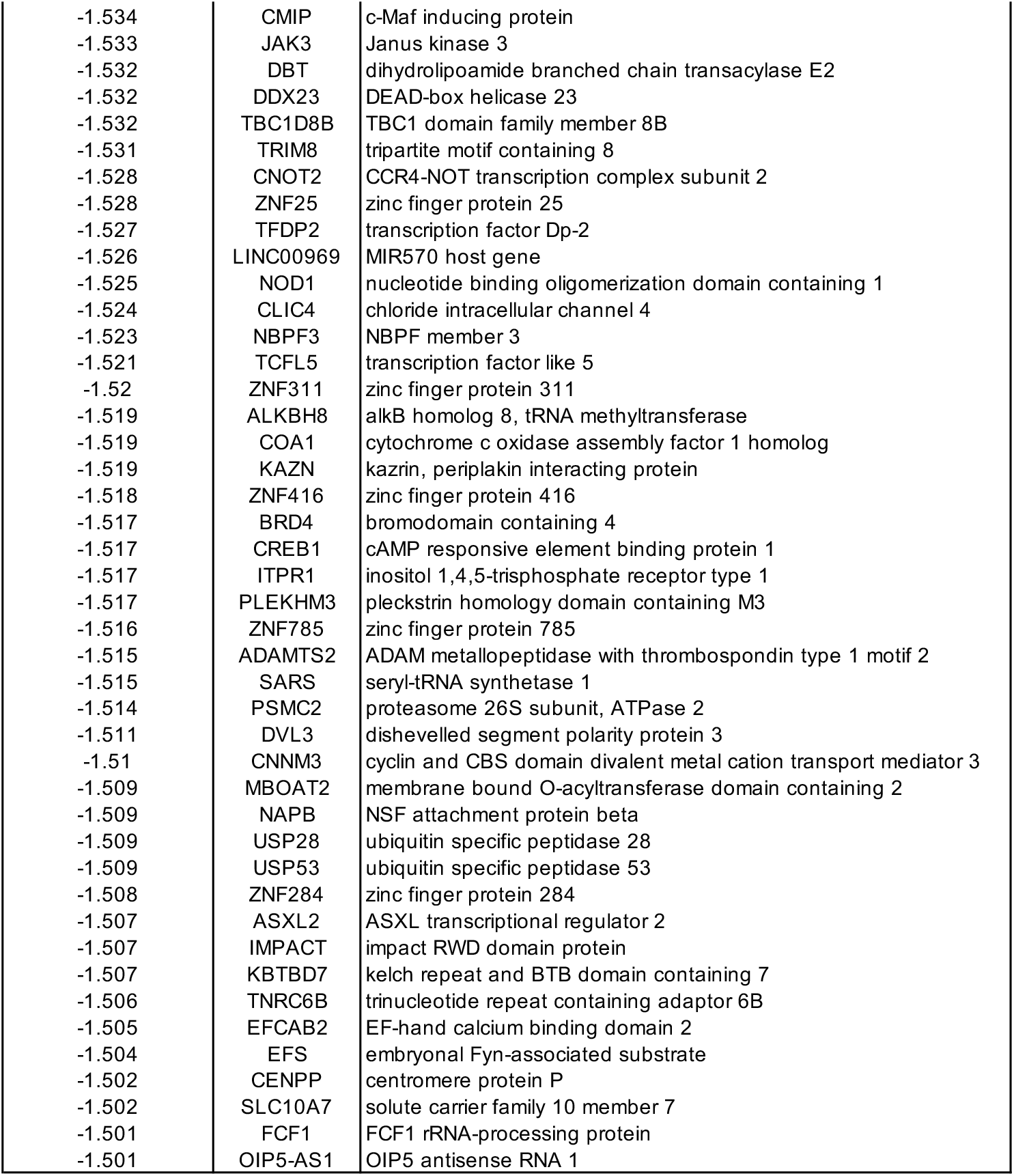
Pt-specifc down-regulated genes (<0.67-fold)

**Table S3.**
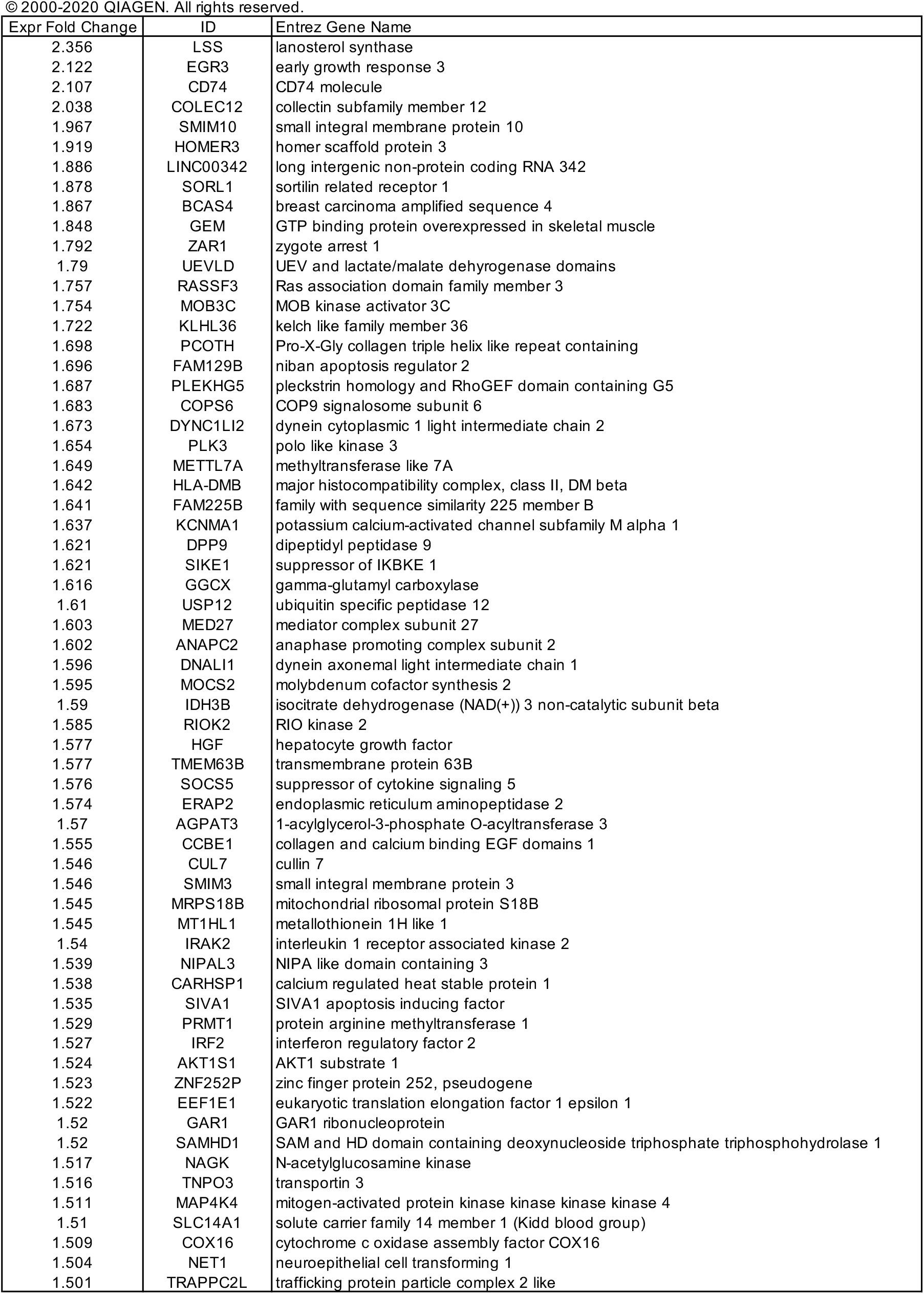
So-specifc up-regulated genes (>1.5-fold)

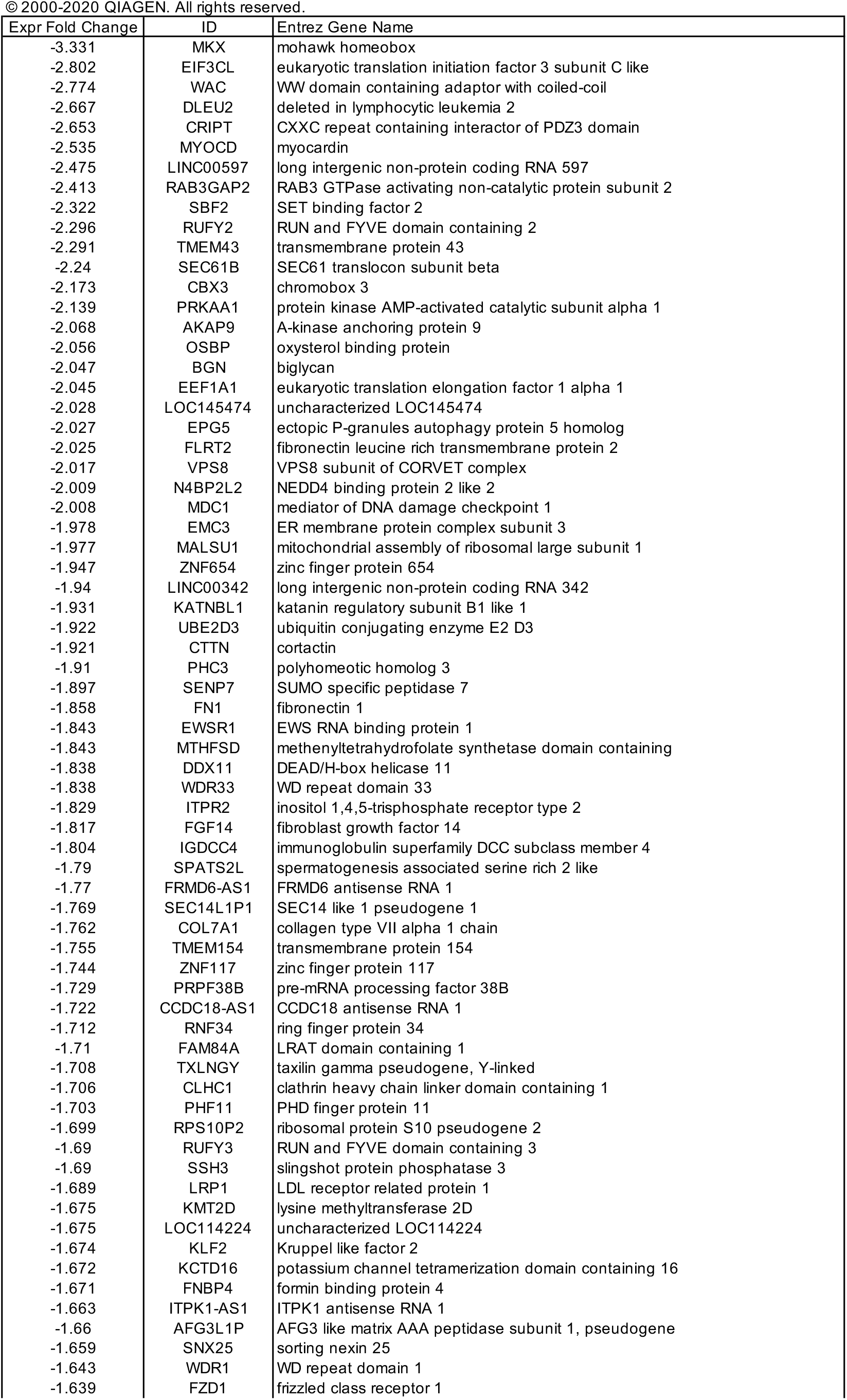
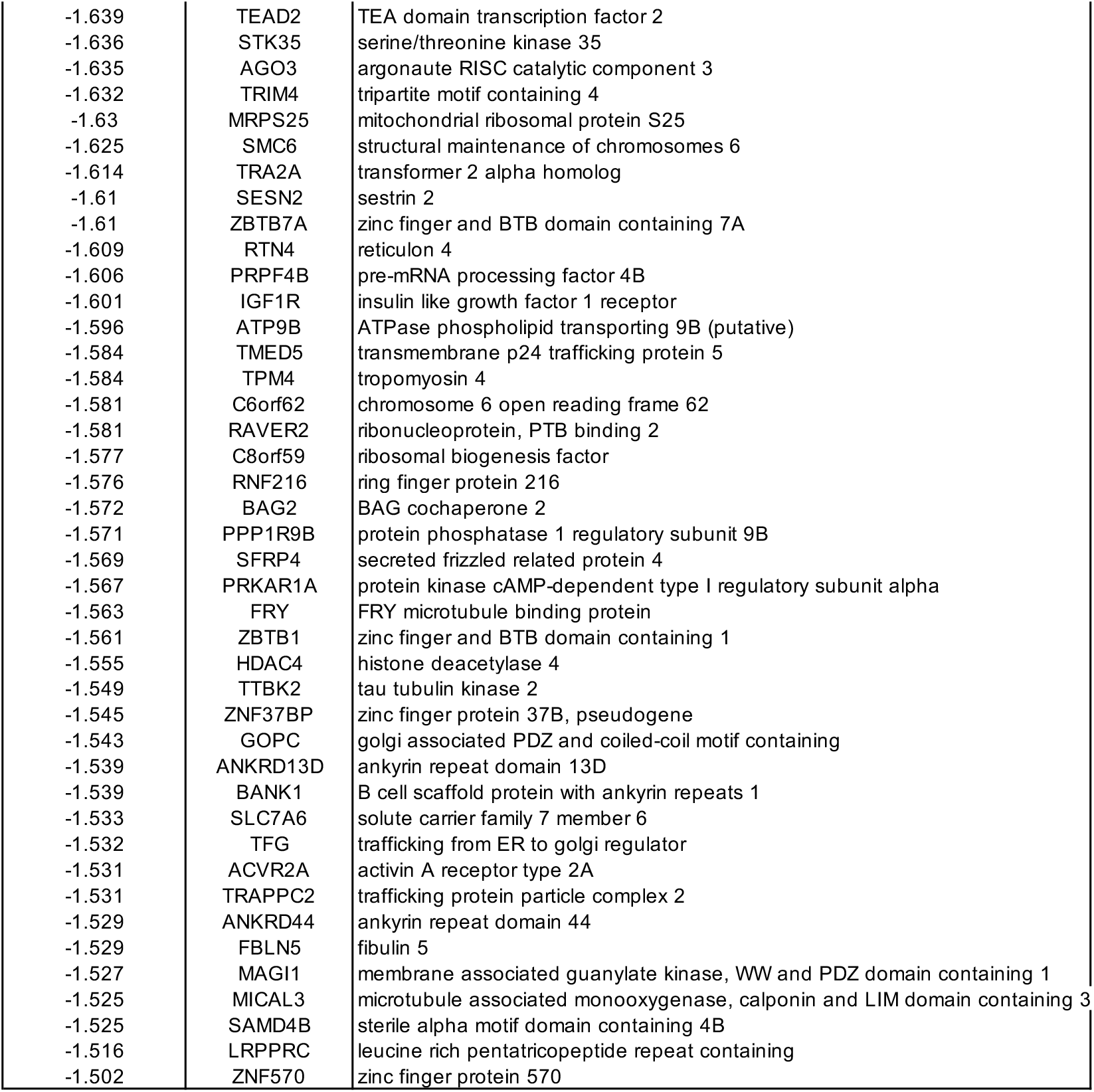
So-specifc down-regulated genes (<0.67-fold)

**Table S4.**
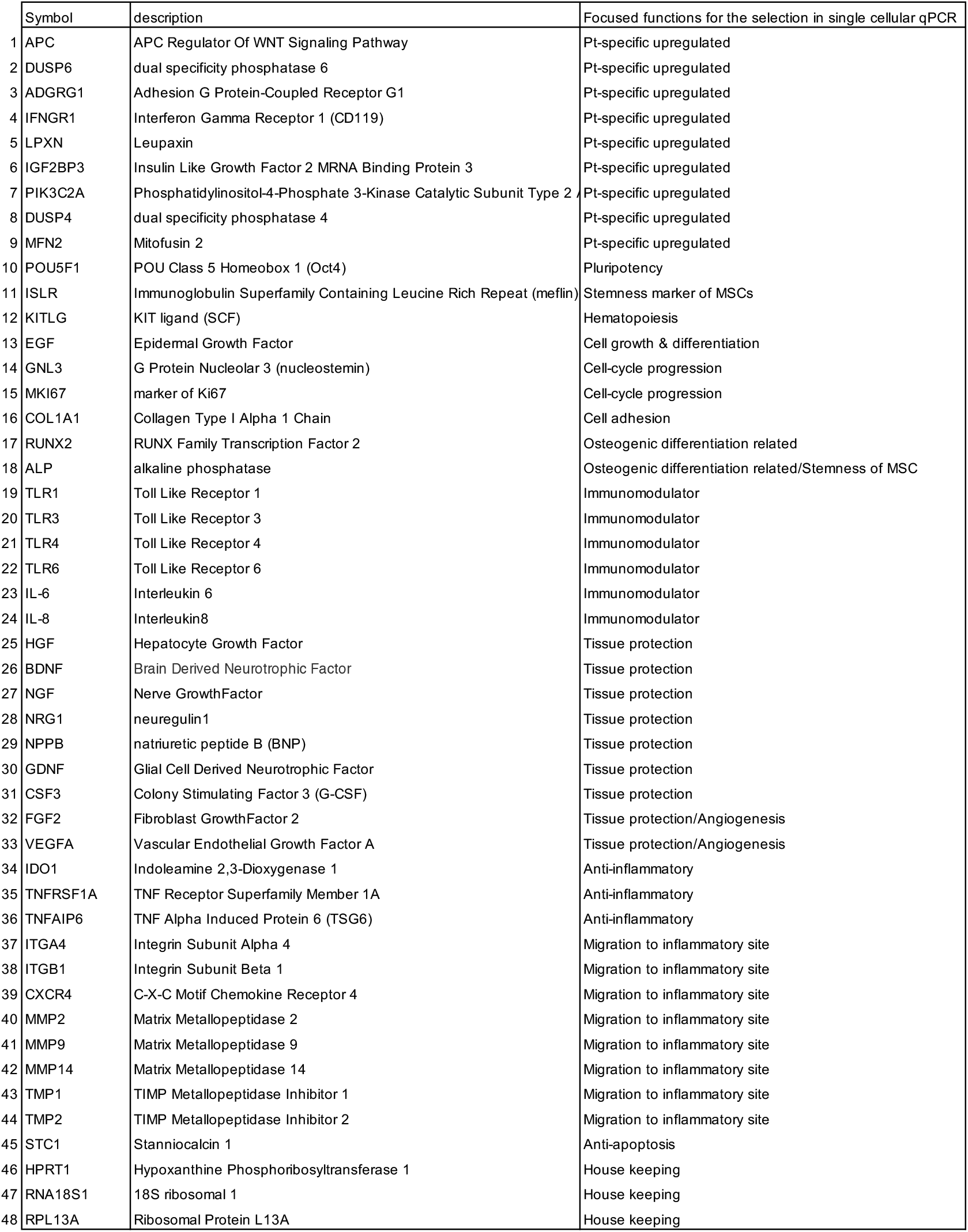
Selected genes for single cellular q-PCR analysis.

